# Discovery of The Clinical Candidate S-892216: A Second-Generation of SARS-CoV-2 3CL Protease Inhibitor for Treating COVID-19

**DOI:** 10.1101/2025.02.26.639208

**Authors:** Yuto Unoh, Keiichiro Hirai, Shota Uehara, Sho Kawashima, Haruaki Nobori, Jun Sato, Hiromitsu Shibayama, Akihiro Hori, Kenji Nakahara, Kana Kurahashi, Masayuki Takamatsu, Shiho Yamamoto, Qianhui Zhang, Miki Tanimura, Reiko Dodo, Yuki Maruyama, Hirofumi Sawa, Ryosuke Watari, Tetsuya Miyano, Teruhisa Kato, Takafumi Sato, Yuki Tachibana

## Abstract

The coronavirus disease 2019 (COVID-19) pandemic crisis has been mitigated by worldwide efforts to develop vaccines and therapeutic drugs. However, there remains concern regarding public health and an unmet need for therapeutic options. Herein, we report the discovery of **S-892216**, a second-generation SARS-CoV-2 3C-like protease (3CL^pro^) inhibitor, to treat COVID-19. **S-892216** is a reversible covalent 3CL^pro^ inhibitor with highly potent antiviral activity and an EC_50_ value of 2.48 nM against SARS-CoV-2 infected cells. Structure-based design of a covalent modifier for compound **1** revealed that introducing a nitrile warhead increased 3CL^pro^ inhibition activity by 180-fold. Subsequent optimization efforts yielded **S-892216**, which combined a favorable pharmacokinetic profile and high off-target selectivity. **S-892216** exhibited antiviral activity against diverse SARS-CoV-2 variants, with no cross-resistance to major mutations reducing antiviral activities of nirmatrelvir and ensitrelvir. In SARS-CoV-2-infected mice, **S-892216** inhibited viral replication in the lungs similar to ensitrelvir, although at a 30-fold lower dose.

## Introduction

The coronavirus disease 2019 (COVID-19) outbreak caused by severe acute respiratory syndrome coronavirus-2 (SARS-CoV-2) has substantially impacted humanity. The development and introduction of vaccines and first-generation antiviral drugs for COVID-19 has enabled us to respond to the acute emergencies. However, the virus is persistently mutating, evading the immune system, and can cause repeated infections, resulting in the disease becoming endemic and not disappearing completely.^1^ Although numerous healthy individuals experience only mild acute symptoms such as cold or seasonal influenza, it remains a potentially fatal illness for high-risk patients.^2^ Additionally, a growing number of patients are experiencing the post-COVID-19 condition (Long COVID), where even mild acute symptoms are followed by various long-term symptoms such as severe chronic fatigue and memory impairment.^3^ Therefore, the need for effective COVID-19 therapeutics remains high as it remains a challenging disease.

The SARS-CoV-2 3C-like protease (3CL^pro^), also known as main protease (M^pro^) and non-structural protein 5 (nsp5), is a promising drug target for COVID-19 therapy. 3CL^pro^ is an essential enzyme for viral growth and is responsible for cleaving polypeptides pp1a/pp1ab.^4–6^ 3CL^pro^ has a characteristic catalytic dyad consisting of C145-H41, where H41 promotes deprotonation of the C145 thiol to generate a highly nucleophilic thiolate that undergoes proteolytic cleavage. Owing to these features, covalent inhibitors targeting C145 have garnered considerable attention in 3CL^pro^ inhibitor research. By attaching a warhead that reacts with the target amino acid residues, covalent inhibitors exert potent activity by forming strong covalent bonds with the target protein.^7–10^ Consequently, several of the developed 3CL^pro^ covalent inhibitors possess a structure that mimics the substrate peptide, incorporating warheads such as aldehydes and nitriles (Figure 1).^11–17^ In general, peptide-mimetic compounds are not optimal for oral drug design owing to their ADME (Absorption, Distribution, Metabolism, and Excretion) profiles, including low membrane permeability and metabolic instability, which complicate the design of suitable candidates. The first-generation SARS-CoV-2 3CL^pro^ inhibitor, nirmatrelvir, developed by Pfizer, requires the co-administration of a pharmacokinetic (PK) booster, ritonavir, to inhibit metabolic enzymes and maintain an adequate blood concentration.^11,18^ Recently, Pfizer discovered PF-07817883 (ibuzatrelvir) as an “upboosted” agent by carefully modifying nirmatrelvir.^19^ To the best of our knowledge, there is limited information on non-peptide covalent inhibitors that exhibit specific and high activity against SARS-CoV-2 3CL^pro^.^20–23^

**Figure 1.**
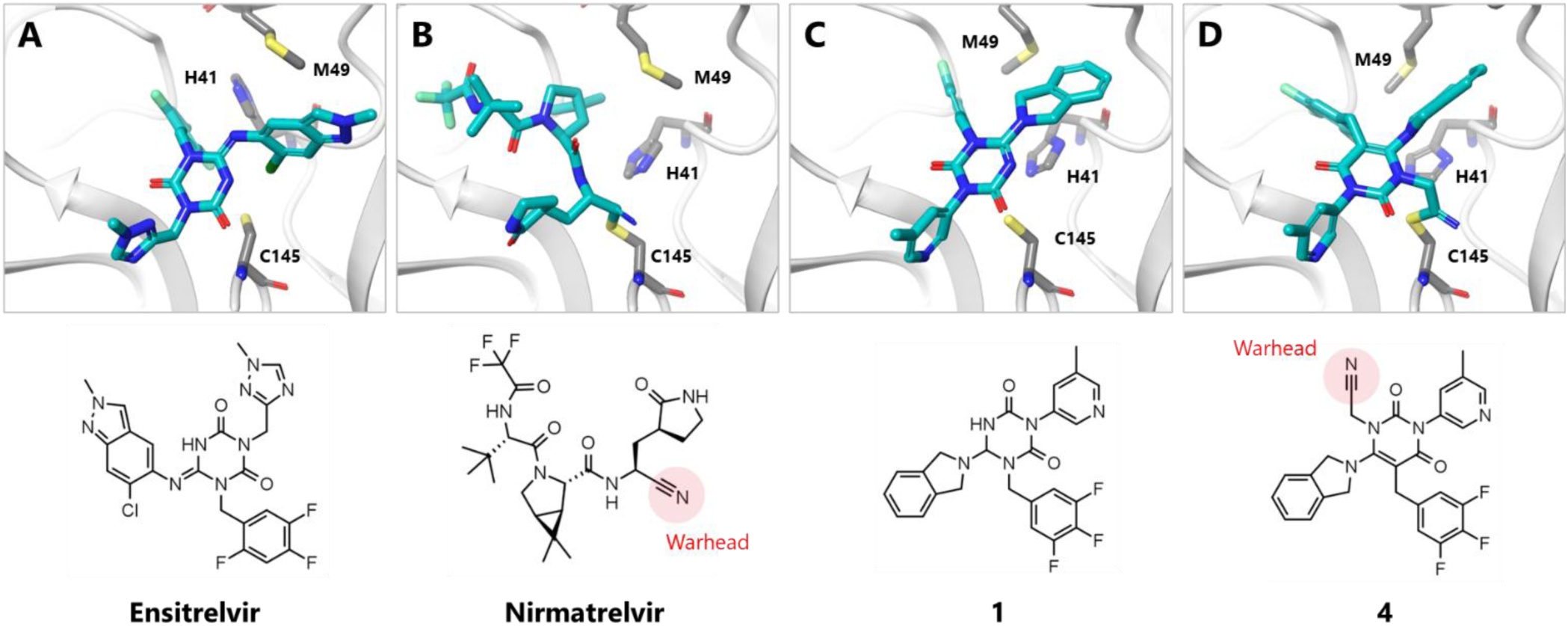
X-ray crystal structures of SARS-CoV2 3CL^pro^ complexed with (A) ensitrelvir (PDB: 7VU6), (B) nirmatrelvir (PDB: 7RFS), (C) compound **1** (PDB: 9LVR) and (D) compound **4** (PDB: 9LVT), respectively. The inhibitors are shown as cyan sticks. The backbone of 3CL^pro^ is shown as a white ribbon, and the three amino acids of H41, M49, and C145 are highlighted as gray sticks. The structures of inhibitors are also depicted at the bottom.

We previously developed a novel non-peptidic and non-covalent novel SARS-CoV-2 3CL^pro^ inhibitor, ensitrelvir (code: **S-217622**), by utilizing *in silico* screening and structure-based drug design.^24^ Ensitrelvir exhibited an excellent PK profile, enabling once-daily oral treatment without a PK booster.^25^ In 2025, ensitrevir was approved for use in Japan and Singapore. Global development of ensitrelvir is ongoing. To meet the therapeutic demands of COVID-19, we have continued research on a second-generation therapeutic drug that exhibits higher activity and superiority to ensitrelvir. In this report, we describe the discovery of a new clinical candidate, **S-892216**, for oral COVID-19 therapy. **S-892216** achieved high antiviral activity against SARS-CoV-2 variants by introducing a reversible covalent warhead to a non-peptidic lead compound while maintaining its excellent PK profile.

## Results and Discussions

To discover compounds that exhibit improved activity while maintaining the excellent ADME profile of ensitrelvir, we endeavored to obtain non-peptide covalent leads by utilizing insights gained during the development of ensitrelvir. As mentioned in previous reports,^24^ ensitrelvir adopts a binding mode distinct from that of other peptide-based inhibitors when interacting with 3CL^pro^ and does not exhibit a clear interaction with the catalytic cysteine of C145 (Figure 1A). Additionally, many other peptidic inhibitors have been found to exhibit high enzymatic affinity by forming a covalent bond with the catalytic cysteine of target proteases, such as nirmatrelvir, which has a nitrile warhead forming a covalent bond with C145 (Figure 1B). Applying a well-designed warhead to a small-molecule inhibitor can enhance its potency substantially. Therefore, we assumed that introducing a covalent interaction with C145 in our compound could further enhance potency. Our attention was directed toward compound **1**, synthesized in a previous study (Table 1). Compound **1** had moderate affinity for SARS-CoV-2 3CL^pro^ (IC_50_=143 nM). However, differences in the three-dimensional conformation between the isoindoline group in **1** and the indazole group in ensitrelvir suggested that they have different binding modes. This intriguing observation prompted us to investigate the binding mode of compound **1**. Figure 1C shows the X-ray crystal structure of the SARS-CoV-2 3CL^pro^ complex with compound **1**. Similar to other inhibitors, compound **1** was bound to the active site of 3CL^pro^, and its core triazinandione moiety interacted with G143 and E166 *via* two carbonyl groups. The 3-methylpyridine group occupied the S1 pocket and formed a key hydrogen bond with H163, whereas the 3,4,5-trifluorobenzene moiety was bound to the S2 pocket, forming lipophilic interactions (Figure S1). The difference between compound **1** and ensitrelvir lies in the conformational changes observed in H41 and M49 side chains. Upon binding ensitrelvir, the side chain of H41 changed its conformation to allow the benzyl group of ensitrelvir to engage in π stacking and induced a binding pocket formation for the 2-chloro-indazole moiety (Figure 1A). In contrast, compound **1** did not impact the H41 conformation but induced an M49 conformational change around the isoindoline group (Figure 1C). The conformation of the 3CL^pro^ active site, especially the catalytic dyad of H41 and C145 bound to compound **1**, was similar to that of other covalent peptide-mimetic inhibitors, such as nirmatrelvir (Figure 1B). H41 is considered a catalytic dyad in 3CL^pro^ and facilitates nucleophilic attack on the substrate *via* deprotonation of the nucleophilic residue C145.^6^ When designing covalent inhibitors, we hypothesized that the docking mode of compound **1** is more suitable than that of ensitrelvir because the conformation of the catalytic dyads in the **1** docking mode is similar to that of the native state, whereas H41 seems to have difficulty accessing C145 in the ensitrelvir docking mode. The crystal structure of compound **1** suggested that C145 was located near the 5-position nitrogen atom of the core triazinandione; therefore, we decided to pursue an approach extending the warhead from the core (Table 1). Because the nitrogen atom on triazinandione could not extend further substituents, we designed compound **2,** which converted the triazinandione core to pyrimidine-dione, eliciting comparable enzymatic activity (IC_50_=335 nM). Exploring substituents on the N atom revealed that compound **3**, which contained a methyl group, did not show any improvement in activity (IC_50_=179 nM). However, introducing a cyanomethyl group, known as a reversible covalent-binding warhead^9^, significantly increased the inhibitory activity of compound **4** by more than 200-fold, reaching an IC_50_ value of 0.760 nM and a SARS-CoV-2 EC_50_ value of 4.94 nM. The X-ray crystal structure of compound **4** complexed with 3CL^pro^ indicated that the cyanomethyl group formed a covalent bond with catalytic cysteine, similar to other covalent 3CL^pro^ inhibitors (Figure 1D). Therefore, we decided to further optimize compound **4** as a starting point to find a second-generation 3CL^pro^ inhibitor.

**Table 1.**
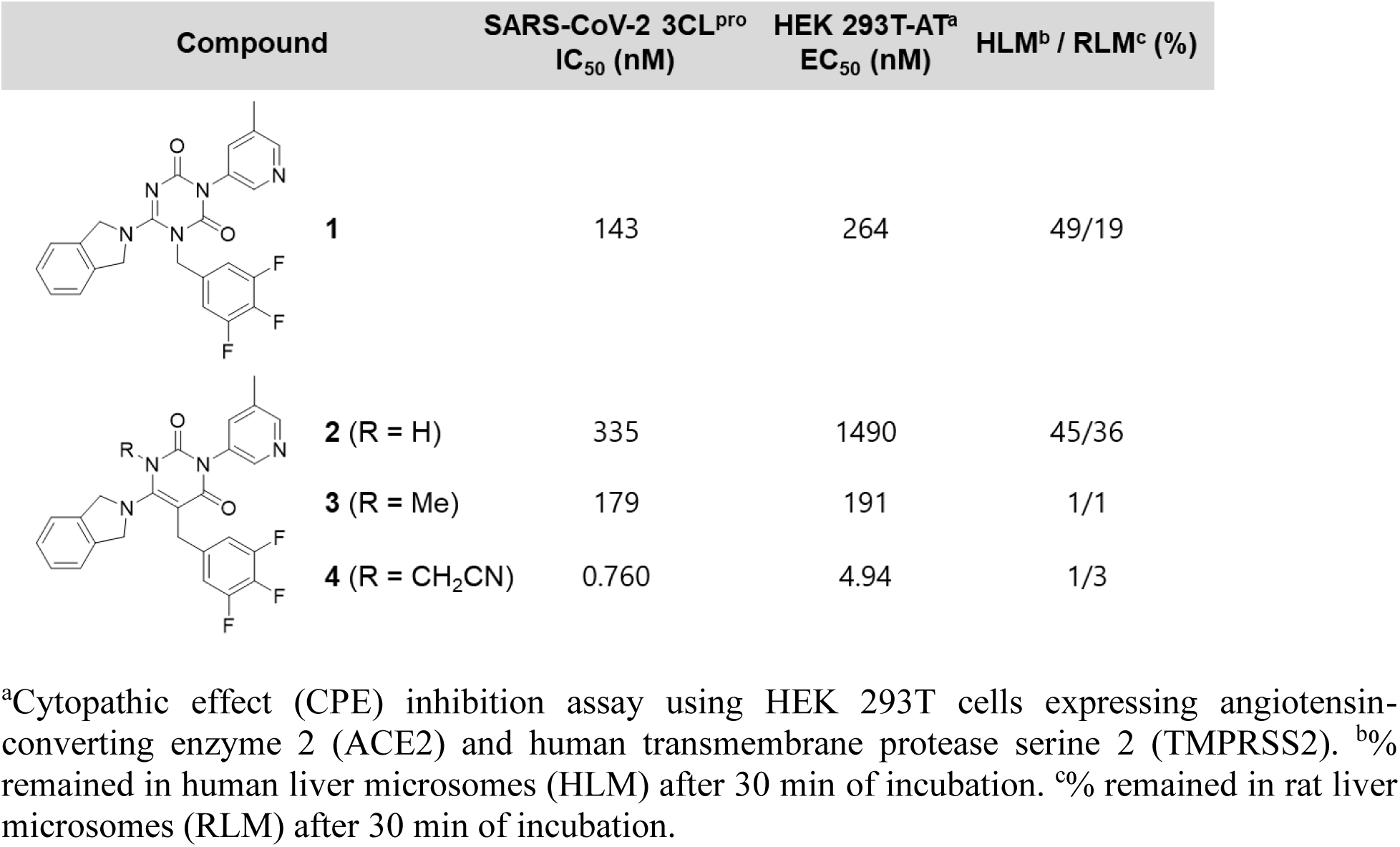
Identification of lead compound 4 as a covalent inhibitor through nitrile-based warhead attachment.

| Compound | SARS-CoV-2 3CL<sup>pro</sup><br>IC<sub>50</sub> (nM) | HEK 293T-AT<sup>a</sup><br>EC<sub>50</sub> (nM) | HLM<sup>b</sup> / RLM<sup>c</sup> (%) |
| --- | --- | --- | --- |
| **1** | 143 | 264 | 49/19 |
| **2** (R = H) | 335 | 1490 | 45/36 |
| **3** (R = Me) | 179 | 191 | 1/1 |
| **4** (R = CH<sub>2</sub>CN) | 0.760 | 4.94 | 1/3 |

<sup>a</sup>Cytopathic effect (CPE) inhibition assay using HEK 293T cells expressing angiotensin-converting enzyme 2 (ACE2) and human transmembrane protease serine 2 (TMPRSS2). <sup>b</sup>% remained in human liver microsomes (HLM) after 30 min of incubation. <sup>c</sup>% remained in rat liver microsomes (RLM) after 30 min of incubation.

Compound **4** exhibited very high activity but was metabolically labile (only 1% remained after incubation in human liver microsomes for 30 min). Therefore, optimization was performed to enhance metabolic stability. Initially, the amine SAR (Structure-Activity-Relationship) at the 6-position of the pyrimidinedione (replacement of isoindoline) was explored (Table 2). Compound **5**, in which the fused isoindoline was replaced with a monocyclic pyrrolidine, showed a >10-fold decrease in activity. However, compound **6**, which has a trifluoromethyl-substituted piperidine moiety, demonstrated improved activity comparable to that of compound **4**. Notably, filling the S1’ pocket with the lipophilic substituents was crucial for activity. Compounds **4** and **5** did not exhibit improved metabolic stability; however, a series of spiroazetidine derivatives of compounds **7**-**9** showed better metabolic stability and slightly reduced activity. Compound **9**, possessing 6-oxa-2-azaspiro[3.4]octane, showed a balanced profile with an IC_50_ value of 8.37 nM and metabolic stability in human liver microsomes (58% remaining at 30 min). Therefore, we decided to attach 6-oxa-2-azaspiro[3.4]octane onto the amine side chain and optimize other parts.

**Table 2.**
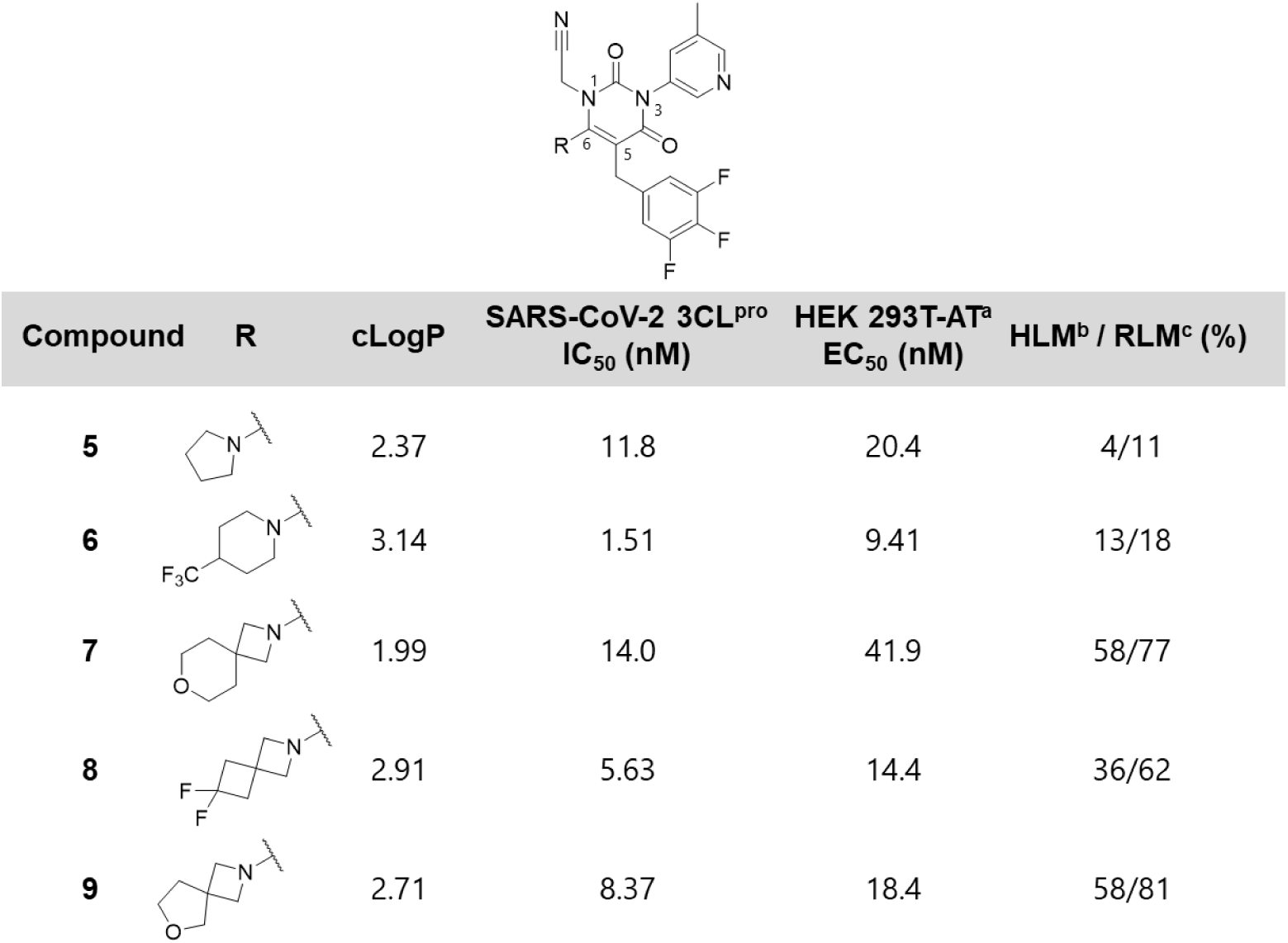
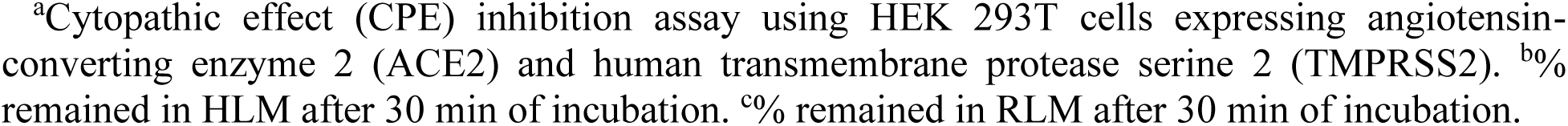
Pyrimidine-dione 6-position amine SAR.

| Compound | R | cLogP | SARS-CoV-2 3CL<sup>pro</sup><br>IC<sub>50</sub> (nM) | HEK 293T-AT<sup>a</sup><br>EC<sub>50</sub> (nM) | HLM<sup>b</sup> / RLM<sup>c</sup> (%) |
| --- | --- | --- | --- | --- | --- |
| **5** |  | 2.37 | 11.8 | 20.4 | 4/11 |
| **6** |  | 3.14 | 1.51 | 9.41 | 13/18 |
| **7** |  | 1.99 | 14.0 | 41.9 | 58/77 |
| **8** |  | 2.91 | 5.63 | 14.4 | 36/62 |
| **9** |  | 2.71 | 8.37 | 18.4 | 58/81 |

<sup>a</sup>Cytopathic effect (CPE) inhibition assay using HEK 293T cells expressing angiotensin-converting enzyme 2 (ACE2) and human transmembrane protease serine 2 (TMPRSS2). <sup>b</sup>% remained in HLM after 30 min of incubation. <sup>c</sup>% remained in RLM after 30 min of incubation.

A detailed optimization study is presented in Table 3. When the 3-methyl group of pyridine, considered one of the metabolized points, was converted to a chlorine group (compound **10**), the inhibitory activity was almost maintained. However, the metabolic stability of compound **10** deteriorated, probably due to its increased lipophilicity (ClogP=3.08). Reducing the fluorine atoms from the benzyl group occupying the S2 pocket (compounds **11** and **12**) decreased lipophilicity (ClogP=2.69 and 2.64) and improved metabolic stability but resulted in lower activities. Subsequently, we found compound **13**, in which the S2 benzyl group was replaced by a phenyl group, balancing activity and metabolic stability, even without fluorine atoms. Reportedly, lipophilic interaction in the S2 pocket is critical for the inhibitory activity of SARS-CoV-2 3CL^pro^ inhibitors.^26^ Therefore, we designed 3-chloro and 4-fluoro substituents on the S2 phenyl group (compounds **14** and **15**), which improved activity by more than 10-fold. Thereafter, we continued exploring the potential of amine substituents at the 6-position to further optimize the metabolic stability of compound **15**. The small monocyclic difluoroazetidine-substituted compound **16** showed good metabolic stability but slightly decreased activity compared with the spiro types. We then carefully examined combinations of substituents and discovered compound **17**, which achieved high activity against 3CL^pro^ (IC_50_=0.655 nM) and SARS-CoV-2 infected cells (EC_50_=2.48 nM) while maintaining excellent metabolic stability. Furthermore, exploring the azetidine substituents led to the discovery of compound **18**, which had a tertiary alcohol and reduced lipophilicity (ClogP=2.48). With the promising compounds **17** and **18** which exhibited good potency and liver microsomal stability in hands, we next examined for their PK profiles.

**Table 3.** Lead optimization.

| compound | R<sup>1</sup> | R<sup>2</sup> | R<sup>3</sup> | cLogP | SARS-CoV-2 3CL<sup>pro</sup><br>IC<sub>50</sub> (nM) | HEK 293T-AT<sup>a</sup><br>EC<sub>50</sub> (nM) | HLM<sup>b</sup> / RLM<sup>c</sup> (%) |
| --- | --- | --- | --- | --- | --- | --- | --- |
| 10 |  |  | Cl | 3.08 | 8.27 | 25.3 | 17 / 54 |
| 11 |  |  | Me | 2.69 | 19.6 | 41.8 | 55 / 71 |
| 12 |  |  | Me | 2.64 | 99.6 | 117 | 70 / 70 |
| 13 |  |  | Cl | 2.66 | 7.16 | 23.1 | 55 / 58 |
| 14 |  |  | Cl | 3.14 | 0.403 | 1.57 | 49 / 33 |
| 15 |  |  | Cl | 3.21 | 0.493 | 4.48 | 55 / 58 |
| 16 |  |  | Cl | 2.49 | 2.36 | 7.79 | 89 / 83 |
| 17<br>(S-892216) |  |  | Cl | 3.21 | 0.655 | 2.48 | 99 / 92 |
| 18 |  |  | Cl | 2.48 | 0.391 | 3.82 | 103 / 99 |

<sup>a</sup>Cytopathic effect (CPE) inhibition assay using HEK 293T cells expressing angiotensin-converting enzyme 2 (ACE2) and human transmembrane protease serine 2 (TMPRSS2). <sup>b</sup>% remained in HLM after 30 min of incubation. <sup>c</sup>% remained in RLM after 30 min of incubation.

*In vitro* metabolic stability assays revealed that compounds **17** and **18** were retained in the hepatocytes of humans, rats, dogs, and monkeys for a prolonged duration (Table 4). As for *in vivo* PK studies, both compounds exhibited similar and sufficient half-lives (t_1/2_) and total clearance (CL_tot_) after a single intravenous administration across species, especially compound **17**, which demonstrated a prolonged t_1/2_ of 24.9 h in dogs. However, the rat oral availability *(F*) of the two compounds showed different values: *F* of compound **18** was 38%, whereas that of compound **17** reached 100%. The low-to-moderate oral availability of **18** could be explained by membrane permeability. *In vitro* permeability assay using the LLC-PK1 cell line revealed that *P_app_* values of **17** and **18** were 6.17 × 10^−6^ cm/sec and 1.26 × 10^−6^ cm/s, respectively. We assumed that the relatively low permeability of **18** could be attributed to an additional hydrogen donor on the azetizine moiety. For both compounds, the serum fraction of unbound (*f_u_*) was in a similar range at approximately 10% for all species. Based on a comprehensive PK study, we selected compound **17** (development code **S-892216**) as a potential clinical candidate for further evaluation.

**Table 4.**
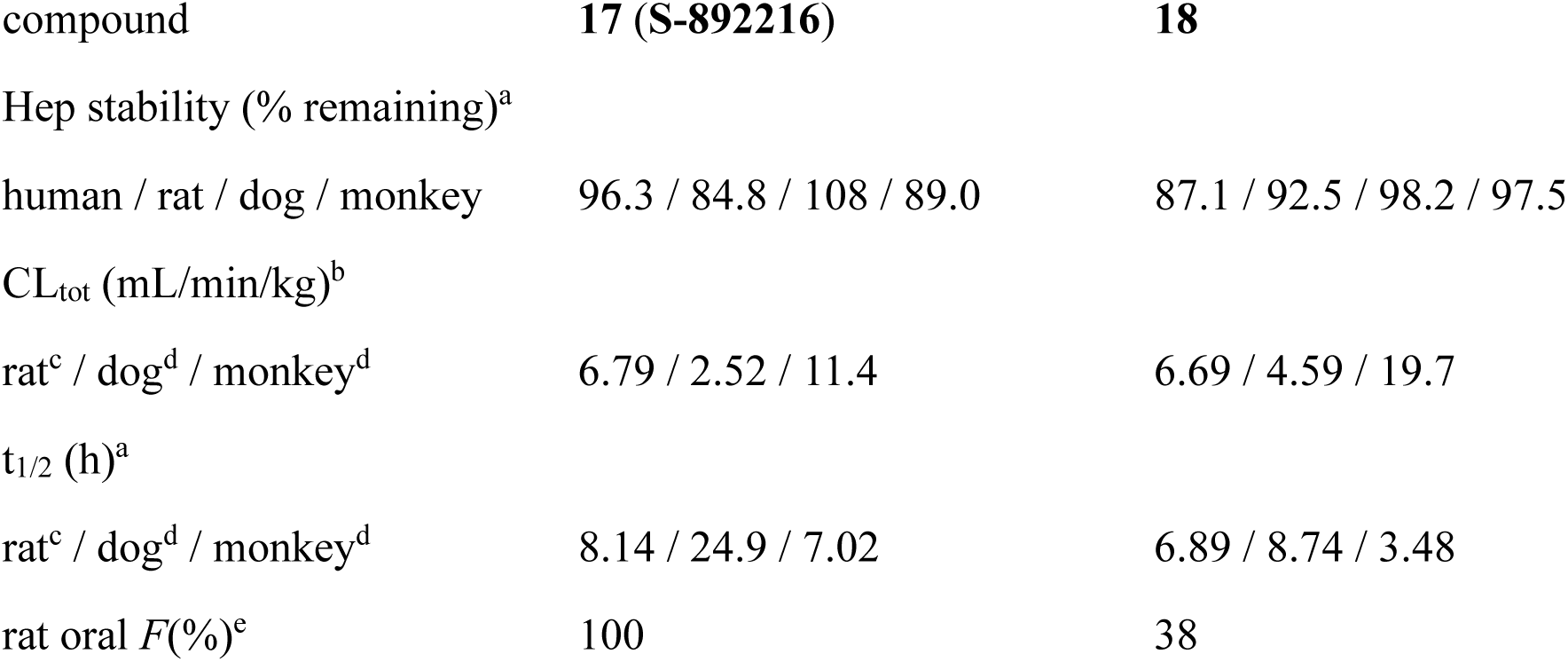
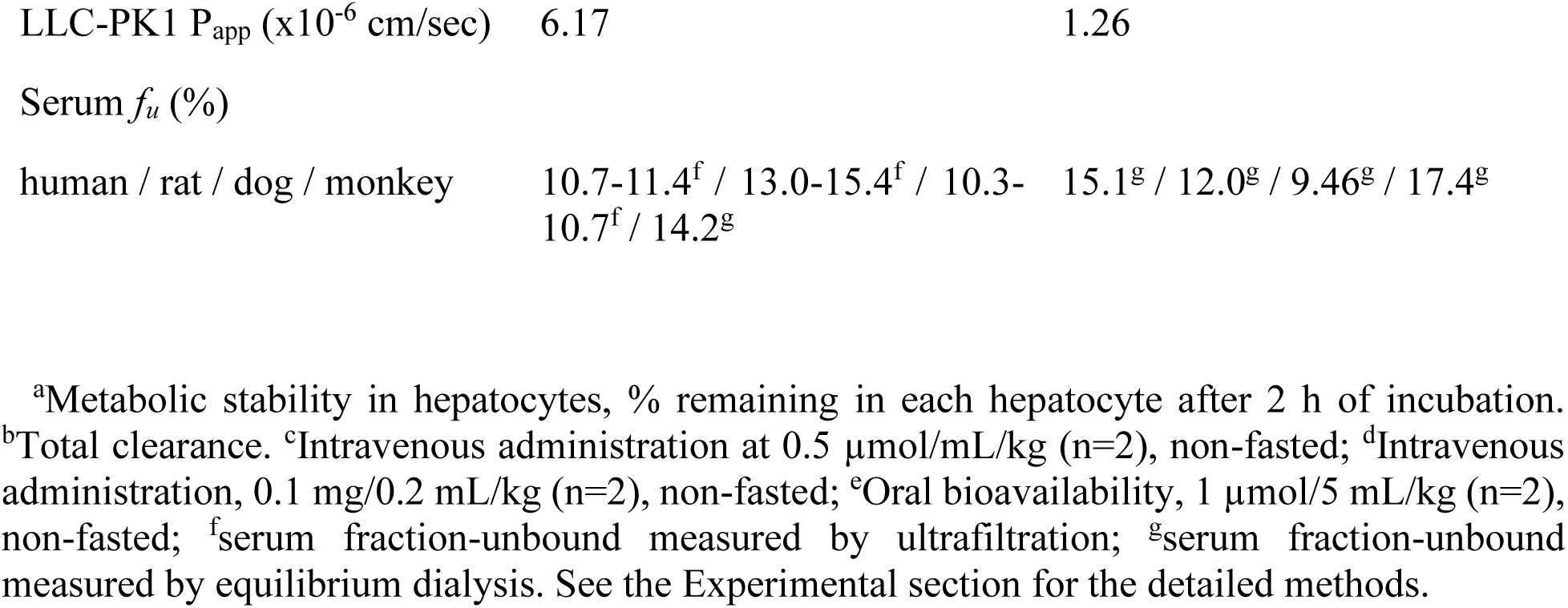
Summary of the pharmacokinetic profiles of compounds 17 (S-892216) and 18.

Considering the binding mode of **S-892216**, the X-ray crystal structure revealed covalent binding of **S-892216** with the catalytic cysteine C145 of SARS-CoV-2 3CL^pro^ (Figure 2). In addition, **S-892216** formed four critical hydrogen bonds and lipophilic interaction in S2 and S1’ pockets. Covalent bond formation of the nitrile warhead produced an imine that formed a hydrogen bond with the main chain of C145, and the two carbonyl oxygen atoms of the pyrimidine-dione core formed hydrogen bonds with the main chains of E166 and G143. At the S1 site, the 3-chloro pyridine unit formed a hydrogen bond with the side chain of H163, similar to other 3CL^pro^ inhibitors (see Figure 1). The 3-chloro-4-fluorobenzene moiety filled the S2 pocket and formed a π stacking with the catalytic histidine of H41, with the 6,6-difluoro-2-azaspiro[3.3]heptane lying on the lipophilic S1’ pocket.

**Figure 2.**
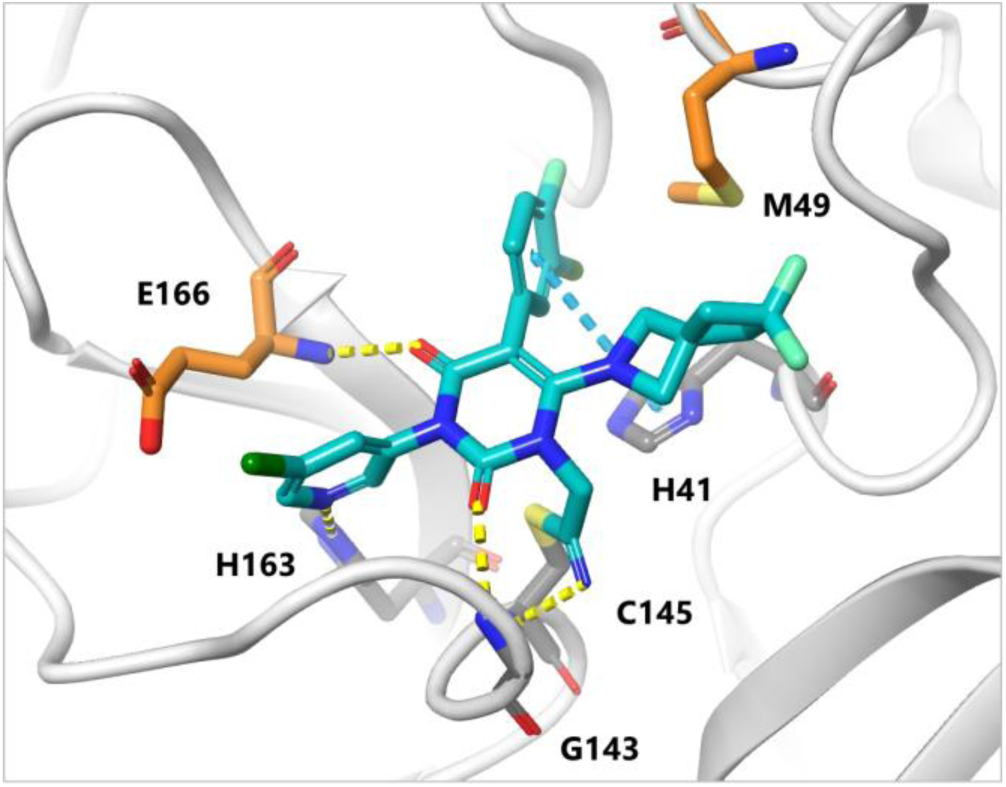
X-ray crystal structure of **S-892216** complexed with SARS-CoV-2 3CL^pro^ (PDB: 9LVV). **S-892216** is shown as a cyan stick, and the protein backbone is shown as a gray ribbon. Residues interacting with **S-892216** are indicated as gray and orange sticks. Hydrogen bonds are indicated as yellow dashed lines, and π–π stacking is indicated as a cyan dashed line.

To characterize the covalent-binding properties of **S-892216**, its reversible property was examined. Reversible covalent inhibitors are considered to have an advantage over irreversible covalent binders in terms of their off-target selectivity.^10, 27^ It is known that nitrile is a reversible covalent warhead adducted to cysteine and serine residues.^9^ However, the slow dissociation of **S-892216** impairs investigations into its reversibility and binding kinetics using surface plasmon resonance (SPR) or other related methods. Therefore, we developed a scintillation proximity assay (SPA)-based competitive experiment using ^14^C isotope-labeled **S-892216** and confirmed its dissociation kinetics (Figure 3). The binding of an isotope-labeled compound to an enzyme can be determined from the scintillation signal. In the case of incubation of [^14^C]-**S-892216** alone with 3CL^pro^, the signal slightly decreased, although no obvious dissociation was observed after 1500 min (Sample 1). In contrast, when a 30-fold higher concentration of non-labeled **S-892216** was added to the pre-incubated [^14^C]-**S-892216** and 3CL^pro^, the complexed [^14^C]-**S-892216** was replaced with a non-labeled one, resulting in gradual down-signaling (Sample 2). Once the isotope-labeled [^14^C]-**S-892216** left 3CL^pro^, it was unable to bind to the enzyme because of competition from the large amount of non-labeled **S-892216**, with the signal eventually reduced to the same level as that at baseline (Sample 3: [^14^C]-**S-892216** and excess non-labeled **S-892216** were added before preincubation). The results indicate that **S-892216** is a reversible covalent inhibitor with an estimated k_off(obs)_ value of 4.06×10^−5^ s^−1^.

**Figure 3.**
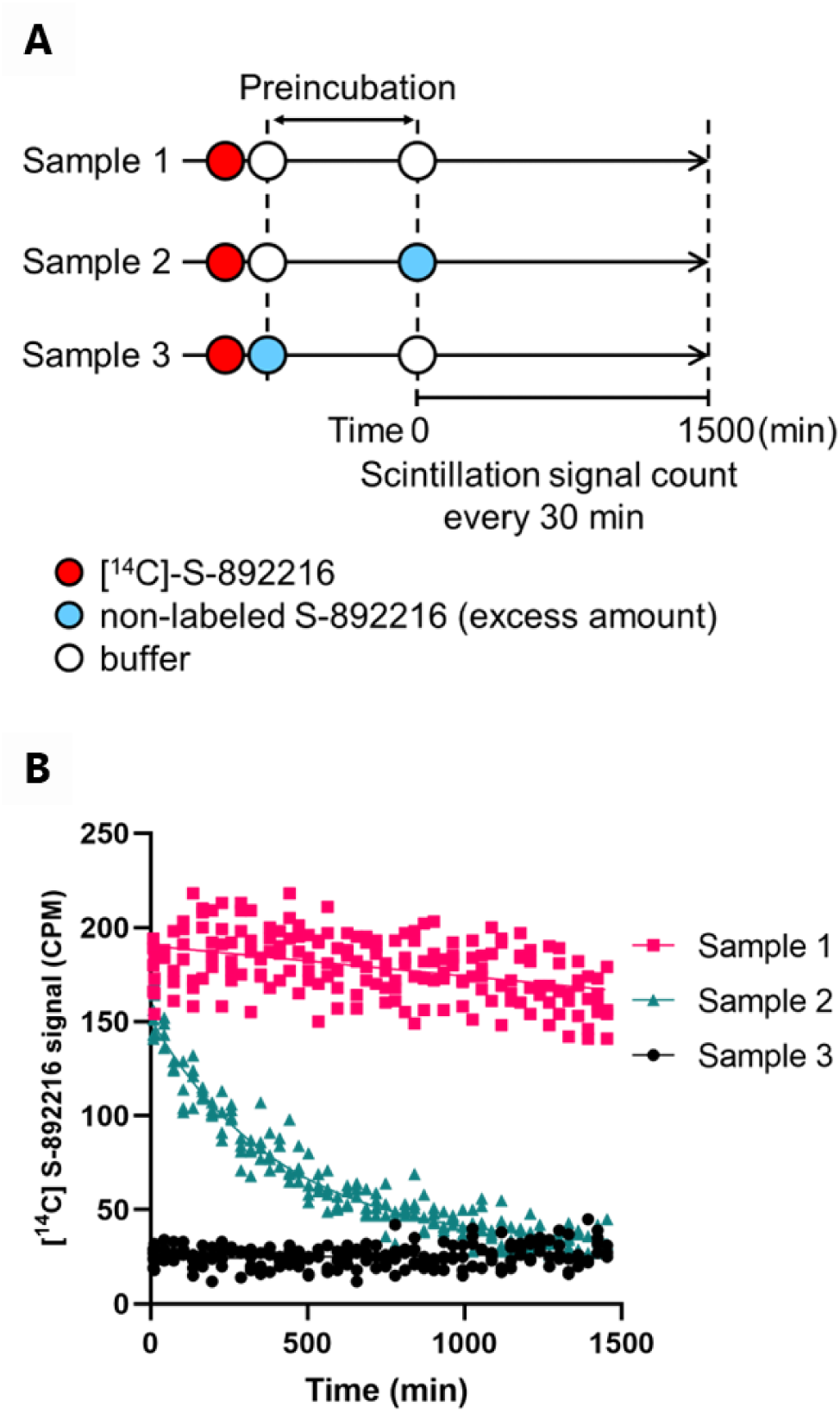
Dissociation kinetics of **S-892216** from 3CL^pro^ measured using a scintillation proximity assay (SPA)-based assay. (A) Schematic illustration of **S-892216** dissociation kinetics study. Sample 1: [^14^C]-**S-892216** alone; Sample 2: Excess amount of non-labeled **S-892216** was added immediately before measurement; Sample 3 Excess amount of non-labeled **S-892216** was added before preincubation. [^14^C]-**S-892216**: red circle, Non-labeled **S-892216**: blue circle, Buffer: white circle. (B) Dissociation kinetics of [^14^C]-**S-892216** was monitored using scintillation signals. Data fitted in GraphPad Prism 9 using one-phase exponential decay.

The off-target selectivity of **S-892216** was examined using *in vitro* enzymatic assays with various human-related proteases and HIV protease (Table S1). **S-892216** lacked inhibitory activity against caspase-2, chymotrypsin, cathepsin B/D/G/L, thrombin, or HIV-1 protease at concentrations up to 10,000 nM, suggesting high target selectivity for SARS-CoV-2 3CL^pro^. Next, we characterized the various cell-based antiviral activities of **S-892216**. **S-892216** exhibited potent antiviral activities against various SARS-CoV-2 variants, including the Omicron JN.1 strain, with EC_50_ values of 2.27 to 12.5 nM in VeroE6/TMPRSS2 cells (Figure 4a and Table S2) without cytotoxicity (CC_50_ > 100,000 nM). Given the multitude of variations in the Spike proteins of circulating variant strains, antibody drugs targeting this protein have lost their efficacy.^28,29^ Conversely, 3CL^pro^, particularly near its active site, exhibits a low propensity for naturally occurring mutations. The 3CL^pro^ residues that directly contact or are located within 5Å of **S-892216** are highly conserved (Table S3), with a frequency of ≥ 99.9%, in the Global Initiative on Sharing Avian Influenza Data (GISAID; https://gisaid.org/) database as of December 31, 2024. Hence, if this trend is maintained, 3CL^pro^ inhibitors such as **S-892216** could retain their effectiveness against the variant strains. Additionally, **S-892216** displayed anti-SARS-CoV-2 activity in VeroE6/TMPRSS2 cells with P-glycoprotein (P-gp) inhibitor and A549-Dual hACE2-TMPRSS2 cells, a human cell line derived from lung cancer overexpressing ACE2 and TMPRSS2, with EC_50_ values of 3.36 and 2.21 nM, respectively (Table S4). Moreover, in human airway epithelial cells (hAECs), **S-892216** demonstrated antiviral activity against the Omicron variants, with EC_90_ values of 2.31 to 2.41 nM (Figure 4b). Notably, **S-892216** exhibited superior anti-SARS-CoV-2 activity compared to the other compounds, including nirmatrelvir and ensitrelvir.

**Figure 4.**
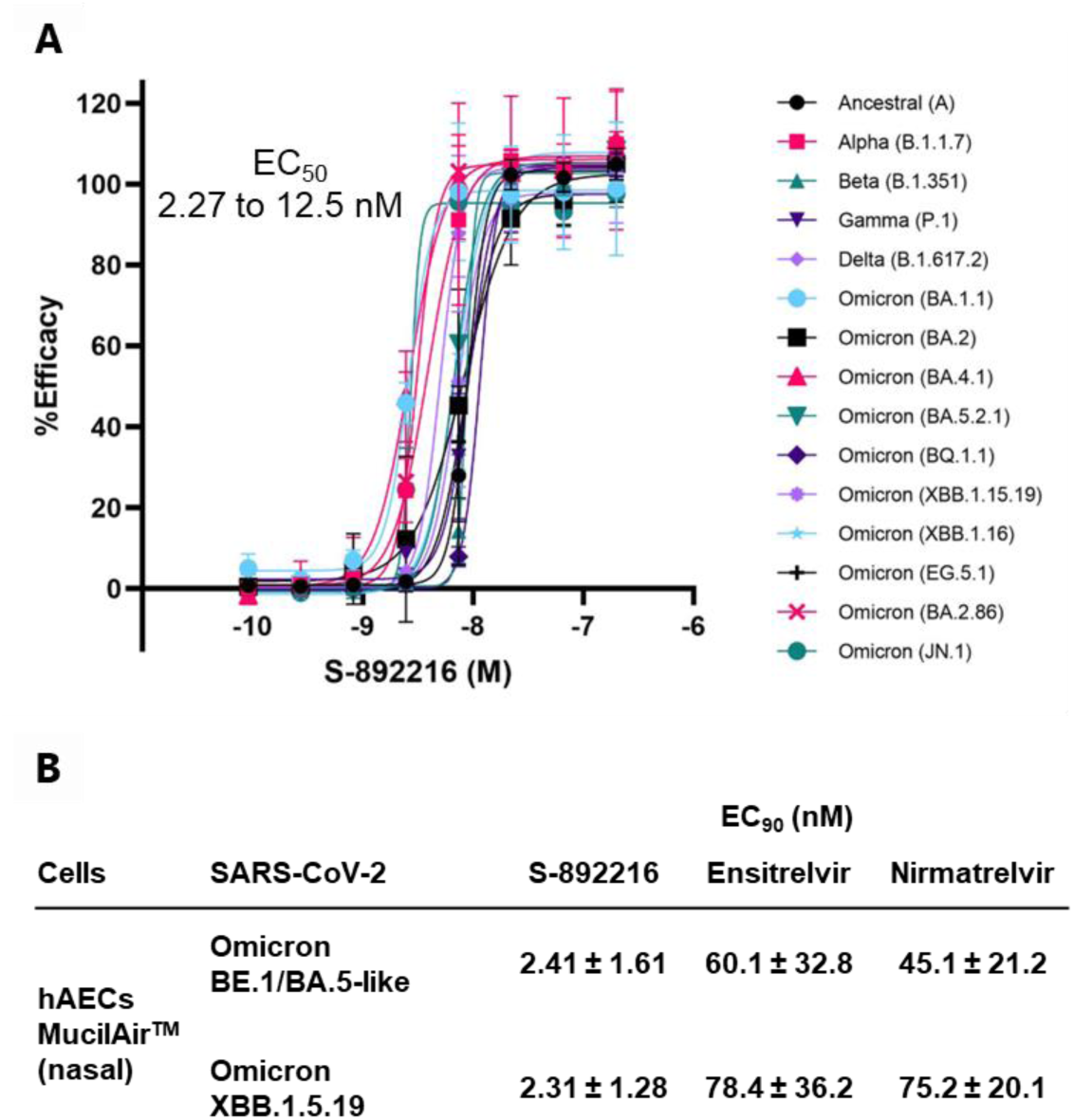
*In vitro* antiviral activity of **S-892216**. (a) Antiviral activity of **S-892216** against various SARS-CoV-2 strains in a cytopathic effect (CPE) inhibition assay using VeroE6/TMPRSS2 cells. (b) Antiviral activity of **S-892216** against Omicron strains in suppressing virus production using human airway epithelial cells (hAECs). Data are the means ± SD; n=3 biological replicates.

The effects of human and mouse serum on the *in vitro* anti-SARS-CoV-2 activity of **S-892216** were examined. A potency shift in anti-SARS-CoV-2 activity was observed due to the addition of serum, with potency shifts of 4.78- and 11.7-fold, respectively, in the presence of 100% human and mouse serum, respectively (Table S5). Human serum *f_u_* and mouse serum *f_u_* were 10.7-11.4% and 5.2-6.2%, respectively. The serum *f_u_* in humans was approximately two times higher than in mice. Consequently, potency shifts were found to correlate with the serum *f_u_*.

Furthermore, **S-892216** exhibited promising antiviral activity against other beta-coronaviruses, including SARS-CoV, Middle East respiratory syndrome coronavirus (MERS-CoV), and human coronavirus OC43 (HCoV-OC43), with EC_50_ values of 6.61, 57.3, and 92.0 nM, respectively (Table S6).

Given that **S-892216** targets 3CL^pro^ similar to nirmatrelvir and ensitrelvir, assessing the cross-resistance profile is also important. The antiviral efficacy of **S-892216** against nirmatrelvir- and ensitrelvir-reduced susceptibility mutants was evaluated using reverse genetics-derived SARS-CoV-2 (rgSARS-CoV-2). In particular, E166V and M49L in 3CL^pro^ have been reported to exhibit reduced susceptibility to nirmatrelvir and ensitrelvir.^30,31^ Antiviral activity of **S-892216** was not significantly influenced by amino acid substitutions, including the E166V and M49L mutations with fold change (FC) values <0.7 (Table S7). These small fold changes against the 3CL^pro^ mutants could be explained by the binding mode of **S-892216**. It is reported that E166V mutation reduces susceptibility to nirmatrelvir owing to the loss of a critical hydrogen bond between the sidechain of E166 and nirmatrelvir.^32^ In contrast, **S-892216** did not interact with the E166 side chain (Figure 2), resulting in similar activity toward the E166V mutant and the wild type. Similarly, the M49L mutants that showed reduced susceptibility to ensitrelvir did not affect **S-892216** activity because the M49L mutation did not significantly influence the hydrophobic interaction between M49 and **S-892216** (Figure 2).

To obtain information regarding viruses with reduced susceptibility to **S-892216**, *in vitro* selection of **S-892216** resistance mutation was conducted (Figure S2A). VeroE6/TMPRSS2 cells infected with the SARS-CoV-2 Omicron BE.1/BA.5-like variant (hCoV-19/Japan/TY41-702/2022) were cultured in the presence of **S-892216**. The initial passage concentrations of **S-892216** were 5.56, 16.7, and 50.0 nM. All SARS-CoV-2 cells cultured with **S-892216** showed CPE from passages 1 to 10. Because the viral stocks from samples maintained at 50 nM **S-892216** lacked sufficient viral titers, sequencing and drug susceptibility testing of these stocks were discontinued. Genotypic analysis revealed amino acid substitutions of P252L, M49K/P252L, L50F/P252L, D48E/L50F/P252L, and M49K/N221K/P252L in 3CL^pro^ (Figure S2B). The antiviral activity of **S-892216** and other anti-SARS-CoV-2 drugs (ensitrelvir, nirmatrelvir, and remdesivir) were evaluated against recombinant SARS-CoV-2 with amino acid substitutions (D48E, M49K, L50F, N221K, P252L, M49K+P252L, L50F+P252L, D48E+L50F+P252L, and M49K+N221K+P252L) in 3CL^pro^, as observed in the **S-892216** resistance isolation test using VeroE6/TMPRSS2 cells. Because the single basic amino acid substitution of M49K reduced susceptibility to **S-892216** (FC value 3.66), a similar basic amino acid substitution M49R mutant was also prepared and evaluated. FC values of **S-892216** against SARS-CoV-2 with these mutations ranged between 1.22 and 13.7 (Table 5). The binding mode of **S-892216** explained the large fold changes against M49K and M49R mutants, given that key hydrophobic interaction with the M49 side chain was lost upon substitution with basic amino acids, resulting in desolvation penalty upon binding (Figure 2). On the other hand, FC values of other anti-SARS-CoV-2 drugs (nirmatrelvir, ensitrelvir, and remdesivir) against these viruses ranged between 0.389 and 2.35. These results suggested that currently approved 3CL^pro^ inhibitors (nirmatrelvir and ensitrelvir) and **S-892216** do not exhibit cross-resistance; in other words, these inhibitors would be complementary to each other against resistant variants that may emerge in clinical settings in the future.

**Table 5.**
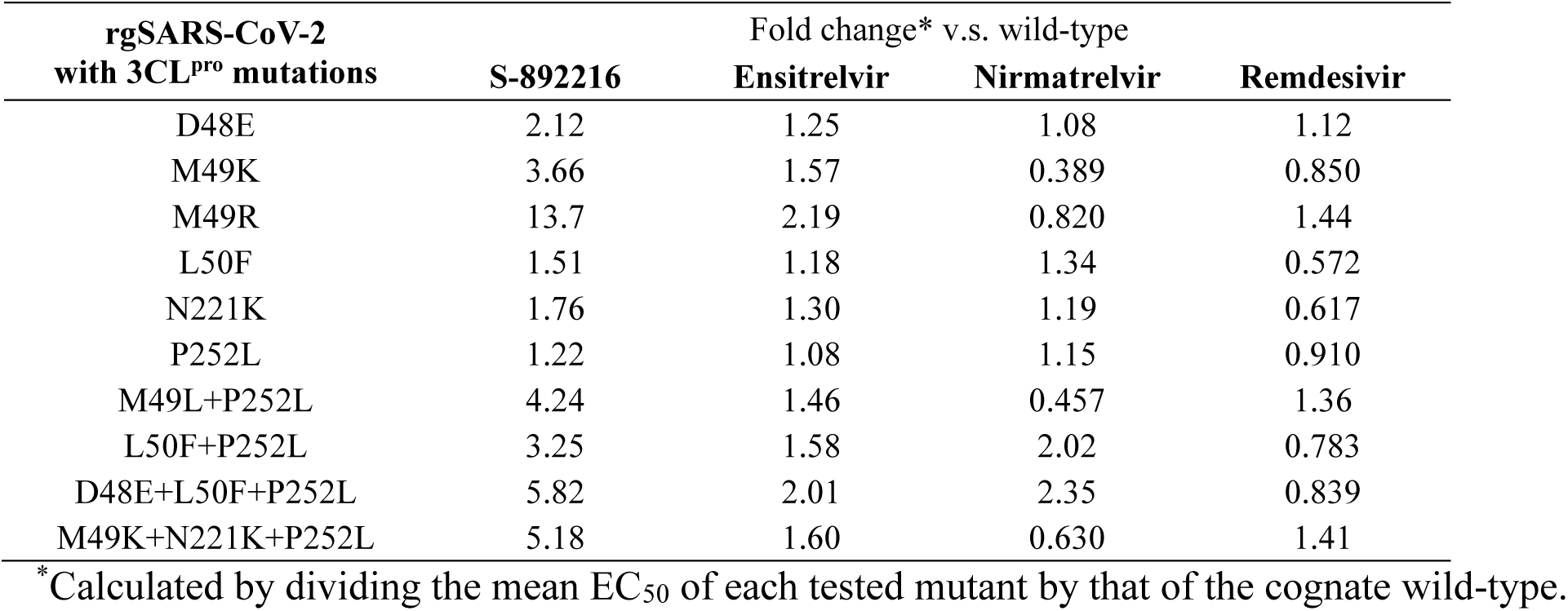
Drug susceptibility of rgSARS-CoV-2 with 3CL^pro^ mutations associated with reduced susceptibility to S-892216.

We examined the antiviral efficacy of **S-892216** *in vivo* in mice infected with SARS-CoV-2 (Figure 5). To evaluate the effect of delayed treatment with **S-892216** against infection with the SARS-CoV-2 Gamma strain (hCoV-19/Japan/TY7-501/2021), **S-892216** was orally administered to mice 24 h after infection, and lung virus titers were measured 48 h after the first administration. **S-892216** reduced lung virus titers in a dose-dependent manner. Virus titer reduction was observed in the 0.3 mg/kg treatment group (Dunnett’s test *p*<0.01), and the 3 mg/kg treatment group appeared to reach maximum efficacy. The plasma concentration increased dose-dependently between 0.03 and 30 mg/kg after a single oral administration in the infected mice. The plasma concentration of 0.3 mg/kg twice a day (bis in die; bid) in mice was estimated to exceed the protein-adjusted EC_90_ (EC_90_ in hAECs×mouse potency shift: 2.31-2.41 nM×11.7-fold=27.0-28.1 nM=14.1-14.7 ng/mL) over time, indicating the importance of the free plasma concentration for *in vivo* efficacy. The virus titer of the group treated with 1 mg/kg of **S-892216** was almost comparable with that of the group treated with 32 mg/kg ensitrelvir, indicating that **S-892216** is more than 30-fold more potent than ensitrelvir. These results strongly support the efficacy of the **S-892216** *in vivo*.

**Figure 5.**
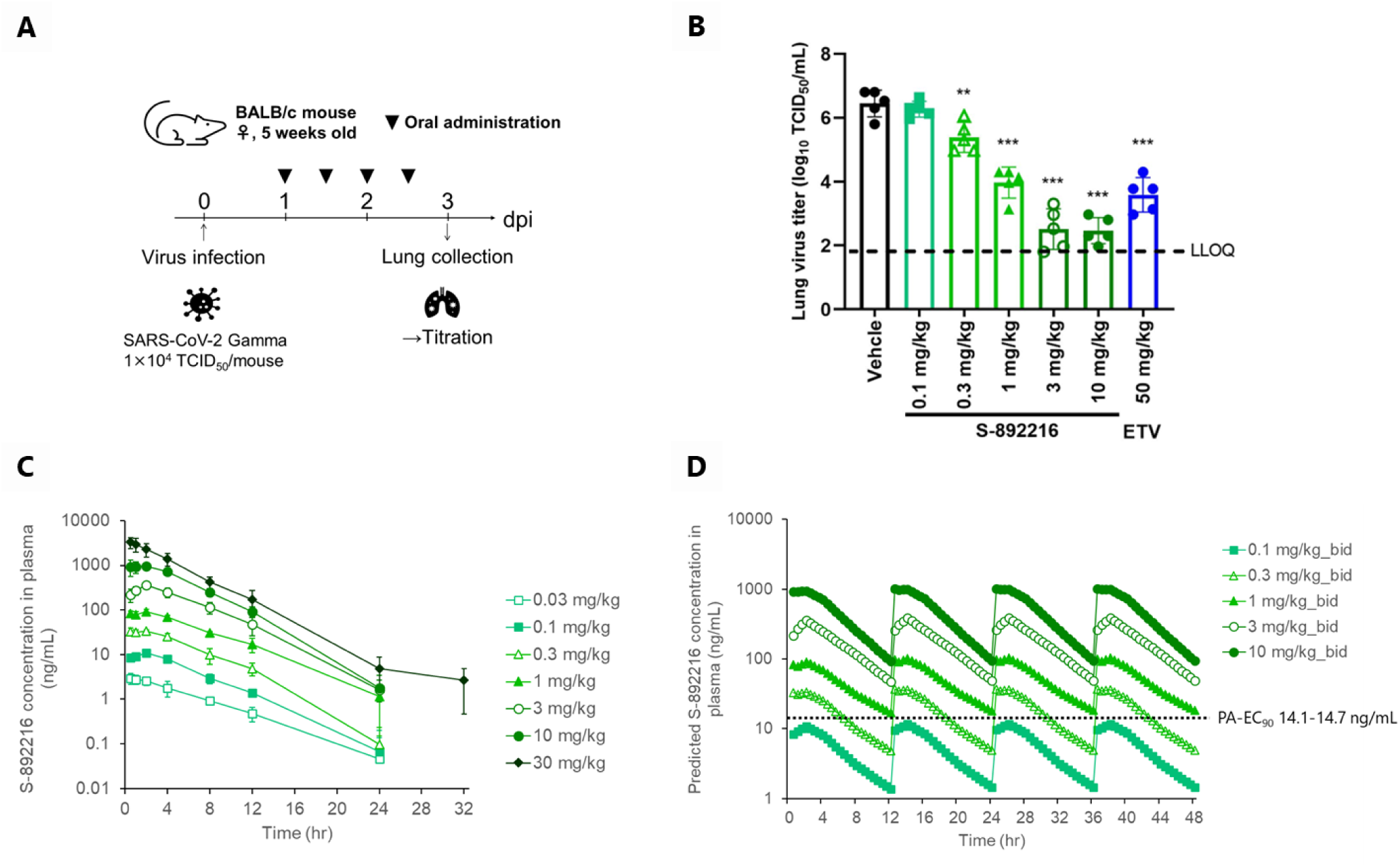
Dose-dependent *in vivo* antiviral efficacy of **S-892216** in mice infected with SARS-CoV-2. (A) Schematic diagram illustrating the *in vivo* study protocol. (B) Effect of **S-892216** (0.1, 0.3, 1, 3, and 10 mg/kg twice a day [bis in die; bid]) and ETV (ensitrelvir fumaric acid, 50 mg/kg [as a free form] bid) treatment on lung virus titers in SARS-CoV-2 Gamma strain (hCoV-19/Japan/TY7-501/2021)-infected mice. Each point represents an individual viral titer (n=5). The broken line represents the LLOQ (1.80 log_10_ TCID_50_/mL). The following p-values were calculated using Dunnett’s test: \*\**p* < 0.01 and \*\**p* < 0.001 vs vehicle. LLOQ=lower limit of quantification. (C) **S-892216** plasma concentration in the infected mice after a single oral administration (n=4). (D) Simulated **S-892216** plasma concentrations after repeated oral administration of **S-892216** twice daily in infected mice as per nonparametric superposition. PA-EC_90_=protein-adjusted EC_90_ extrapolated to 100% mouse serum.

## Chemistry

Scheme 1 illustrates the synthesis scheme for benzyl-type compounds **2-12** *via* the ring synthesis approach. Beginning with the corresponding benzyl bromides **19a-c**, alkylation with dimethyl malonate was conducted to yield **20a-c**. The reaction of trichloroacetyl isocyanate with amino pyridines **21a-c** and ammonia furnished urea derivatives **22a-b**. Cyclization of **20a-c** and **22a-b** resulted in **23a-d**, which then underwent deoxychlorination to furnish **24a-d**. Compound **2** was synthesized from **24a** with isoindoline *via* S_N_Ar amine substitution. Compound **3** was prepared from **24a** *via* alkylation with methyl iodide to produce **25e**, followed by the introduction of isoindoline. Similarly, compounds **4**-**12** were synthesized *via* alkylation with 2-bromoacetonitrile, followed by S_N_Ar substitution with the corresponding amines *via* intermediates **25a-d**.

**Scheme 1.**
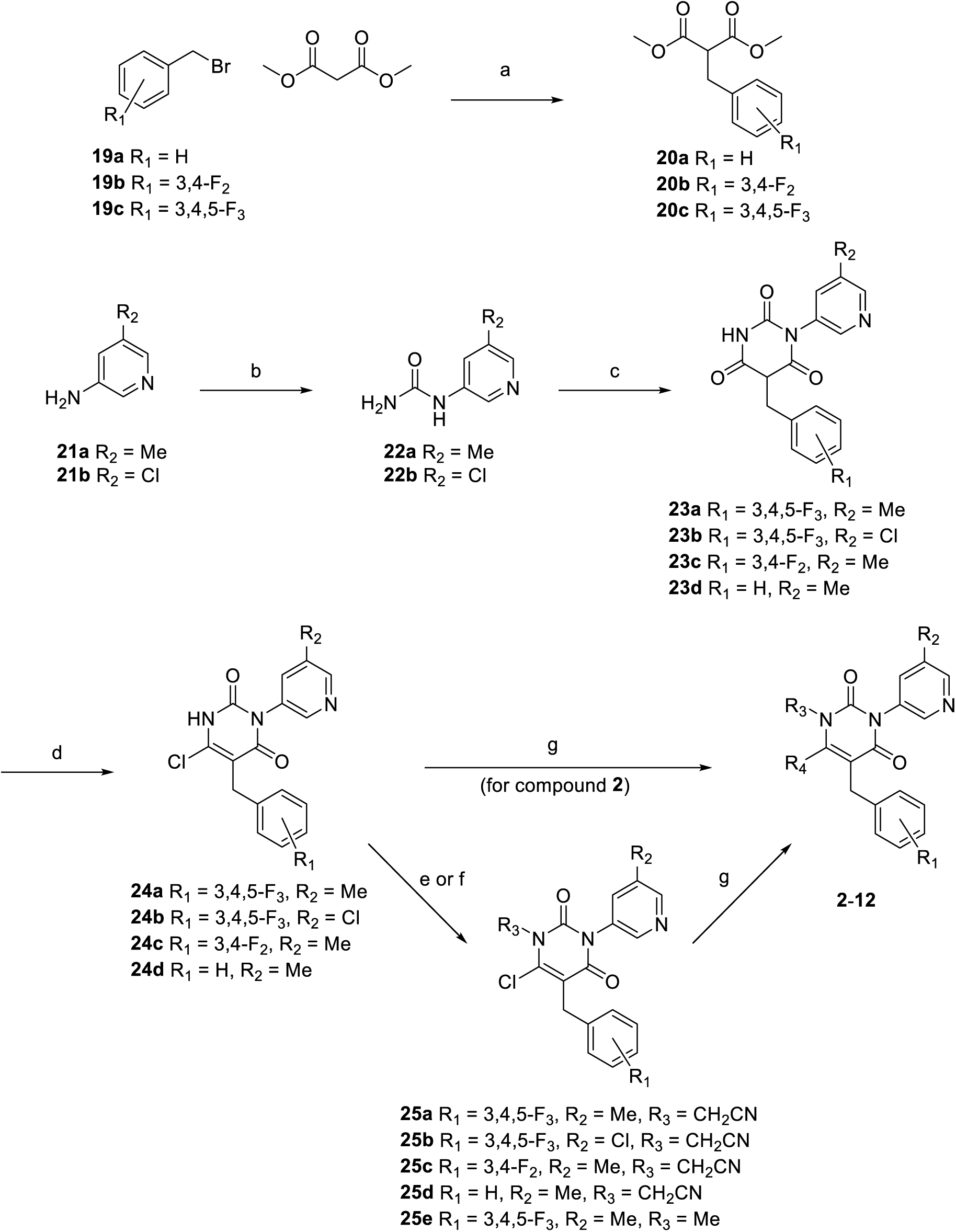
Synthetic scheme of benzyl type pyrimidine-dione derivatives **2**-**12** (a) Corresponding benzyl bromide **19a**-**c**, K_2_CO_3_, acetone, 45°C; (b) trichloroacetyl isocyanate, THF then NH_3_ aq., 0°C to rt, or phenyl chloroformate, pyridine, MeCN then NH_3_ aq, 0°C to rt; (c) **20a**-**c**, 20% NaOEt in EtOH, 90°C; (d) phosphorus oxychloride, H_2_O, rt to 100°C; (e) 2-bromoacetonitrile, DIPEA, DMF, rt; (f) K_2_CO_3_, MeI, DMF, rt; (g) corresponding amine, solvent, 80-150°C

The synthesis scheme for aryl-type compounds **13**-**18** *via* pyrimidine ring substitution is depicted in Scheme 2. Negishi cross-coupling of 6-chloro-2,4-dimethoxypyrimidine **26** with bromobenzene derivatives **27a-c** furnished **28a-c**. Subsequently, acidic hydrolysis of dimethoxypyrimidine **28a-c** furnished **29a-c**. The chloropyridine fragment was introduced *via* Cu-catalyzed Ullmann coupling with 3-bromo-5-chloropyridine to yield **30a-c**. The subsequent steps were performed as described in Scheme 1, with **30a-c** derivatized to **13-18** *via* alkylation with 2-bromoacetonitrile, followed by amine substitution.

**Scheme 2.**
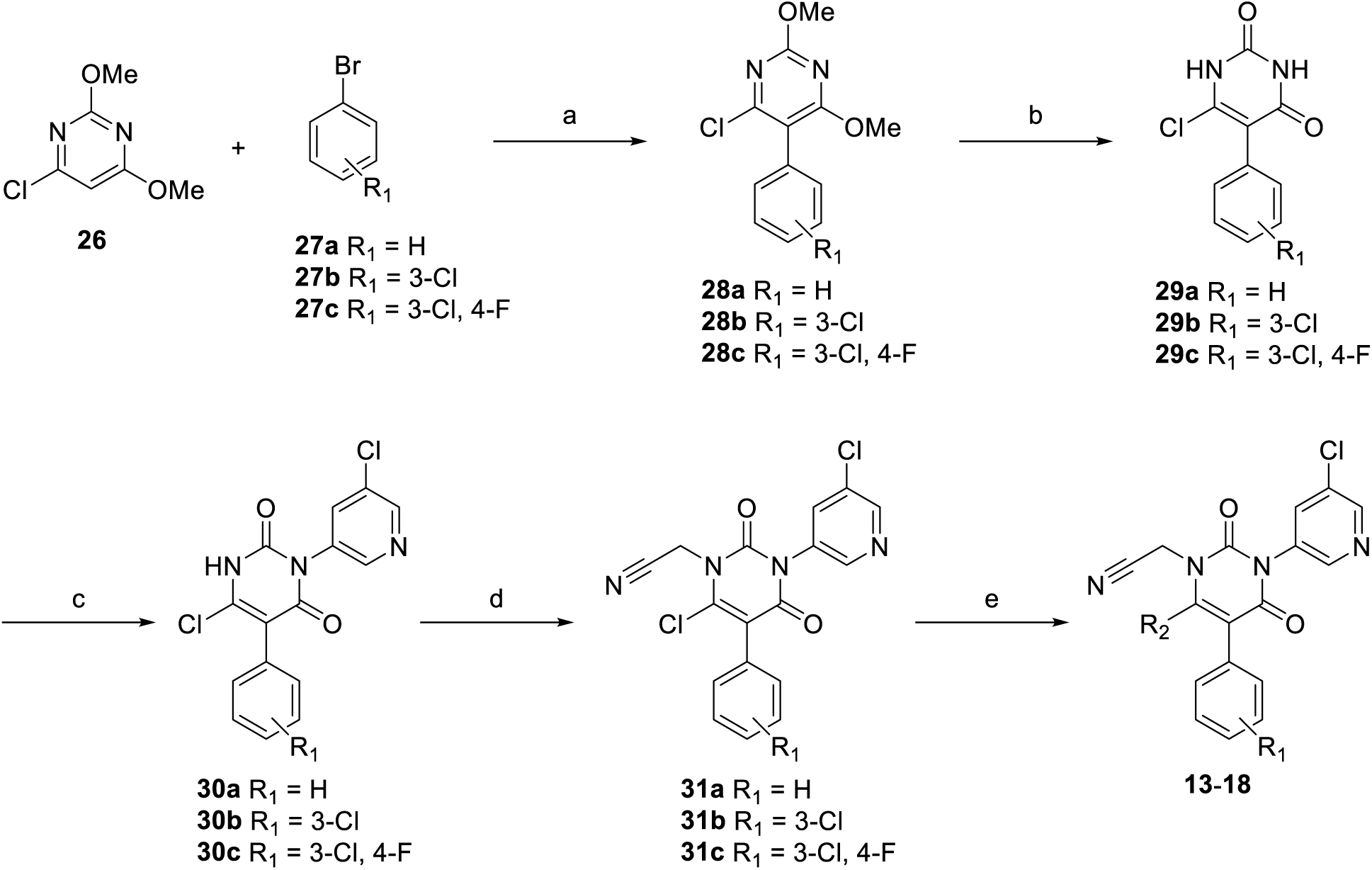
Synthetic scheme of aryl-type pyrimidine-dione derivatives **13**-**18** (a) 6-Chloro-2,4-dimethoxypyrimidine (**26**), *n*-BuLi, ZnCl_2_, THF, -78°C to rt then Pd(PPh_3_)_4_, corresponding bromobenzene (**27a-c**), 80°C; (b) conc. HCl, AcOH, 110°C; (c) 3-bromo-5-chloropyridine, CuI, *N,N’-*dimethylethane-1,2-diamine, K_2_CO_3_, 110°C; (d) 2-bromoacetonitrile, DIPEA, DMF, rt; (e) corresponding amine, DIPEA, DMF, 60°C

## Conclusions

Here, we describe the development of a novel non-peptidic covalent inhibitor, **S-892216** (compound **17**), as a clinical candidate for oral COVID-19 therapy. Starting from compound **1**, which was identified during SAR optimization of the first-generation 3CL^pro^ inhibitor, ensitrelvir, we successfully developed a potent, non-peptidic covalent inhibitor by introducing a nitrile warhead *via* structure-based drug design and side chain modifications. **S-892216** demonstrated potent 3CL^pro^ inhibitory activity and a favorable PK profile for oral treatment without the need for a PK booster. **S-892216** exhibited *in vitro* antiviral activity against various circulating SARS-CoV-2 variants without loss of potency. Furthermore, profiling of rgSARS-CoV-2 with 3CL^pro^ mutations revealed that **S-892216** was less likely to exhibit cross-resistance to the clinically utilized ensitrelvir and nirmatrelvir. *In vivo* efficacy against the SARS-CoV-2 Gamma variant was confirmed in a mouse infection model, demonstrating equivalent activity at a 30-fold lower dose than that of ensitrelvir. No teratogenicity was observed in non-clinical safety studies using pregnant animals (data not shown). Moreover, given the higher potency than ensitrelvir and the need for lower doses, **S-892216** is expected to reduce the limitations of drug-drug interactions (DDI). Clinical trials for **S-892216** are ongoing, and clinical data, including DDI data, will be reported in the near future.

## Experimental Section

### General Chemistry

All commercial reagents and solvents were used as received without further purification. Reactions were monitored *via* thin-layer chromatography performed on Merck silica gel plates (60 F254) or analytical liquid chromatography/mass spectroscopy (LC/MS) performed on a ACQUITY UPLC BEH (C18, 1.7 µm, 2.1 x50 mm, linear gradient from 5% to 100% B over 3.5 min, then 100% B for 0.5 min [A = water + 0.1% formic acid, B = MeCN + 0.1% formic acid], flow rate: 0.8 mL/min) using a Shimadzu Nexera system equipped with LC-20AD binary gradient module and SPD-20AV detector (detection at 255 nm) and SIL-30AC MP sample manager or a Shimadzu UFLC system equipped with a LCMS-2020 mass spectrometer, LC-20AD binary gradient module, SPD-M20A photodiode array detector (detection at 254 nm) and SIL-20AC sample manager. All compounds used in the bioassay are >95% pure by HPLC analysis. Flash column chromatography was performed on an automated purification system using Fuji Silysia prepacked silica gel columns. ^1^H and ^13^C NMR spectra were recorded on a Bruker Advance at 400 and 100 MHz, respectively. Spectral data are reported as follows: chemical shift (as ppm referenced to tetramethylsilane), integration value, multiplicity (s=singlet, d=doublet, t=triplet, q=quartet, m=multiplet, br=broad), and coupling constant. High-resolution mass spectra (HRMS) were recorded on a Thermo Fisher Scientific LTQ Orbitrap using electrospray positive ionization.

### Anti-SARS-CoV-2 reference compounds

Note that ensitrelvir fumaric acid was used in each experiment described as ensitrelvir. Ensitrelvir fumaric acid (Shionogi & Co. Ltd., Osaka, Japan) was synthesized as previously described.^24^ Nirmatrelvir was also synthesized at Shionogi & Co., Ltd according to the literature.^11^ Remdesivir and NHC, the active ingredients of the prodrug molnupiravir, were obtained from MedChemExpress (Monmouth Junction, NJ, USA). The P-gp inhibitor CP-100356 was purchased from Sigma–Aldrich Co. Ltd (St Louis, MO, USA).

### 6-(Isoindolin-2-yl)-3-(5-methylpyridin-3-yl)-1-(3,4,5-trifluorobenzyl)-1,3,5-triazine-2,4(1*H*,3*H*)-dione (1)

A mixture of 3-(5-methylpyridin-3-yl)-6-(methylthio)-1-(3,4,5-trifluorobenzyl)-1,3,5-triazine-2,4(1*H*,3*H*)-dione^33^ (100 mg, 0.254 mmol) and isoindoline (91.0 mg, 0.761 mmol) in DMA (1.00 mL) was stirred at 60°C for 1.5 h. The reaction mixture was quenched with 0.2 N HCl (2.54 mL) and extracted with CHCl_3_ (3×4.00 mL). The combined organic layer was washed with brine, dried over Na_2_SO_4_, filtered and concentrated under vacuum to give a crude product. Silica gel chromatography (CHCl_3_ : MeOH = 10 : 0 to 9.2 : 0.8), followed by amino-functionalized silica gel chromatography (hexane : EtOAc = 10 : 0 to 1 : 9), afforded compound **1** (63.9 mg, 54% yield) as a white solid. ^1^H NMR (400 MHz, CDCl_3_) δ: 2.36 (3H, s), 5.01 (4H, s), 5.26 (2H, s), 6.93 (2H, dd, *J* = 6.8, 6.8 Hz), 7.20-7.24 (2H, m), 7.29-7.32 (2H, m), 7.44 (1H, br s), 8.35 (1H, d, *J* = 1.8 Hz), 8.43 (1H, br s). ^13^C NMR (400 MHz, CDCl_3_) δ: 18.3, 49.1, 56.8, 110.4 (dd, *J* = 16.0, 6.6 Hz), 122.4, 128.4, 131.4, 132.0-132.2 (m), 134.0, 134.5, 136.4, 139.5 (dt, *J* = 251.6, 15.3 Hz), 146.4, 150.3, 151.8 (ddd, *J* = 250.8, 9.5, 3.6 Hz), 152.8, 152.8, 156.5. LC/MS (ESI): m/z = 466.1 [M+H]^+^.

### Dimethyl 2-(3,4,5-trifluorobenzyl)malonate (20c)

5-(Bromomethyl)-1,2,3-trifluorobenzene **19c** (5.85 mL, 44.4 mmol), acetone (40.0 mL), K_2_CO_3_ (9.21 g, 66.7 mmol), and dimethyl malonate (8.39 mL, 66.7 mmol) were mixed, and the mixture thus obtained was stirred at 45°C for 2 h, and then, it was allowed to stand at room temperature for 16 h. The precipitate was filtered off and washed with acetone. The filtrate was concentrated, the residue thus obtained was purified by silica gel column chromatography (hexane : EtOAc = 10 : 0 to 9 : 1), the solvent was distilled off under reduced pressure, and compound **20c** (5.95 g, 21.5 mmol, yield 49%) was obtained. ^1^H NMR (400 MHz, CDCl_3_) δ: 3.15 (2H, d, *J* = 7.8 Hz), 3.59-3.72 (1H, m), 3.73 (6H, s), 6.83 (2H, dd, *J* = 8.3, 6.5 Hz). LC/MS (ESI): m/z = 277.0 [M+H]^+^.

### 1-(5-Methylpyridin-3-yl)urea (22a)

A solution obtained by mixing THF (150 mL) to 5-methylpyridin-3-amine **21a** (10.0 g, 92.5 mmol) was ice-cooled, trichloroacetyl isocyanate (12.1 mL, 102 mmol) was added thereto, and the mixture was stirred at room temperature for 2 h. A 7 mol/L methanol solution of ammonia (16.0 mL, 111 mmol) was added, and the mixture was stirred at room temperature for 3 h. Then, the mixture was stirred at 50°C for 2 h. The solution was concentrated, CHCl_3_ (50.0 mL) were added thereto, and the obtained precipitate was collected by filtration and washed with chloroform. The obtained solid was trituated in ethanol (50 mL), filtered, washed with ethanol. Compound **22a** (11.5 g, 76.1 mmol, yield 82%) was obtained after dried under reduced pressure. ^1^H NMR (400 MHz, DMSO-*d*_6_) δ: 2.24 (3H, s), 5.98 (2H, s), 7.73-7.76 (1H, m), 7.95 (1H, d, *J* = 1.2 Hz), 8.29 (1H, d, *J* = 2.4 Hz), 8.63 (1H, s). LC/MS (ESI): m/z = 152.2 [M+H]^+^.

### 1-(5-Methylpyridin-3-yl)-5-(3,4,5-trifluorobenzyl)pyrimidine-2,4,6(*1H,3H,5H*)-trione (23a)

Compound **20c** (1.33 g, 4.82 mmol), compound **22a** (607 mg, 4.02 mmol), and a 20% ethanol solution of sodium ethoxide (4.66 mL, 12.1 mmol) were mixed, and the obtained solution was stirred at 90°C for 4 h. The reaction solution was cooled in an ice bath, and was neutralized with a 2 mol/L aqueous hydrochloric acid solution (4.0 mL). H_2_O (7.0 mL) was added, and the obtained precipitate was collected by filtration and washed with water and IPE. The residue thus obtained was dried under reduced pressure, and compound **23a** (1.42 g, 3.91 mmol, yield 97%) was obtained. ^1^H NMR (400 MHz, DMSO-*d*_6_, appeared as 7:3 mixture of tautomers) δ: 2.35 (3H, s), 3.52 (2H, s), 4.31-4.37 (0.3H, m), 7.05-7.14 (1.4H, m), 7.15-7.22 (0.7H, m), 7.45-7.50 (0.3H, m), 7.63-7.70 (0.7H, m), 8.19-8.23 (0.3H, m), 8.28-8.34 (0.7H, m), 8.38-8.47 (1H, m), 10.51 (0.7H, br s), 11.71 (0.3H, s). LC/MS (ESI): m/z = 364.1 [M+H]^+^.

### 6-Chloro-3-(5-methylpyridin-3-yl)-5-(3,4,5-trifluorobenzyl)pyrimidine-2,4(*1H,3H*)-dione (24a)

H_2_O (0.317 mL, 17.6 mmol) was carefully dropwised to a mixture of **23a** (1.28 g, 3.52 mmol) and POCl_3_ (6.55 mL, 70.5 mmol) and stirred at rt until exothermic reaction terminates. The resulting solution was stirred at 100°C for 6 h. The reaction solution was poured into ice and stirred until exothermic reaction terminates. Then LiOH aq. (4.0 mol/L, 53.0 mL, 212 mmol) was carefully added to neutralize (∼pH 5). The resulting suspension was filtered, and washed with H_2_O. The obtained solid was dried under reduced pressure to give **24a** (1.20 g, 3.14 mmol, yield 89%) as a crude dark pink powder, which was used for next step without further purification. ^1^H NMR (400 MHz, DMSO-*d*_6_) δ: 2.35 (3H, s), 3.75 (2H, s), 7.23 (2H, dd, *J* = 9.2, 6.8 Hz), 7.63-7.66 (1H, m), 8.34 (1H, d, *J* = 2.1 Hz), 8.46 (1H, d, *J* = 1.3 Hz), 12.61 (1H, br s). LC/MS (ESI): m/z = 381.9 [M+H]^+^.

### 6-(Isoindolin-2-yl)-3-(5-methylpyridin-3-yl)-5-(3,4,5-trifluorobenzyl)pyrimidine-2,4(1*H*,3*H*)-dione (2)

A solution obtained by mixing compound **24a** (70.0 mg, 0.183 mmol), isoindoline (61.8 μL, 0.55 mmol), and ethanol (0.7 mL) was stirred at 100°C for 17 h. The reaction solution was cooled to room temperature, water (0.7 mL) was added. The obtained precipitate was collected by filtration and washed with water and diethyl ether. The residue thus obtained was dried under reduced pressure, and compound **2** (62.0 mg, 0.133 mmol, yield 73%) was obtained. ^1^H NMR (400 MHz, DMSO-*d*_6_) δ: 2.35 (3H, s), 4.04 (2H, s), 4.94 (4H, s), 7.21 (2H, dd, *J* = 8.4, 7.2 Hz), 7.29 (4H, m), 7.57 (1H, s), 8.27 (1H, s), 8.40 (1H, s), 10.40 (1H, br s). ^13^C NMR (100 MHz, DMSO-*d*_6_) δ 17.6, 29.3, 55.2, 85.2, 111.8 (dd, *J* = 15.3, 4.3 Hz), 122.3, 127.5, 132.5, 133.1, 135.5, 137.1, 137.4 (dt, *J* = 111, 20.4 Hz), 140.0-140.2 (m), 147.0, 148.9, 150.1 (ddd, *J* = 245, 10.2, 4.4 Hz), 150.2, 151.1, 164.1. LC/MS (ESI): m/z = 465.1 [M+H]^+^.

### 6-Chloro-1-methyl-3-(5-methylpyridin-3-yl)-5-(3,4,5-trifluorobenzyl)pyrimidine-2,4(1*H*,3*H*)-dione (25e)

To a mixture of compound **24a** (300 mg, 0.786 mmol), K_2_CO_3_ (326 mg, 2.36 mmol), and DMF (3.0 mL), iodomethane (147 μL, 2.36 mmol) was added and stirred at room temperature for 1 h. To the reaction mixture, water (10 mL) was added, and extracted with EtOAc. The organic layer was washed with water and brine, dried over sodium sulfate, and filtered. The filtrate was concentrated, the residue thus obtained was purified by silica gel column chromatography (CHCl_3_ : MeOH = 10 : 0 to 9.5 : 0.5), the solvent was distilled off under reduced pressure, and compound **25e** (174 mg, 0.440 mmol, yield 56%) was obtained. ^1^H NMR (400 MHz, CDCl_3_) δ: 2.41 (3H, s), 3.67 (3H, s), 3.85 (2H, s), 6.98 (2H, dd, *J* = 8.3, 6.6 Hz), 7.35-7.41 (1H, m), 8.30 (1H, d, *J* = 2.3 Hz), 8.51 (1H, d, *J* = 1.3 Hz). LC/MS (ESI): m/z = 396.0 [M+H]^+^.

### 6-(Isoindolin-2-yl)-1-methyl-3-(5-methylpyridin-3-yl)-5-(3,4,5-trifluorobenzyl)pyrimidine-2,4(1*H*,3*H*)-dione (3)

A solution obtained by mixing compound **25e** (44.5 mg, 0.112 mmol), isoindoline (126 μL, 1.12 mmol), and toluene (1.3 mL) was stirred at 130°C for 2 h. DIPEA (29.5 μL, 0.169 mmol) was added, and the mixture was stirred at 130°C for 2 h and at 150°C for 1 h. Then, DIPEA (98.0 μL, 0.562 mmol) was added again, and the mixture was stirred at 150°C for 6 h. The reaction solution was cooled to room temperature, a saturated aqueous citric acid solution (5 mL) and water (10 mL) were added thereto, and the mixture was extracted with EtOAc. The organic layer was washed with a saturated aqueous NaHCO_3_ solution, dried over Na_2_SO_4_, and filtered. The filtrate was concentrated, the residue thus obtained was purified by silica gel column chromatography (CHCl_3_ : MeOH = 10 : 0 to 9.7 : 0.3), the solvent was distilled off under reduced pressure. The residue thus obtained was purified by reversed-phase chromatography, the solvent was distilled off under reduced pressure and compound **3** (27.0 mg, 0.056 mmol, yield 50%) was obtained. ^1^H NMR (400 MHz, CDCl_3_) δ: 2.40 (3H, s), 3.42 (3H, s), 3.67 (2H, s), 4.56 (4H, s), 6.71 - 6.81 (2H, m), 7.28 - 32 (2H, m,), 7.34 - 7.40 (2H, m), 7.43 (1H, s), 8.34 (1H, d, *J* = 1.9 Hz), 8.48 (1H, s). ^13^C NMR (100 MHz, CDCl_3_) δ 18.4, 30.9, 32.5, 57.3, 108.6, 112.0 (dd, *J* = 15.3, 5.9 Hz), 122.9, 128.1, 131.8, 134.0, 136.0 (dt, *J* = 6.6, 4.4 Hz), 136.6, 137.1, 138.4 (dt, *J* = 249, 15.4 Hz), 146.5, 150.3, 151.2 (ddd, *J* = 249, 9.9, 4.0 Hz), 1520, 154.3, 164.0. LC/MS (ESI): m/z = 479.2 [M+H]^+^.

### 2-(6-Chloro-3-(5-methylpyridin-3-yl)-2,4-dioxo-5-(3,4,5-trifluorobenzyl)-3,4-dihydropyrimidin-1(2*H*)-yl)acetonitrile (25a)

A solution obtained by mixing compound **24a** (200 mg, 0.524 mmol) and DMF (2.0 mL) was ice-cooled, DIPEA (275 μL, 1.57 mmol) and 2-bromoacetonitrile (105 μL, 1.57 mmol) were added thereto, and the obtained solution was stirred at room temperature for 4 h. Then the reaction solution was cooled in an ice bath, a 2 mol/L aqueous hydrochloric acid solution (0.8 mL) and water (6.0 mL) were added thereto, and the mixture was extracted with EtOAc. The organic layer was washed with a saturated aqueous NaHCO_3_ solution and brine, dried over Na_2_SO_4_, and filtered. The filtrate was concentrated, and EtOAc (1.0 mL) and hexane (3.0 mL) were added thereto. The obtained precipitate was collected by filtration and washed with IPE. The residue thus obtained was dried under reduced pressure, and compound **25a** (157 mg, 0.374 mmol, yield 71%) was obtained. ^1^H NMR (400 MHz, CDCl_3_) δ: 2.43 (3H, s), 3.87 (2H, s), 5.08 (2H, s), 6.94-7.02 (2H, dd, *J* = 8.0, 6.8 Hz), 7.39-7.41 (1H, m), 8.31 (1H, d, *J* = 2.1 Hz), 8.55 (1H, d, *J* = 1.2 Hz). LC/MS (ESI): m/z = 420.9 [M+H]^+^.

### 2-(6-(Isoindolin-2-yl)-3-(5-methylpyridin-3-yl)-2,4-dioxo-5-(3,4,5-trifluorobenzyl)-3,4-dihydropyrimidin-1(2*H*)-yl)acetonitrile (4)

A solution obtained by mixing compound **25a** (51.5 mg, 0.122 mmol), isoindoline (41.3 μL, 0.367mmol), and ethanol (1.0 mL) was stirred at 100°C for 2 h. The reaction solution was cooled to room temperature, water (2 mL) was added thereto, and the mixture was extracted with CHCl_3_. The organic layer was dried over sodium sulfate, and filtered. The filtrate was concentrated, the residue thus obtained was purified by silica gel column chromatography (CHCl_3_ : MeOH = 10 : 0 to 9.5 : 0.5), the solvent was concentrated, and IPE (1.0 mL) was added thereto. The obtained precipitate was collected by filtration and washed with IPE. The residue thus obtained was dried under reduced pressure, and compound **4** (6.10 mg, 0.012 mmol, yield 10%) was obtained. ^1^H NMR (400 MHz, CDCl_3_) δ: 2.42 (3H, s), 3.57 (2H, s), 4.64 (4H, s), 4.95 (2H, s), 6.64 (2H, dd, *J* = 7.6, 6.8 Hz), 7.30 (2H, dd, *J* = 5.6, 3.6 Hz), 7.39 (2H, dd, *J* = 6.0, 3.2 Hz), 7.45 (1H, s), 8.35 (1H, d, *J* = 1.6 Hz), 8.51 (1H, s). ^13^C NMR (100 MHz, CDCl_3_) δ: 18.4, 30.5, 32.3, 58.9, 110.4, 112.0 (dd, *J* = 16.0, 5.9 Hz), 114.9, 122.9, 128.5, 131.1, 134.2, 134.6 (dt, *J* = 7.3 4.4 Hz), 136.4, 136.7, 138.6 (dt, *J* = 249, 15.3 Hz), 146.3, 150.8, 150.9, 151.2 (ddd, *J* = 249, 9.9, 4.0 Hz), 154.0, 163.4. LC/MS (ESI): m/z = 504.1 [M+H]^+^.

### 2-(3-(5-Methylpyridin-3-yl)-2,4-dioxo-6-(pyrrolidin-1-yl)-5-(3,4,5-trifluorobenzyl)-3,4-dihydropyrimidin-1(2*H*)-yl)acetonitrile (5)

A solution obtained by mixing compound **25a** (50.0 mg, 0.119 mmol), pyrrolidine (14.7 μL, 0.178mmol), DIPEA (62.3 μL, 0.356mmol), and DMF (1.0 mL) was stirred at 80°C for 2 h. The reaction solution was cooled to room temperature, a saturated aqueous citric acid solution (0.2 mL) and water (2 mL) were added thereto. The obtained precipitate was collected by filtration and washed with water. The residue thus obtained was dried under reduced pressure, and compound **5** (35.0 mg, 0.077 mmol, yield 65%) was obtained. ^1^H NMR (400 MHz, CDCl_3_) δ: 2.01-2.12 (4H, m), 2.40 (3H, s), 3.26-3.32 (4H, m), 3.71 (2H, s), 4.85 (2H, s), 6.80 (2H, dd, *J* = 8.0, 6.8 Hz), 7.41 (1H, s), 8.32 (1H, d, *J* = 1.9 Hz), 8.49 (1H, s). ^13^C NMR (100 MHz, DMSO-*d*_6_) δ: 17.6, 25.4, 29.5, 33.3, 50.5, 104.3, 112.0 (dd, *J* = 15.7, 4.8 Hz), 116.5, 132.3, 133.4, 136.7, 136.9 (dt, *J* = 245, 15.3 Hz), 137.6-137.8 (m), 146.5, 149.4, 150.0 (ddd, *J* = 246, 9.5, 4.4 Hz), 151.2, 153.1, 163.3. LC/MS (ESI): m/z = 456.1 [M+H]^+^.

### 2-(3-(5-Methylpyridin-3-yl)-2,4-dioxo-5-(3,4,5-trifluorobenzyl)-6-(4-(trifluoromethyl)piperidin-1-yl)-3,4-dihydropyrimidin-1(2*H*)-yl)acetonitrile (6)

A solution obtained by mixing compound **25a** (25.0 mg, 0.059 mmol), 4-(trifluoromethyl)piperidine (14.0 mg, 0.089 mmol), DIPEA (31.1 μL, 0.178 mmol), and DMF (0.25 mL) was stirred at 80°C for 2 h. The reaction solution was cooled to room temperature, water was added thereto, and the mixture was extracted with CHCl_3_. The organic layer was separated and concentrated. The residue was purified by reverse phase HPLC to give compound **6** (10.0 mg, 0.019 mmol, yield 32%). ^1^H NMR (400 MHz, DMSO-*d_6_*) δ: 1.62-1.89 (4H, m), 2.35 (3H, s), 2.84-3.07 (2H, m), 3.12-3.26 (3H, m), 3.32-3.35 (1H, m) 3.79 (2H, s), 4.92 (2H, s), 7.27 (2H, dd, *J* = 8.5, 7.3 Hz), 7.57 (1H, s), 8.28 (1H, d, *J* = 1.6 Hz), 8.44 (1H, s). ^13^C NMR (100 MHz, DMSO-*d_6_*) δ: 17.6, 24.0, 29.1, 32.6, 38.2 (q, *J* = 27.1 Hz), 47.9, 106.5, 112.2 (dd, *J* = 16.0, 4.4 Hz), 116.5, 127.7 (q, *J* = 278.5 Hz), 132.2, 133.5, 136.6, 136.9 (dt, *J* = 246.1, 15.3 Hz), 137.5-137.8 (m), 146.4, 149.5, 150.0 (ddd, *J* = 245.8, 9.6, 3.5 Hz), 151.1, 154.9, 163.2. LC/MS (ESI): m/z = 538.2 [M+H]^+^.

### 2-(3-(5-Methylpyridin-3-yl)-2,4-dioxo-6-(7-oxa-2-azaspiro[3.5]nonan-2-yl)-5-(3,4,5-trifluorobenzyl)-3,4-dihydropyrimidin-1(2*H*)-yl)acetonitrile (7)

A solution obtained by mixing compound **25a** (25.0 mg, 0.059 mmol), 7-oxa-2-azaspiro[3.5]nonane (11.0 mg, 0.089 mmol), DIPEA (31.1 μL, 0.178 mmol), and DMF (0.25 mL) was stirred at 80°C for 2 h. The reaction solution was cooled to room temperature, water was added thereto, and the mixture was extracted with CHCl_3_. The organic layer was separated and concentrated. The residue was purified by reverse phase HPLC to give compound **7** (16.4 mg, 0.032 mmol, yield 54%). ^1^H-NMR (400 MHz, DMSO-*d_6_*) δ: 1.62 (4H, t, *J* = 4.9 Hz), 2.34 (3H, s), 3.43 (4H, t, *J* = 4.9 Hz), 3.84 (2H, s), 4.09 (4H, s), 4.79 (2H, s), 7.28 (2H, dd, *J* = 9.1, 6.9 Hz), 7.51 (1H, s), 8.23 (1H, d, *J* = 2.0 Hz), 8.41 (1H, s). ^13^C NMR (100 MHz, DMSO-*d_6_*) δ: 17.6, 28.4, 32.4, 35.0, 36.2, 63.7, 65.8, 89.0, 112.0 (dd, *J* = 16.0, 4.4 Hz), 116.7, 132.4, 133.3, 136.8 (dt, *J* = 246, 15.6 Hz), 136.8, 139.3-139.5 (m), 146.6, 149.1, 150.1 (ddd, *J* = 247, 9.8, 4.0 Hz), 151.6, 155.2, 162.6. LC/MS (ESI): m/z = 512.6 [M+H]^+^.

### 2-(6-(6,6-Difluoro-2-azaspiro[3.3]heptan-2-yl)-3-(5-methylpyridin-3-yl)-2,4-dioxo-5-(3,4,5-trifluorobenzyl)-3,4-dihydropyrimidin-1(2*H*)-yl)acetonitrile (8)

A solution obtained by mixing compound **25a** (60.0 mg, 0.143 mmol), 6,6-difluoro-2-azaspiro[3.3]heptane 2,2,2-trifluoroacetate (52.9 μL, 0.214 mmol), DIPEA (74.7 μL, 0.428 mmol) and DMF (1.2 mL) was stirred at 100°C for 2 h. The reaction solution was cooled to room temperature, water (2.4 mL) was added thereto, and the mixture was extracted with EtOAc. The organic layer was dried over Na_2_SO_4_, and filtered. The filtrate was concentrated, the residue thus obtained was purified by silica gel column chromatography (CHCl_3_ : MeOH = 10 : 0 to 9.5 : 0.5), the solvent was distilled off under reduced pressure, and compound **8** (52.5 mg, 0.101 mmol, yield 71%) was obtained. ^1^H NMR (400 MHz, DMSO-*d_6_*) δ: 2.34 (3H, s), 2.79 (4H, t, *J* = 12.4 Hz), 3.82 (2H, s), 4.42 (4H, s), 4.76 (2H, s), 7.24 (1H, d, *J* = 5.2 Hz), 7.25 (1H, d, *J* = 5.2 Hz), 7.51 (1H, s), 8.24 (1H, br), 8.42 (1H, br). ^13^C NMR (100 MHz, DMSO-*d*_6_) δ: 17.6, 27.1 (t, *J* = 11.0 Hz), 28.4, 36.2, 44.6 (t, *J* = 22.6 Hz), 66.3, 89.5, 111.9 (dd, *J* =16.1, 4.4 Hz), 116.5, 119.5 (t, *J* = 275 Hz), 132.3, 133.3, 136.8, 136.9 (dt, *J* = 247, 15.8 Hz), 139.7-139.8 (m), 146.6, 149.2, 150.2 (ddd, *J* = 246, 9.9, 4.0 Hz), 151.6, 155.2, 162.6. LC/MS (ESI): m/z = 518.2 [M+H]^+^.

### 2-(3-(5-Methylpyridin-3-yl)-2,4-dioxo-6-(6-oxa-2-azaspiro[3.4]octan-2-yl)-5-(3,4,5-trifluorobenzyl)-3,4-dihydropyrimidin-1(2*H*)-yl)acetonitrile (9)

A solution obtained by mixing compound **25b** (60.0 mg, 0.143 mmol), 6-oxa-2-azaspiro[3.4]octane oxalate (43.5 mg, 0.214 mmol), DIPEA (100 uL, 0.570 mmol), and ethanol (1.2 mL) was stirred at 100 °C for 1 h. The reaction solution was cooled to room temperature and then concentrated. The residue thus obtained was purified by silica gel column chromatography (CHCl_3_ : MeOH = 10 : 0 to 9 : 1, then EtOAc : MeOH = 10 : 0 to 8 : 2), the solvent was distilled off under reduced pressure, and compound **9** (7.4 mg, 0.015 mmol, yield 10%) was obtained. ^1^H NMR (400 MHz, DMSO-*d_6_*) δ: 2.04 (2H, t, *J* = 7.1 Hz), 2.34 (3H, s), 3.65 (2H, t, *J* = 7.1 Hz), 3.70 (2H, s), 3.82 (2H, s), 4.30 (4H, s), 4.78 (2H, s), 7.27 (2H, dd, *J* = 9.1, 6.9 Hz), 7.51 (1H, s), 8.23 (1H, d, *J* = 2.1 Hz), 8.42 (1H, s). ^13^C NMR (100 MHz, DMSO-*d_6_*) δ: 17.6, 28.4, 36.2, 36.8, 40.3, 65.6, 66.7, 75.7, 89.4, 112.0 (dd, *J* = 15.8, 4.8 Hz), 116.7, 132.4, 133.3, 136.7, 136.8 (dt, *J* = 247, 15.8 Hz), 139.3-139.4 (m), 146.6, 149.2, 150.2 (ddd, *J* = 246.3, 9.9, 4.0 Hz), 151.6, 155.4, 162.6. LC/MS (ESI): m/z = 498.1 [M+H]^+^.

### 1-(5-Chloropyridin-3-yl)urea (22b)

5-Chloropyridin-3-amine (20 g, 156 mmol), and pyridine (18.8 mL, 233 mmol) were dissolved in MeCN (120 ml), and cooled to 0°C. Phenyl chloroformate (26.8 g, 171 mmol) was dropwised over 40 min. The resulting reaction mixture was dropwised to 28% aqueous ammonia in a separate flask over 40 min at rt. The resulting suspension was stirred 40 min at rt. The precipitate was filtered, washed with 50% MeCN/H_2_O (80 mL) and dried to give compound **22b** (24.0 g, 140 mmol, yield 90%). ^1^H NMR (400 MHz, DMSO-*d_6_*) δ: 6.16 (2H, s), 8.12-8.17 (2H, m), 8.36-8.40 (1H, m), 8.98 (1H, m). LC/MS (ESI): m/z = 171.8 [M+H]^+^.

### 1-(5-Chloropyridin-3-yl)-5-(3,4,5-trifluorobenzyl)pyrimidine-2,4,6(*1H,3H,5H*)-trione (23b)

**22b** (1.00 g, 5.83 mmol), **20c** (2.42 g, 8.74 mmol) and sodium ethoxide 20% ethanol solution (6.75 mL, 17.5 mmol) were stirred at 90°C for 7 h. Then, the reaction mixture was cooled to 0°C and HCl (2 mol/L) was added. The mixture was extracted with EtOAc, washed with H_2_O and brine, and dried over Na_2_SO_4_. The organic layer was concentrated to give precipitate, which is tritueated and filterd with EtOAc to give **23b** (1.74 g, 4.44 mmol, yield 76%) as a light yellow solid. ^1^H NMR (400 MHz, CDCl_3_) δ: 3.52 (2H, d, *J* = 2.9 Hz), 3.98 (1H, t, *J* = 4.8 Hz), 6.88 (2H, dd, *J* = , 7.1 Hz), 7.44 (1H, t, *J* = 2.0 Hz), 8.13 (1H, br s), 8.24 (1H, d, *J* = 2.1 Hz), 8.67 (1H, d, *J* = 2.0 Hz). LC/MS (ESI): m/z = 384.0 [M+H]^+^.

### 6-Chloro-3-(5-chloropyridin-3-yl)-5-(3,4,5-trifluorobenzyl)pyrimidine-2,4(*1H,3H*)-dione (24b)

H_2_O (0.386 mL, 21.45 mmol) was carefully dropwised to a mixture of **23b** (1.68 g, 4.29 mmol) and POCl_3_ (7.97 mL, 86 mmol) and stirred at rt until exothermic reaction terminates. The resulting solution was stirred at 100°C for 7 h. The reaction solution was poured into ice and stirred until exothermic reaction terminates. Then LiOH aq. (4 mol/L, 80.0 mL, 322 mmol) was carefully added to neutralize (∼pH 5). The resulting suspension was filtered, and washed with H_2_O. The obtained solid was dried under reduced pressure to give **24b** (1.58 g, 3.93 mmol, yield 92%) as a crude off-white solid, which was used for next step without further purification. ^1^H NMR (400 MHz, DMSO-*d_6_*) δ: 3.76 (2H, s), 7.23 (2H, dd, *J* = 7.8, 7.8 Hz), 8.06 (1H, s), 8.53 (1H, s), 8.70 (1H, s), 12.72 (1H, br s). LC/MS (ESI): m/z = 402.0 [M+H]^+^.

### 2-(6-Chloro-3-(5-chloropyridin-3-yl)-2,4-dioxo-5-(3,4,5-trifluorobenzyl)-3,4-dihydropyrimidin-1(2*H*)-yl)acetonitrile (25b)

To a suspension of **24b** (1.47 g, 3.66 mmol) and K_2_CO_3_ (1.52 g, 11.0 mmol) in DMF (11.8 mL), bromoacetonitrile (0.764 mL, 11.0 mmol) was added and stirred at rt for 4 h. The reaction mixture was cooled in an ice-bath, and quenched with cold water. The resulting mixture was extracted with EtOAc, washed with H_2_O and brine, and dried over Na_2_SO_4_. The resulting solution was evaporated, and an obtained slurry was trituated CHCl_3_. The precipitate was collected by filtration, and washed with CHCl_3_ and EtOAc. Compound **25b** (1.25 g, 2.80 mmol, yield 65%) was obtained as a pale brown solid after drying. ^1^H NMR (400 MHz, CDCl_3_) δ: 3.86 (2H, s), 5.07 (2H, s), 6.96 (2H, dd, *J* = 7.2, 7.2 Hz), 7.63 (1H, dd, *J* = 2.0, 2.0 Hz), 8.40 (1H, d, *J* = 1.6 Hz), 8.67 (1H, d, *J* = 1.6 Hz). LC/MS (ESI): m/z = 440.9 [M+H]^+^.

### 2-(3-(5-Chloropyridin-3-yl)-2,4-dioxo-6-(6-oxa-2-azaspiro[3.4]octan-2-yl)-5-(3,4,5-trifluorobenzyl)-3,4-dihydropyrimidin-1(2*H*)-yl)acetonitrile (10)

To a solution of **25b** (150 mg, 0.337 mmol) in DMF (3.0 mL), 6-oxa-2-azaspiro[3.4]octane HCl salt (57.1 mg, 0.505 mmol) and DIPEA (0.18 mL, 1.01 mmol) were added and stirred at 80°C for 2 h. Then, 6-oxa-2-azaspiro[3.4]octane HCl salt (19.0 mg, 0.168 mmol) and DIPEA (0.059 mL, 0.34 mmol) were added and stirred at 90°C for 1 h. After cooling, the reaction mixture was extracted with EtOAc, washed with H_2_O and brine, dried over Na_2_SO_4_. The resulting solution was evaporated, and formed solid was trituated in EtOAc and MeOH to give compound **10** (89.0 mg, 0.168 mmol, yield 50%) as a yellow solid. ^1^H NMR (400 MHz, DMSO-*d_6_*) δ: 2.03 (2H, t, *J* = 6.8 Hz), 3.65 (2H, t, *J* = 6.8 Hz), 3.70 (2H, s), 3.83 (2H, s), 4.31 (4H, s), 4.80 (2H, s), 7.26 (2H, dd, *J* = 7.8, 7.8 Hz), 7.95 (1H, s), 8.45 (1H, s), 8.68 (1H, s). ^13^C NMR (100 MHz, DMSO-*d_6_*) δ: 28.8, 36.7, 37.2, 40.8, 66.1, 67.2, 76.1, 89.6, 112.4 (dd, *J* = 16.1, 4.4 Hz), 117.1, 130.8, 133.9, 137.0, 137.3 (ddd, *J* = 249.0, 15.2, 15.2 Hz), 139.5-139.8 (m), 148.0, 148.5, 150.7 (ddd, *J* = 245.7, 9.5, 3.7 Hz), 151.9, 155.8, 162.8. LC/MS (ESI): m/z = 518.1 [M+H]^+^.

### Dimethyl 2-(3,4-difluorobenzyl)malonate (20b)

To a mixture of 4-(bromomethyl)-1,2-difluorobenzene (**19b)** (3.00 g, 14.49 mmol) and dimethyl malonate (2.87 g, 21.74 mmol) in acetone (12 mL) was added K_2_CO_3_ (3.00 g, 21.74 mmol), and stirred at 45°C for 2 h then 65°C for 6 h. After cooling, the resulting suspension was filtered, and the filtrate was concentrated. The crude residue was purified by silica gel column chromatography (hexane/EtOAc = 10 : 0 to 9 : 1) to give **20b** (2.12 g, 8.13 mmol, yield 56%) as a colorless oil. ^1^H NMR (400 MHz, CDCl_3_) δ: 3.18 (2H, d, *J* = 7.8 Hz), 3.62 (1H, t, *J* = 7.8 Hz), 3.72 (6H, s), 6.89-6.95 (1H, m), 6.99-7.11 (2H, m). LC/MS (ESI): m/z = 259.0 [M+H]^+^.

### 5-(3,4-Difluorobenzyl)-1-(5-methylpyridin-3-yl)pyrimidine-2,4,6(*1H,3H,5H*)-trione (23c)

**22a** (0.800 g, 5.29 mmol), **20b** (2.07 g, 7.94 mmol) and sodium ethoxide 20% ethanol solution (6.14 ml, 15.88 mmol) were stirred at 90°C for 1 h. Then, the reaction mixture was cooled to 0°C and HCl (2 mol/L, ca. 5 mL) was added. The reaction mixture was basicified with aqueous NaHCO_3_ to ∼pH 8. The mixture was extracted with EtOAc and H_2_O, and organic layer was removed. The aqueous layer was acidified with 2 mol/L HCl (∼pH 4), and precipitate was filtered and dried to give **23c** (1.74 g, 4.94 mmol, yield 93%) as a light yellow solid. ^1^H-NMR (400 MHz, CDCl_3_) δ: 2.40 (3H, s), 3.51-3.58 (2H, m), 6.92-7.18 (5H, m), 8.02-8.12 (2H, m), 8.51 (1H, s). LC/MS (ESI): m/z = 346.1 [M+H]^+^.

### 6-Chloro-5-(3,4-difluorobenzyl)-3-(5-methylpyridin-3-yl)pyrimidine-2,4(*1H,3H*)-dione (24c)

H_2_O (0.435 mL, 24.1 mmol) was carefully dropwised to a mixture of **23c** (1.70 g, 4.82 mmol) and POCl_3_ (8.97 mL, 96.0 mmol) and stirred at rt until exothermic reaction terminates. The resulting solution was stirred at 100°C for 4 h. The reaction solution was poured into ice and stirred until exothermic reaction terminates. Then LiOH aq. (4 mol/L, 90.0 mL, 362 mmol) was carefully added to neutralize (∼pH 5). The resulting suspension was filtered, and washed with H_2_O. The obtained solid was dried under reduced pressure to give **24c** (2.22 g, 6.10 mmol, yield 127%) as a crude off-white solid, which was used for next step without further purification. ^1^H-NMR (400 MHz, CDCl_3_) δ: 2.40 (3H, s), 3.79 (2H, s), 7.04-7.19 (3H, m), 7.40 (1H, s), 8.32 (1H, s), 8.51 (1H, s). LC/MS (ESI): m/z = 364.0 [M+H]^+^.

### 2-(6-Chloro-5-(3,4-difluorobenzyl)-3-(5-methylpyridin-3-yl)-2,4-dioxo-3,4-dihydropyrimidin-1(2*H*)-yl)acetonitrile (25c)

To a suspension of **24c** (2.15 g, 5.91 mmol) and K_2_CO_3_ (1.52 g, 11.0 mmol) in DMF (17.2 mL), bromoacetonitrile (1.24 mL, 17.7 mmol) was added and stirred at rt for 4 h. The reaction mixture was cooled in an ice-bath, and quenched with cold water. The resulting mixture was extracted with EtOAc, washed with H_2_O and brine, and dried over Na_2_SO_4_. The resulting solution was evaporated, and an obtained residue was purified by silica gel column chromatography (hexane/ EtOAc = 10 : 0 to 8 : 2) followed by trituation/filtration in CHCl_3_/EtOAc to give compound **25c** (0.360 g, 0.89 mmol, yield 19% from **23c** in 2 steps) as a pale brown solid. ^1^H-NMR (400 MHz, CDCl_3_) δ: 2.41 (3H, s), 3.88 (2H, s), 5.06 (2H, s), 7.04-7.19 (3H, m), 7.39 (1H, br s), 8.30 (1H, d, *J* = 2.1 Hz), 8.53 (1H, d, *J* = 1.1 Hz). LC/MS (ESI): m/z = 403.0 [M+H]^+^.

### 2-(5-(3,4-Difluorobenzyl)-3-(5-methylpyridin-3-yl)-2,4-dioxo-6-(6-oxa-2-azaspiro[3.4]octan-2-yl)-3,4-dihydropyrimidin-1(2*H*)-yl)acetonitrile (11)

To a solution of **25c** (150 mg, 0.372 mmol) in DMF (3 mL), 6-oxa-2-azaspiro[3.4]octane HCl salt (63.2 mg, 0.559 mmol) and DIPEA (0.195 mL, 1.12 mmol) were added and stirred at 80°C for 2 h. Then, 6-oxa-2-azaspiro[3.4]octane HCl salt (21.1 mg, 0.186 mmol) and DIPEA (0.065 mL, 0.372 mmol) were added and stirred at 90°C for 1 h. After cooling, the reaction mixture was extracted with EtOAc, washed with H_2_O and brine, dried over Na_2_SO_4_. The resulting solution was evaporated, and the residue was purified by silica gel column chromatography (EtOAc/MeOH = 10 : 0 to 8 : 2) followed by trituation/filtration in EtOAc in to give compound **11** (98.9 mg, 0.204 mmol, yield 55%) as a yellow solid. ^1^H-NMR (CDCl_3_) δ: 2.10 (2H, t, *J* = 7.2 Hz), 2.39 (3H, s), 3.80 (2H, s), 3.82 (2H, t, *J* = 7.2 Hz), 3.94 (2H, s), 4.27 (4H, s), 4.77 (2H, s), 6.86-6.92 (1H, m), 6.94-7.02 (1H, m), 7.13 (1H, ddd, *J* = 10.0, 8.4, 8.4 Hz), 7.41 (1H, s), 8.31 (1H, d, *J* = 2.0 Hz), 8.47 (1H, s). ^13^C-NMR (100 MHz, DMSO-*d_6_*) δ: 17.6, 28.1, 36.2, 36.8, 40.4, 65.5, 66.7, 75.7, 90.4, 116.5 (d, *J* = 16.9 Hz), 116.7, 117.1 (d, *J* = 16.9 Hz), 124.1 (dd, *J* = 5.9, 2.9 Hz), 132.4, 133.3, 136.8, 139.5 (dd, *J* = 5.1, 3.7 Hz), 146.6, 147.4 (dd, *J* = 172.1, 12.4 Hz), 149.2, 149.8 (dd, *J* = 173.6, 13.1 Hz), 151.6, 154.9, 162.7. LC/MS (ESI): m/z = 480.1 [M+H]^+^.

### Dimethyl 2-benzylmalonate (20a)

To a mixture of benzyl bromide (**19a)** (3.00 g, 17.5 mmol) and dimethyl malonate (3.48 g, 26.3 mmol) in acetone (12 mL) was added K_2_CO_3_ (3.64 g, 26.3 mmol), and stirred at 45°C for 2 h then 65°C for 6 h. After cooling, the resulting suspension was filtered, and the filtrate was concentrated. The crude residue was purified by silica gel column chromatography (hexane/EtOAc = 10 : 0 to 9 : 1) to give **20a** (3.32 g, 14.86 mmol, yield 85%) as a colorless oil. ^1^H-NMR (400 MHz, CDCl_3_) δ: 3.22 (2H, d, *J* = 7.9 Hz), 3.70 (6H, s), 7.17-7.31 (5H, m). LC/MS (ESI): m/z = 223.1 [M+H]^+^.

### 5-Benzyl-1-(5-methylpyridin-3-yl)pyrimidine-2,4,6(*1H,3H,5H*)-trione (23d)

**22a** (1.20 g, 7.94 mmol), **20a** (2.65 g, 11.9 mmol) and sodium ethoxide 20% ethanol solution (9.21 mL, 23.8 mmol) were stirred at 90°C for 1 h. Then, the reaction mixture was cooled to 0°C and HCl (2 mol/L, 7.94 ml, 15.9 mmol) and H_2_O 24 mL were added. The resulting suspension was filtered and washed with H_2_O and IPE. The collected solid was air-dried to give **23d** (1.18 g, 3.40 mmol, yield 43%) as a light brown solid. ^1^H-NMR (400 MHz, CDCl_3_) δ: 2.37 (3H, s), 3.55 (1H, dd, *J* = 13.4, 4.8 Hz), 3.62 (1H, dd, *J* = 13.4, 4.8 Hz), 3.94 (1H, dd, *J* = 4.8, 4.8 Hz), 7.04 (1H, br s), 7.16-7.20 (2H, m), 7.32-7.38 (3H, m), 7.95 (1H, br s), 8.07 (1H, br s), 8.48 (1H, s). LC/MS (ESI): m/z = 310.1 [M+H]^+^.

### 5-Benzyl-6-chloro-3-(5-methylpyridin-3-yl)pyrimidine-2,4(*1H,3H*)-dione (24d)

H_2_O (0.295 ml, 16.4 mmol) was carefully dropwised to a mixture of **23d** (1.14 g, 3.28 mmol) and POCl_3_ (6.10 ml, 65.6 mmol) and stirred at rt until exothermic reaction terminates. The resulting solution was stirred at 100°C for 3 h. After cooling, H_2_O (11 mL) was carefully added at 0°C and stirred until exothermic reaction terminates. Then LiOH aq. (4 mol/L, 61.5 ml, 246 mmol) was carefully added to neutralize (∼pH 5). The resulting slurry was filtered, and washed with H_2_O. The obtained solid was dried under reduced pressure to give **24d** (1.74 g, 5.31 mmol, yield 162%) as a crude off-white solid, which was used for next step without further purification. ^1^H-NMR (400 MHz, DMSO-*d_6_*) δ: 2.32 (3H, s), 3.69 (2H, s), 7.13-7.17 (1H, m), 7.18-7.30 (5H, m), 7.46 (1H, s), 8.16 (1H, s), 8.35 (1H, s). LC/MS (ESI): m/z = 328.0 [M+H]^+^.

### 2-(5-Benzyl-6-chloro-3-(5-methylpyridin-3-yl)-2,4-dioxo-3,4-dihydropyrimidin-1(*2H*)-yl)acetonitrile (25d)

To a suspension of **24d** (1.63 g, 4.97 mmol) and K_2_CO_3_ (2.06 g, 14.9 mmol) in DMF (13 mL), bromoacetonitrile (1.04 mL, 14.9 mmol) was added and stirred at rt for 2 h. The reaction mixture was cooled in an ice-bath, and quenched with cold water. The resulting mixture was extracted with EtOAc, washed with H_2_O and brine, and dried over Na_2_SO_4_. The resulting solution was evaporated, and an obtained residue was purified by silica gel column chromatography (hexane/EtOAc = 8 : 2 to 0 : 10) followed by trituation/filtration in CHCl_3_/EtOAc to give compound **25d** (0.360 g, 0.981 mmol, yield 30% from **23d** in 2 steps) was obtained as a pale brown solid after drying. ^1^H-NMR (400 MHz, CDCl_3_) δ: 2.41 (3H, s), 3.93 (2H, s), 5.05 (2H, s), 7.23-7.40 (6H, m), 8.30 (1H, d, *J* = 2.3 Hz), 8.52 (1H, br s). LC/MS (ESI): m/z = 367.1 [M+H]^+^.

### 2-(5-Benzyl-3-(5-methylpyridin-3-yl)-2,4-dioxo-6-(6-oxa-2-azaspiro[3.4]octan-2-yl)-3,4-dihydropyrimidin-1(2*H*)-yl)acetonitrile (12)

To a solution of **25d** (150 mg, 0.41 mmol) in DMF (3.00 mL), 6-oxa-2-azaspiro[3.4]octane HCl salt (69.4 mg, 0.61 mmol) and DIPEA (0.214 mL, 1.23 mmol) were added and stirred at 80°C for 2 h. Then, 6-oxa-2-azaspiro[3.4]octane HCl salt (23.1 mg, 0.204 mmol) and DIPEA (0.071 mL, 0.409 mmol) were added and stirred at 90°C for 3 h. After cooling, the reaction mixture was extracted with EtOAc, washed with H_2_O and brine, dried over Na_2_SO_4_. The resulting solution was evaporated, and formed solid was trituated in EtOAc and filtered to give compound **12** (124 mg, 0.278 mmol, yield 68 %) as a yellow solid. ^1^H-NMR (400MHz, CDCl_3_) δ: 2.00 (2H, t, *J* = 7.0 Hz), 2.38 (3H, s), 3.67 (2H, s), 3.77 (2H, t, *J* = 7.0 Hz), 4.02 (2H, s), 4.21 (4H, s), 4.79 (2H, s), 7.16 (2H, d, *J* = 7.4 Hz), 7.21-7.26 (1H, m), 7.31-7.37 (2H, m), 7.42 (1H, br s), 8.33 (1H, d, *J* = 2.0 Hz), 8.47 (1H, s). ^13^C-NMR (100 MHz, DMSO-*d_6_*) δ: 17.6, 28.7, 36.0, 36.7, 40.4, 65.3, 66.7, 75.6, 91.4, 116.7, 125.8, 127.5, 128.4, 132.4, 133.3, 136.8, 141.6, 146.7, 149.2, 151.5, 154.3, 162.7. LC/MS (ESI): m/z = 444.2 [M+H]^+^.

### 4-Chloro-2,6-dimethoxy-5-phenylpyrimidine (28a)

To a mixed solution of a 1.55 mol/L *n*-butyllithium solution in hexane (8.87 mL, 13.8 mmol) and THF (5.00 mL), a THF (6.50 mL) solution of 6-chloro-2,4-dimethoxypyrimidine (**26**) (2.00 g, 11.5 mmol) was added dropwise at -78°C over 15 minutes. The mixture was stirred at -78°C for 1 h. A 2.0 mol/L zinc chloride 2-methyltetrahydrofuran solution (7.16 mL, 14.3 mmol) was added dropwise over 5 minutes. The mixture was stirred at room temperature for 2 h. Bromobenzene (**27a**) (1.38 mL, 13.2 mmol) and tetrakis(triphenylphosphine)palladium (0.662 g, 0.573 mmol) were added, and the mixture was stirred at 80°C for 1.5 h. The reaction solution was cooled to room temperature, water (14.0 mL) and 2.00 mol/L hydrochloric acid (6.00 mL) were added thereto, and the mixture was extracted with EtOAc. The organic layer was concentrated under reduced pressure, and isopropanol (10.0 mL) was added to the obtained residue. The precipitate was collected by filtration and washed with IPA to afford compound **28a** (1.29 g, 5.15 mmol yield 45%). ^1^H NMR (400 MHz, CDCl_3_) δ: 3.94 (3H, s), 4.05 (3H, s), 7.29-7.31 (2H, m), 7.37-7.45 (3H, m). LC/MS (ESI): m/z = 251.0 [M+H]^+^.

### 6-Chloro-5-phenylpyrimidine-2,4(1H,3H)-dione (29a)

Acetic acid (4.29 mL) and concentrated hydrochloric acid (4.29 mL) were added to compound **28a** (1.29 g, 5.15 mmol), and the mixture was stirred at 110°C for 5 h. The reaction solution was cooled to room temperature, and then water (12.0 mL) was added thereto. The precipitate was collected by filtration and washed with water to afford compound **29a** (1.09 g, 4.90 mmol). ^1^H NMR (400 MHz, DMSO-*d*_6_) δ: 7.27 (2H, m), 7.38 (3H, m), 11.49 (1H, brs), 12.11 (1H, brs). LC/MS (ESI): m/z = 222.9 [M+H] ^+^.

### 6-Chloro-3-(5-chloropyridin-3-yl)-5-phenylpyrimidine-2,4(1*H*,3*H*)-dione (30a)

A mixture of compound **29a** (500 mg, 2.25 mmol), 3-bromo-5-chloropyridine (864 mg, 4.49 mmol), copper iodide (428 mg, 2.25 mmol), K_2_CO_3_ (621 mg, 4.49 mmol) in NMP (4.00 mL) was stirred at 110°C, and then a mixture of *N,N’*-dimethylethylenediamine (483 μL, 4.50 mmol) and water (121 μL, 6.74 mmol) in NMP (0.20 ml) was added dropwise. After the resulting mixture was stirred at 110°C for 1 h and 40 minutes, it was allowed to cool in a water bath and then diluted with water (10 mL) and 2 mol/L citric acid aq. (2.50 mL). The precipitate was collected by filteration and washed with IPA. The obtained solid was dried under reduced pressure to afford a crude product of compound **30a** (888 mg). ^1^H NMR (400 MHz, DMSO-*d*_6_) δ: 7.31-7.34 (5H, m), 8.09 (1H, s), 8.65 (1H, br), 8.78 (1H, br). LC/MS (ESI): m/z = 333.9 [M+H]^+^.

### 2-(6-Chloro-3-(5-chloropyridin-3-yl)-2,4-dioxo-5-phenyl-3,4-dihydropyrimidin-1(*2H*)-yl)acetonitrile (31a)

A mixture of compound **30a** (300 mg, 0.754 mmol, 84 % purity), DIPEA (198 μL, 1.13 mmol) , and 2-bromoacetonitrile (79.0 μL, 1.13 mmol) in DMF (1.50 mL) was stirred at 60°C for 1 h and 20 minutes. The reaction mixture was allowed to cool to room temperature and diluted with EtOAc (8.00 mL), water (2.00 mL) and 2 mol/L hydrochloric acid solution (2.00 mL). After the mixture was filtered through celite, the organic layer was washed with 1 mol/L hydrochloric acid solution and water, dried over Na_2_SO_4_, and concentrated under reduced pressure. The crude mixture was recrystallized from IPA to afford compound **31a** (191 mg, 0.512 mmol, yield 68% over 2 steps). ^1^H NMR (400 MHz, CDCl_3_) δ: 5.14 (2H, s), 7.32-7.35 (2H, dd, *J* = 8.0, 2.0 Hz), 7.43-7.49 (3H, m), 7.69 (1H, t, *J* = 2.0 Hz), 8.47 (1H, d, *J* = 2.0 Hz), 8.66 (1H, d, *J* = 2.0 Hz). LC/MS (ESI): m/z = 373.0 [M+H]^+^.

### 2-(3-(5-Chloropyridin-3-yl)-2,4-dioxo-5-phenyl-6-(6-oxa-2-azaspiro[3.4]octan-2-yl)-3,4-dihydropyrimidin-1(2*H*)-yl)acetonitrile (13)

A mixture of compound **31a** (82.0 mg, 0.220 mmol), 6-oxa-2-azaspiro[3.4]octane hydrochloric acid salt (39.4 mg, 0.264 mmol), and DIPEA (115 μL, 0.659 mmol) in DMA (0.5 mL) was stirred at 60°C for 1 h. The reaction mixture was cooled to room temperature and diluted with water (1.00 mL) and ethyl acetate. The organic layer was washed with 2 mol/L hydrochloric acid solution and water, and concentrated under reduced pressure. The residue was purified by silica gel column chromatography (CHCl_3_ : MeOH = 24 : 1) to afford compound **13** (58.5 mg, 0.13 mmol, yield 59%). ^1^H NMR (400 MHz, CDCl_3_) δ: 2.05 (2H, t, *J* = 7.0 Hz), 3.70 - 3.81 (4H, m), 3.85 (4H, s), 4.78 (2H, s), 7.28 (2H, dd, *J* = 8.0, 2.0 Hz), 7.36-7.41 (3H, m), 7.68 (1H, t, *J* = 2.0 Hz), 8.46 (1H, br s), 8.60 (1H, br s). ^13^C NMR (100 MHz, CDCl_3_) δ: 35.0, 37.7, 40.9, 66.6, 67.6, 76.8, 99.0, 114.7, 128.4, 128.5, 131.4, 131.8 (br s), 132.1 (br s), 132.2, 136.4, 147.5, 148.6, 151.2, 152.6, 161.6. LC/MS (ESI): m/z = 450.1 [M+H]^+^.

### 4-Chloro-5-(3-chlorophenyl)-2,6-dimethoxypyrimidine (28b)

To a mixed solution of a 1.55 mol/L *n*-butyllithium solution in hexane (8.87 mL, 13.8 mmol) and THF (5.00 mL), a THF (6.50 mL) solution of 6-chloro-2,4-dimethoxypyrimidine (**26**) (2.00 g, 13.17 mmol) was added dropwise at -78°C over 15 minutes. The mixture was stirred at -78°C for 1 h. A 2.00 mol/L zinc chloride 2-methyltetrahydrofuran solution (7.16 mL, 14.3 mmol) was added dropwise over 5 minutes. The mixture was stirred at room temperature for 2 h. 1-Bromo-3-chlorobenzene (**27b**) (1.54 mL, 13.2 mmol) and tetrakis(triphenylphosphine)palladium (0.662 g, 0.573 mmol) were added, and the mixture was stirred at 80°C for 1.5 h. The reaction solution was cooled to room temperature, water (14.0 mL) and 2.00 mol/L hydrochloric acid (6.00 mL) were added thereto, and the mixture was extracted with EtOAc. The organic layer was concentrated under reduced pressure, and IPA (10.0 mL) was added to the obtained residue. The precipitate was collected by filtration and washed with IPA to afford compound **28b** (1.89 g, 6.63 mmol). ^1^H NMR (400 MHz, CDCl_3_) δ: 3.95 (3H, s), 4.05 (3H, s), 7.17-7.20 (1H, m), 7.30 (1H, m), 7.36-7.38 (2H, m). LC/MS (ESI): m/z = 285.0 [M+H]^+^.

### 6-Chloro-5-phenylpyrimidine-2,4(1*H,3H*)-dione (29b)

Acetic acid (4.30 mL) and concentrated hydrochloric acid (4.3 mL) were added to compound **28b** (1.89 g, 6.63 mmol), and the mixture was stirred at 110°C for 5 h. The reaction solution was cooled to room temperature, and then water (12.0 mL) was added thereto. The precipitate was collected by filtration and washed with water to afford compound **29b** (1.66 g, 6.46 mmol). ^1^H NMR (400 MHz, DMSO-*d*_6_) δ: 7.25-7.26 (1H, m), 7.35 (1H, s), 7.40-7.44 (2H, m), 11.55 (1H, br s), 12.20 (1H, br s). LC/MS (ESI): m/z = 256.9 [M+H]^+^.

### 6-Chloro-5-(3-chlorophenyl)-3-(5-chloropyridin-3-yl)pyrimidine-2,4(1*H*,3*H*)-dione (30b)

A mixture of compound **29b** (600 mg, 2.33 mmol), 3-bromo-5-chloropyridine (898 mg, 4.67 mmol), copper iodide (445 mg, 2.33 mmol), potassium carbonate (645 mg, 4.67 mmol) in NMP (5.00 mL) was stirred at 110°C, and then a mixture of *N*,*N’*-dimethylethylenediamine (502 μL, 4.67 mmol) and water (126 μL, 7.00 mmol) in NMP (0.200 mL) and was added dropwise. After the reaction mixture was stirred at 110°C for 3 h, the mixture was allowed to cool in a water bath, and diluted with water (10 mL), 2 mol/L citric acid aq. (2 mL). The precipitate was collected by filteration and washed with IPA. The obtained solid was dried under reduced pressure to afford a crude product of compound **30b** (970 mg). ^1^H NMR (400 MHz, DMSO-*d*_6_) δ: 7.31 (1H, br d, *J* = 6.4 Hz), 7.39 (1H, br s), 7.44-7.49 (2H, m), 8.09 (1H, s), 8.93 (2H, br s). LC/MS (ESI): m/z = 367.9 [M+H]^+^.

### 2-(6-Chloro-5-(3-chlorophenyl)-3-(5-chloropyridin-3-yl)-2,4-dioxo-3,4-dihydropyrimidin-1(2*H*)-yl)acetonitrile (31b)

A mixture of compound **30b** (500 mg, 1.09 mmol, 80 %purity), DIPEA(284 μL, 1.63 mmol), and 2-bromoacetonitrile (113 μL, 1.63 mmol) in DMF (2.00 mL) was then stirred at 60°C for 1 h. The reaction mixture was allowed to cool to room temperature and diluted with EtOAc (8.00 ml), water (2.00 mL) and 2 mol/L hydrochloric acid solution (2.00 mL). After the mixture was filtered through a short pad of Celite eluting with EtOAc, the organic layer was washed with 1 mol/L hydrochloric acid solution and water, dried over Na_2_SO_4_, and concentrated under reduced pressure. The crude mixture was recrystallized from IPA to afford compound **31b** (294 mg, 0.721 mmol, yield 66%, 2 steps from **29b**). ^1^H NMR (400 MHz, CDCl_3_) δ: 5.13 (2H, s), 7.23 (1H, ddd, *J* = 6.4, 2.0, 1.6 Hz), 7.35 (1H, br s), 7.38-7.43 (2H, m), 7.68 (1H, t, *J* = 2.0 Hz), 8.46 (1H, br s), 8.67 (1H, br s). LC/MS (ESI): m/z = 406.9 [M+H]^+^.

### 2-(5-(3-Chlorophenyl)-3-(5-chloropyridin-3-yl)-2,4-dioxo-6-(6-oxa-2-azaspiro[3.4]octan-2-yl)-3,4-dihydropyrimidin-1(2*H*)-yl)acetonitrile (14)

A mixture of compound **31b** (90.0 mg, 0.220 mmol), 6-oxa-2-azaspiro[3.4]octane hydrochloric acid salt (39.6 mg, 0.265 mmol), and DIPEA (116 μL, 0.662 mmol) in DMA (0.50 mL) was stirred at 60°C for 1.5 h. The reaction mixture was cooled to room temperature and diluted with water (1 mL) and EtOAc. The organic layer was washed with 2.0 mol/L hydrochloric acid solution and water, and concentrated under reduced pressure. The residue was purified by silica gel column chromatography (CHCl_3_ : MeOH = 24 : 1) to afford compound **14** (92.8 mg, 0.192 mmol, yield 87%). ^1^H NMR (400 MHz, CDCl_3_) δ: 2.09 (2H, t, *J* = 7.0 Hz), 3.72 - 3.85 (4H, m), 3.90 (4H, s), 4.76 (2H, *J* = 6.2 Hz, br d), 7.18 (1H, td, *J* = 1.8, 6.4 Hz), 7.29-7.36 (3H, m), 7.67 (1H, t, *J* = 2.0 Hz), 8.45 (1H, d, *J* = 2.0 Hz), 8.61 (1H, d, *J* = 2.0 Hz). ^13^C NMR (100 MHz, CDCl_3_) δ: 35.1, 37.8, 41.0, 67.0, 67.6, 77.4, 97.3, 114.5, 128.6, 129.5, 130.5, 131.9 (br s), 131.9 (br s), 132.2, 133.4, 134.2, 136.4, 147.4, 148.8, 151.0, 153.1, 161.3. LC/MS (ESI): m/z = 484.1 [M+H]^+^.

### 6-Chloro-5-(3-chloro-4-fluorophenyl)pyrimidine-2,4(1*H*,3*H*)-dione (28c)

To a mixed solution of a 2.64 mol/L *n*-butyllithium solution in hexane (1.50 L, 3.96 mol) and THF (1.44 L), a THF solution of 6-chloro-2,4-dimethoxypyrimidine **26** (576 g, 3.30 mol) was added dropwise at -78°C over 1 h under N_2_. The mixture was stirred at -78°C for 1 h. A 1.9 mol/L zinc chloride 2-methyltetrahydrofuran solution (2.08 L, 3.96 mol) was added dropwise over 5 minutes. The mixture was stirred at room temperature for 2 h. 4-bromo-2-chloro-1-fluorobenzene (**27c**) (760 g, 3.63 mol) and tetrakis(triphenylphosphine)palladium (191 g, 165 mmol) were added, and the mixture was stirred at 80°C for 1.5 h. The reaction solution was cooled to room temperature, EtOAc (5.80 L), water (4.00 L) and 2 mol/L hydrochloric acid (1.70 L) were added thereto. The organic layer was washed with H_2_O and 10% NaCl aq., dried over MgSO_4_, and concentrated under reduced pressure. The residue was solidified from IPA to obtain **28c** as a white solid (626 g, 1.90 mol, 92wt%, yield 58%). ^1^H NMR (400 MHz, CDCl_3_) δ: 3.95 (3H, s), 4.05 (3H, s), 7.16-7.21 (2H, m), 7.35 (1H, dd, *J* = 7.2, 2.0 Hz). LC/MS (ESI): m/z = 302.9 [M+H]^+^.

### 6-Chloro-5-(3-chloro-4-fluorophenyl)pyrimidine-2,4(1*H*,3*H*)-dione (29c)

Acetic acid (2.60 L) and concentrated hydrochloric acid (2.60 L) were added to compound **28c** (1.00 kg, 3.12 mmol), and the mixture was stirred at 90°C for 3 h. The reaction solution was cooled to room temperature, and then water (5.20 L) was added thereto. The precipitate was collected by filtration and washed with water (18.0 L). The filter cake was air-dried to obtain compound **29c** (890 g, 2.83 mol, 88wt%, yield 90%). ^1^H NMR (DMSO-*d*_6_) δ: 7.31 (1H, ddd, *J* = 8.8, 4.9, 2.1 Hz), 45 (1H, dd, *J* = 8.8, 8.8 Hz), 7.53 (1H, dd, *J* = 7.3, 2.1 Hz), 11.6 (1H, s), 12.2 (1H, br s). LC/MS (ESI): m/z = 274.9 [M+H]^+^.

### 6-Chloro-5-(3-chloro-4-fluorophenyl)-3-(5-chloropyridin-3-yl)pyrimidine-2,4(1*H*,3*H*)-dione (30c)

The reaction was carried out in two paralleled batches: Compound **29c** (250 g, 0.909 mol), 3-bromo-5-chloropyridine (350 g, 1.82 mol), copper(I) iodide (173 g, 0.909 mol), NMP (2.30 L), potassium carbonate (251 g, 1.82 mol), *N,N’*-dimethylethane-1,2-diamine (160 g, 1.82 mol) and distilled water (49.1 ml, 2.73 mmol) were mixed, and the solution was stirred at 110°C under N_2_ for 3 h. The water (2.80 L) and citric acid monohydrate solution (458 g, 2.18 mol in 2.75 L water) were added to the reaction mixture under water cooling. The batches were combined and filtered. The filter cake was washed with water (11.0 L) and IPE (2.00 L), air-dried at room temperature, and then, dried under vacuum at 40 °C to afford compound **30c** (701 g, 1.55 mol, 86wt%, yield 86%). ^1^H NMR (400 MHz, DMSO-*d*_6_) δ: 7.34-7.37 (1H, m), 7.47 (1H, dd, *J* = 9.0, 9.0 Hz), 7.52- 54 (1H, m), 8.06 (1H, t, *J* = 2.1 Hz), 8.50-8.75 (2H, br m), 12.7-13.5 (1H, br s) . LC/MS (ESI): m/z = 385.8 [M+H]^+^.

### 2-(6-Chloro-5-(3-chloro-4-fluorophenyl)-3-(5-chloropyridin-3-yl)-2,4-dioxo-3,4-dihydropyrimidin-1(2*H*)-yl)acetonitrile (31c)

To a solution obtained by mixing compound **30c** (1.09 kg, 2.33 mmol, 82wt%), DIPEA (1.22 L, 6.99 mol), and DMF (9.00 L), 2-bromoacetonitrile (838 g, 6.99 mol) was added, and the solution was stirred at room temperature 14 h. EtOAc (27.0 L), water (27.0 L) and 2.00 mol/L hydrochloric acid solution (3.49 L) was added to the reaction mixture. The organic layer was washed with water, 10% NaCl aq., dried over Na_2_SO_4_, filtered and concentrated under reduced pressure. The obtained residue was filtrated through a short column of silica gel (1.80 kg) which was washed with EtOAc (9.00 L). The filtrate was evaporated in vacuo, and the residue was recrystallized from acetone and water twice to give compound **31c** (760 g, 1.69 mol, 95wt%, yield 73%) as a yellow-brown solid. ^1^H NMR (400 MHz, DMSO-*d*_6_ ) δ: 5.29 (2H, s), 7.36-7.40 (1H, m), 7.52-7.57 (2H, m), 8.08 (1H, m), 8.56 (1H, br s), 8.76 (1H, br s). LC/MS (ESI): m/z = 424.9 [M+H]^+^.

### 2-(5-(3-Chloro-4-fluorophenyl)-3-(5-chloropyridin-3-yl)-2,4-dioxo-6-(6-oxa-2-azaspiro[3.4]octan-2-yl)-3,4-dihydropyrimidin-1(2*H*)-yl)acetonitrile (15)

Compound **31c** (100 mg, 0.235 mmol), 6-oxa-2-azaspiro[3.4]octane hydrochloride (17.4 mg, 0.070 mmol), DIPEA (82.0 μL, 0.470 mmol), and DMF (2.00 mL) were mixed, and the solution was stirred at 60°C for 30 mins. Water and ethylacetate were added to the reaction solution, and the mixture was extracted with EtOAc. The organic layer was washed with water, dried over sodium sulfate, and filtered. The filtrate was concentrated, and EtOAc (0.20 mL), hexane (0.50 mL) were added. The obtained precipitate was collected by filtration and washed with hexane-diisopropyl ether (2 : 1). The solid obtained was purified by silica gel column chromatography (CHCl_3_ : MeOH = 10 : 0 to 9.6 : 0.4), the solvent was distilled off under reduced pressure, and compound **15** (91.0 mg, 0.18 mmol, yield 77%) was obtained. ^1^H NMR (DMSO-*d*_6_) δ: 2.04 (2H, t, *J* = 6.8 Hz), 3.62 (2H, t, *J* = 6.8 Hz), 3.68 (2H, s), 3.90 (4H, s), 4.85 (2H, s), 7.22 (1H, m), 7.34-7.49 (2H, m), 7.97 (1H, s), 8.47 (1H, s), 8.69 (1H, s). ^13^C NMR (DMSO-*d*_6_) δ: 36.2, 36.7, 39.9, 66.2, 66.7, 75.7, 92.3, 116.0 (d, *J* = 21.2 Hz), 116.4, 118.6 (d, *J* = 17.5 Hz), 130.4, 130.6 (d, *J* = 3.6 Hz), 132.8 (d, *J* = 7.3 Hz), 133.1, 133.8, 136.5, 147.6, 148.1, 151.1, 154.8, 156.1 (d, *J* = 246 Hz), 161.0. LC/MS (ESI): m/z = 502.1 [M+H]^+^.

### 2-(5-(3-Chloro-4-fluorophenyl)-3-(5-chloropyridin-3-yl)-6-(3,3-difluoroazetidin-1-yl)-2,4-dioxo-3,4-dihydropyrimidin-1(2*H*)-yl)acetonitrile (16)

Compound **31c** (50.0 mg, 0.117 mmol), 3,3-difluoroazetidine (16.4 mg, 0.176 mmol), DIPEA (40.9 μL, 0.235 mmol), and DMF (1.00 mL) were mixed, and the solution was stirred at 60°C for 2 h. Water and EtOAc were added to the reaction solution, and the mixture was extracted with EtOAc. The organic layer was washed with water, dried over Na_2_SO_4_, and filtered. The filtrate was concentrated, and EtOAc (0.05 mL), hexane (0.125 mL), and IPE (0.125 mL) were added. The obtained precipitate was collected by filtration and washed with IPE. The obtained solid was dried under reduced pressure to obtain compound **16** (18.7 mg, 0.039 mmol, yield 33%). ^1^H NMR (DMSO-*d*_6_) δ: 4.32 (4H, t, *J* = 12.4 Hz), 4.92 (2H, s), 7.23-7.29 (1H, m), 7.44 (1H, dd, *J* = 9.2, 8.8 Hz), 7.43 (1H, m), 7.97 (1H, s), 8.47 (1H, s), 8.71 (1H, s). ^13^C NMR (DMSO-*d*_6_) δ: 35.7, 66.4 (t, *J* = 27.7 Hz), 95.3, 114.4 (t, *J* = 266.1 Hz), 116.1, 116.5 (d, *J* = 21.1 Hz), 119.1 (d, *J* = 17.5 Hz), 129.8 (d, *J* = 3.6 Hz), 130.5, 132.7 (d, *J* = 8.3 Hz), 132.9, 133.8, 136.6, 147.9, 147.9, 151.0, 153.4 (m), 156.6 (d, *J* = 247 Hz), 161.1. LC/MS (ESI): m/z = 482.0 [M+H]^+^.

### [5-(3-Chloro-4-fluorophenyl)-3-(5-chloropyridin-3-yl)-6-(6,6-difluoro-2-azaspiro[3.3]heptan-2-yl)-2,4-dioxo-3,4-dihydropyrimidin-1(2*H*)-yl]acetonitrile (17)

Compound **31c** (1.29 kg, 3.04 mol), 6,6-difluoro-2-azaspiro[3.3]heptane trifluoroacetic acid salt (900 g, 3.64 mol), DIPEA (1.59 L, 9.12 mol), and DMF (12.9 L) were mixed, and the solution was stirred at 60°C for 2 h. Water (18.3 L) was added to the reaction solution, and the mixture was extracted with EtOAc (18.3 L). The organic layer was washed with water (18.3 L) and 10% NaCl aq. (18.3 L), treated with an activated carbon, dried over MgSO_4_, and concentrated under reduced pressure. The obtained residue was crystallized from EtOAc (2.30 L) to obtain white solid (1.59 kg, 2.58 mol, 85wt%, yield 85%) as an ethyl acetate solvate. The ethyl acetate solvate (70.0 g, 11.3 mmol) was suspended in EtOH (350 mL) and stirred at room temperature for 5 h. The suspension was filtrated to afford compound **17** (59.1 g, 11.3 mmol, 99%) as a white solid. Melting point: 261°C (differential scanning calorimetry). ^1^H NMR (400 MHz, DMSO-*d_6_*) δ: 2.79 (4H, t, *J* = 12.4 Hz), 4.04 (4H, s), 4.83 (2H, s), 7.19-7.23 (1H, m), 7.39 (1H, dd, *J* = 9.0, 9.0 Hz), 7.43 (1H, dd, *J* = 7.6, 2.4 Hz), 7.97 (1H, t, *J* = 2.0 Hz), 8.47 (1H, d, *J* = 2.0 Hz), 8.69 (1H, d, *J* = 2.4 Hz). ^13^C NMR (100 MHz, DMSO-*d*_6_) δ: 26.8 (t, *J* = 10.9 Hz), 36.2, 44.6 (t, *J* = 22.6 Hz), 66.9, 92.1, 116.0 (d, *J* = 17.6 Hz), 116.4, 118.6 (d, *J* = 20.4 Hz), 119.5 (t, *J* = 275.6 Hz), 130.4, 130.4-130.5 (m), 132.7 (d, *J* = 7.3 Hz), 133.1, 133.8, 136.6, 147.7, 148.1, 151.1, 154.8, 156.2 (d, *J* = 245.8 Hz), 161.0. HRMS-ESI (m/z): [M+H]^+^ calcd for C_23_H_17_Cl_2_F_3_N_5_O_2_: 522.0706; found: 522.0705.

### 2-(5-(3-Chloro-4-fluorophenyl)-3-(5-chloropyridin-3-yl)-6-(3-hydroxy-3-(trifluoromethyl)azetidin-1-yl)-2,4-dioxo-3,4-dihydropyrimidin-1(2*H*)-yl)acetonitrile (18)

Compound **31c** (180 mg, 0.423 mmol), 3-(trifluoromethyl)azetidin-3-ol hydrochloride (120 mg, 0.676 mmol), DIPEA (0.200 mL, 1.15 mmol), and DMF (2.00 mL) were mixed, and the solution was stirred at 60°C for 1 h 45 min and then 70°C for 1 h. Additional 3-(trifluoromethyl)azetidin-3-ol hydrochloride (11.0 mg, 0.062 mmol) was added and then the solution was stirred at 70°C for 20 min. Water was added to the reaction solution, and the mixture was extracted with EtOAc. The organic layer was washed with water, dried over MgSO_4_, and filtered. The residue obtained was purified by silica gel column chromatography (CHCl_3_ : MeOH = 10 : 0 to 6 : 4), and the solvent was distilled off under reduced pressure. IPE was added to the residue and the precipitate was collected by filtration and washed with IPE. The obtained solid was recrystallized from hexane-EtOAc and then the solid obtained was dried under reduced pressure to obtain compound **18** (90.0 mg, 0.170 mmol, yield 40%). ^1^H NMR (DMSO-*d*_6_) δ: 3.91-4.19 (4H, m), 4.87 (2H, s), 7.25 (1H, m), 7.42 (1H, s), 7.42-7.49 (2H, m), 8.01 (1H, s), 8.49 (1H, s), 8.70 (1H, s). ^13^C NMR (DMSO-*d*_6_) δ: 36.0, 63.1, 66.9 (q, *J* = 32.5 Hz), 93.7, 116.2, 116.3 (d, *J* = 20.5 Hz), 118.9 (d, *J* = 17.6 Hz), 122.9, 124.3 (q, *J* = 280.0 Hz), 130.5 (d, *J* = 3.7 Hz), 130.5, 132.8 (d, *J* = 7.3 Hz), 133.0, 133.9, 136.6, 147.8, 148.1, 150.9, 153.7, 156.5 (d, *J* = 246 Hz), 161.1. LC/MS (ESI): m/z = 529.9 [M+H]^+^.

### 3C-like Protease Inhibition Assay

SARS-CoV-2 3CL^pro^ inhibition studies were conducted as described previously.^24, 34^ Briefly, testing compounds at various concentrations were dispensed to 384-well plate (Corning 3702) by an ECHO 555 dispenser (Labcyte Inc.) or manual. Next, 5 or 7.5 μL of substrate (Dabcyl-KTSAVLQSGFRKME [Edans]-NH_2_, 3249-v, Peptide Institute, Inc.) in assay buffer (1 mM ethylenediaminetetraacetic acid (EDTA), 10 mM DL -dithiothreitol (DTT), 0.01% bovine serum albumin (BSA), and 20 mM Tris-HCl (pH 7.5) was dispensed using Multidrop Combi (Thermo Scientific) or manual. The final substrate concentration was 4 μM in reaction mixture. The reaction was initiated by adding 5 or 7.5 μL of 3CL^pro^ (R&D Systems, Inc.) in assay buffer and incubated at room temperature for 4 to 5 h. The final concentration of enzyme was 0.3 nM in reaction mixture. After incubation, the reaction was stopped by adding 45 μL of water solution containing 0.1% formic acid, 10% acetonitrile, and 0.05 μmol/L Internal Standard (IS) peptide (Dabcyl-KTSAVLeu [^13^C_6_,^15^N]-Q, custom-synthesized by Peptide Institute, Inc.). The reactions were analyzed with mass spectrometry (MS) using a RapidFire 365 high-throughput sampling robot (Agilent Technologies) connected 6495 Triple Quadrupole LC/MS (Agilent technologies) accurate mass quadrupole time-of-flight mass spectrometer. Peak areas were acquired and analyzed using a RapidFire Integrator (Agilent Technologies). Reaction product peak areas were acquired from m/z 499.27; ISpeak areas were acquired from m/z 502.78. IC_50_ values were determined by plotting the compound concentration versus inhibition and fitting data with a four-parameter logistical fit (Model 205, XLfit, IDBS).

### SPA-based Competitive Experiment

SARS-CoV-2 3CL protease (R&D systems) were attached to SPA beads by incubating 0.27 µM random biotinylated-SARS-CoV-2 3CL protease and 7 mg/ml streptavidin YSi SPA scintillation beads (Revvity) in assay buffer (10mM Tris pH7.5, 5 mM DTT, 0.5 mM EDTA, 0.005% Tween20). Thirty microliters of the 3CL protease-SPA beads complex and 20 µL of assay buffer containing 0.6 µM ^14^C-labeled **S-892216** were dispensed to 384 well-plate (greiner, 781093). For the non-specific binding control wells, thirty microliters of the 3CL protease-SPA beads complex and 20 µL of assay buffer containing 0.6 µM ^14^C -labeled and 18 µM non-labeled **S-892216** were dispensed. Then the plate was sealed and incubated until an equilibrium was reached. Dissociation of ^14^C-labeled **S-892216** were initiated by the addition of 5 µL of 72 µM of non-labeled **S-892216**. The equilibrium and dissociation were monitored using a Microbeta^2^ plate reader (Revvity). The binding values were recorded in corrected counts per minute (CCPM). Dissociation experiments were fitted to a one-phase exponential decay model using GraphPad Prism version 9 (GraphPad Software, Inc., San Diego, CA, USA).

### Cell Culture and Virus

HEK293T/ACE2-TMPRSS2 cells were obtained from GeneCopoeia (Rockville, MD, USA). VeroE6/TMPRSS2 cells were obtained from the Japanese Collection of Research Bioresources (JCRB) Cell Bank (Osaka, Japan). A549-Dual hACE2-TMPRSS2 cells were obtained from InvivoGen (San Diego, CA, USA). These cells were maintained in culture medium of Dulbecco’s modified Eagle medium (DMEM) (Thermo Fisher Scientific, Waltham, MA, USA) supplemented with 10% heat-inactivated FBS (Corning, Steuben County, NY, USA or SERANA, Pessin, Germany) and 1% penicillin-streptomycin (P/S, Thermo Fisher Scientific). Human nasal cavity MucilAir™ cells and MucilAir™ medium were purchased from Epithelix (Plan-les-Ouates, Switzerland).

The following SARS-CoV-2 were obtained from the National Institute of Infectious Diseases (Tokyo, Japan): hCoV-19/Japan/TY/WK-521/2020 (Ancestral, A), hCoV-19/Japan/QHN002/2020 (Alpha, B.1.1.7), hCoV-19/Japan/TY8-612/2021 (Beta, B.1.351), hCoV-19/Japan/TY7-501/2021 (Gamma, P.1), hCoV-19/Japan/TY11-927/2021 (Delta, B.1.617.2), hCoV-19/Japan/TY38-871/2021 (Omicron, BA.1.1), hCoV-19/Japan/TY40-385/2022 (Omicron, BA.2), hCoV-19/Japan/TY41-721/2022 (Omicron, BA.2.12.1), hCoV-19/Japan/TY41-716/2022 (Omicron, BA.2.75), hCoV-19/Japan/TY41-703/2022 (Omicron, BA.4.1), hCoV-19/Japan/TY41-763/2022 (Omicron, BA.4.6), hCoV-19/Japan/TY41-702/2022 (Omicron, BE.1/BA.5-like), hCoV-19/Japan/TY41-704/2022 (Omicron, BA.5.2.1), hCoV-19/Japan/TY41-820/2022 (Omicron, BF.7), hCoV-19/Japan/TY41-828/2022 (Omicron, BF.7.4.1), hCoV-19/Japan/TY41-796/2022 (Omicron, BQ.1.1), hCoV-19/Japan/TY41-832/2022 (Omicron, CH.1.1.11), hCoV-19/Japan/TY41-795/2022 (Omicron, XBB.1), hCoV-19/Japan/23-018/2022 (Omicron, XBB.1.5.19), hCoV-19/Japan/TY41-951/2023 (Omicron, XBB.1.9.1), hCoV-19/Japan/TY41-984/2023 (Omicron, XBB.1.16), hCoV-19/Japan/TY41-831/2022 (Omicron, XBF), and hCoV-19/Japan/TY41-686/2022 (Omicron, XE).

The following SARS-CoV-2 were obtained from BEI Resources (Manassas, VA, USA): hCoV-19/USA/MI-UM-10052670540/2023 (Omicron, BA.2.86 [NR-59638]).

The following SARS-CoV-2 were obtained from Repository of data and biospecimen of infectious disease (REBIND, Tokyo, Japan): hCoV-19/Japan/RB23-006-81/2023 (Omicron, EG.5.1), hCoV-19/Japan/RB24-001-48/2024 (Omicron, JN.1).

All SARS-CoV-2 strains were propagated in VeroE6/TMPRSS2 cells and infectious titers were determined based on the standard TCID_50_ in VeroE6/TMPRSS2 cells.

### Compound Screening of Virus Replication Inhibition Assay Using HEK293T/ACE2-TMPRSS2 Cells

Antiviral activity of compounds against SARS-CoV-2 using HEK293T/ACE2-TMPRSS2 cells was conducted as described previously.^35^ Briefly, the HEK293T/ACE2-TMPRSS2 cells were suspended in minimal essential medium (MEM) (Thermo Fisher Scientific) supplemented with 2% FBS and 1% P/S and seeded with the diluted compounds into 384-well plates (3.0 × 10^3^ cells/well). The cells were then infected with hCoV-19/Japan/TY/WK-521 strain at 600 TCID_50_/well and cultured for 3 days at 37°C with 5% CO_2_. Cell viability was assessed using a CellTiter-Glo 2.0 assay (Promega, Madison, WI, USA). The luminescent signal (Relative Light Unit) was measured using an EnSpire multiplate reader (PerkinElmer Japan Co., Ltd.), and the percent inhibition of CPE induced by SARS-CoV-2 was calculated. Cell-control wells were not infected nor treated with any test or reference substance. Virus-control wells were infected with virus but not treated with a test or reference substance.

The concentrations of substances achieving 50% inhibition (EC_50_) against SARS-CoV-2 replication were calculated using software XLfit 5.3.1.3 (fit model: 205).

### Cellular Antiviral Activity Using VeroE6/TMPRSS2 Cells

Antiviral activity of compounds against SARS-CoV-2 in VeroE6/TMPRSS2 cells was assessed by monitoring the cell viability as previously reported.^35^ Briefly, VeroE6/TMPRSS2 cells (1.5 × 10^4^/well) suspended in MEM supplemented with 2% FBS and 1% P/S were seeded into 96-well plates with diluted compounds in each well. Cells were infected with each SARS-CoV-2 at 30– 3,000 TCID_50_/well and cultured at 37 °C with 5% CO_2_ for 3 or 4 days. Cell viability was assessed using a CellTiter-Glo 2.0 assay (Promega). The CC_50_ was assessed in the absence of viruses after being cultured for 3 or 4 days. EC_50_ and CC_50_ values were determined by plotting the compound concentration versus inhibition and fitting data with a four-parameter logistical fit (Model 205, XLfit). The mean and SD values were calculated based on three independent experiments.

### Virus Replication Inhibition Assay Using Human Airway Epithelial Cells

Antiviral activity of compounds against SARS-CoV-2 in hAECs assessed by monitoring viral production as previously reported.^35^ Human nasal epithelial cells (MucilAir™; Epithelix Sàrl, Switzerland) were seeded at approximately 5.0 × 10^5^ cells/well into a 24-well Transwell plate and then infected with the SARS-CoV-2 Omicron BE.1/BA.5-like strain (hCoV-19/Japan/TY41-702/2022) or XBB.1.5.19 strain (hCoV-19/Japan/23-018/2022) at approximately 5,000 TCID_50_/well. The cells were incubated at 37°C in a 5% CO_2_ incubator for 2 h. After incubation, the cells were washed with MucilAir™ medium (Epithelix Sàrl) to remove unabsorbed viruses and transferred to a 24-well Transwell plate containing serially diluted compound solutions prepared in 700 μL of MucilAir™ culture medium. The infected cells were incubated at 37°C in 5% CO_2_. The cell culture fluids were collected at 2 days (BE.1/BA.5-like) and 3 days (XBB.1.5.19) post infection and subjected to viral titration using VeroE6/TMPRSS2 cells. The concentrations achieving EC_90_ against SARS-CoV-2 replication were calculated using the two-point method. The mean and SD values were calculated based on three independent experiments.

### *In vitro* Selection of S-892216 Resistance Mutation

To select SARS-CoV-2 with reduced susceptibility to **S-892216**, BE.1/BA.5-like strain was passaged in VeroE6/TMPRSS2 cells with **S-892216** concentrations ranging from 5.56 to 50.0 nmol/L. The cells were seeded at a density of 2.00 × 10^5^ cells/well in a 12-well plate and infected with SARS-CoV-2 (2.00 × 10^4^ TCID_50_/well) in MEM supplemented with 2% FBS and 1% P/S. After incubation, supernatant was removed, and cells were washed before adding fresh medium with or without **S-892216**. The initial passage concentrations of S-892216 were 5.56, 16.7, or 50.0 nmol/L. This subculture was repeated 10 times, and when CPE was observed at the fifth subculture, **S-892216** concentration was increased three-fold. The virus titer of passage 10 samples was determined, with control viruses passaged without **S-892216**. At the end of the culture period, the supernatant was collected, and genotypic analysis of passaged SARS-CoV-2 was performed. Sequencing of the 3CL^pro^ and its cleavage sites region in the passaged viruses was achieved using Sanger sequencing (Eurofins Genomics K.K., Tokyo, Japan) and GENETYX v14.1.0 software (Nihon Server, Tokyo, Japan).

### Drug Susceptibility Testing for Recombinant SARS-CoV-2

Recombinant reverse-genetics-derived (rg)SARS-CoV-2 based on ancestral strain (hCoV-19/ Japan/TY/WK-521/2020) with 3CL^pro^ mutations were generated using the circular polymerase extension reaction method.^36^ Antiviral assays were conducted as previously reported.^31^ Briefly, VeroE6/TMPRSS2 cells (1.5 × 10^4^/well) suspended in MEM supplemented with 2% FBS and 1% P/S were seeded into 96-well plates with diluted compounds in each well. Cells were infected with each SARS-CoV-2 at 1,000 TCID_50_/well to and cultured at 37 °C with 5% CO_2_ for 3 days. Cell viability was assessed using a CellTiter-Glo 2.0 assay. Fold change values were calculated for each strain relative mean EC_50_ to the wild-type virus for the compounds. EC_50_ values were determined by plotting the compound concentration versus inhibition and fitting data with a four-parameter logistical fit (Model 205, XLfit) and mean and SD values were calculated based on three independent experiments.

### *In Vivo* SARS-CoV-2 Infection and Treatment Studies

*In vivo* SARS-CoV-2 infection experiments were conducted in accordance with the guidelines of the Association for Assessment and Accreditation of Laboratory Animal Care (AAALAC). The animal study protocol was approved by the director of the institute based on the report of the Institutional Animal Care and Use Committee of Shionogi Research Laboratories (Approved No. S22152D).

Mouse *in vivo* SARS-CoV-2 infection studies were done at Shionogi Pharmaceutical Research Center (Osaka, Japan) as previously reported.^37^ Female BALB/cAJcl mice were purchased from CLEA Japan, Inc. (Tokyo, Japan) and used at 5 weeks of age. The mice were maintained and treated as previously detailed.^37^ Briefly, mice were intranasally inoculated with SARS-CoV-2 Gamma strain (hCoV-19/Japan/TY7-501/2021) (1×10^4^ TCID_50_/mouse) under anesthesia. 24 h after infection, the mice were orally administered **S-892216** (0.1, 0.3, 1, 3, or 10 mg/kg, n=5 per group), ensitrelvir fumaric acid (50 mg/kg [as a free form], n =5 per group), or vehicle (n = 5 per group) bid (every 12 h) for 2 days. 72 h postinfection, the mice were euthanized *via* cervical dislocation under anesthesia; their lungs were removed, and the viral titers in the lung homogenates were determined using VeroE6/TMPRSS2 cells. Viral titers are expressed as log_10_ TCID_50_/mL. No adverse effects were observed in rodents under the same exposure conditions as the maximum dose of **S-892216** administered to mice.

### PK Study in Infected Mice

PK experiments in infected mice were conducted in accordance with the guidelines provided by AAALAC and were approved by Institutional Animal Care and Use Committee of Shionogi Research Laboratories (Approved No. S22152D).

Mouse PK studies were done at Shionogi Pharmaceutical Research Center (Osaka, Japan). BALB/cAJcl mice were intranasally inoculated with SARS-CoV-2 Gamma strain (hCoV-19/Japan/TY7-501/2021) (10,000 TCID_50_/mouse) and orally administered with **S-892216** (0.03, 0.1, 0.3, 1, 3, 10, or 30 mg/kg) 24 h post infection. Blood was taken at 0.5, 1, 2, 4, 8, 12, and 24 h after dosing for 0.03 to 10 mg/kg, and 0.5, 1, 2, 4, 8, 12, 24, and 32 h after dosing for 30 mg/kg (n = 4 per group per time point), and plasma concentrations of **S-892216** were determined by LC/MS/MS. LC/MS/MS analysis was performed using Triple Quad 7500 (Sciex). The plasma concentrations after repeated oral administration were simulated using Phoenix WinNonlin (Certara, L.P., Version 8.3) by non-parametric analysis.

### Metabolic Stability Studies

Rat liver microsomes (pool of 5, male) were purchased from the Jackson Laboratory Japan, Inc. (Yokohama, Japan). Human liver microsomes (HLM, pool of 15, male and female) and human hepatocyte (female, 55 years, Lot. HC-5-23) were purchased from Sekisui XenoTech (Kansas City, KS). Rat, dog, and monkey hepatocytes were prepared in our laboratory. Metabolic stabilities of the test compounds in rat and human liver microsomes and rat, dog, monkey, and human hepatocytes were determined at 0.5 μM. Regarding liver microsomes, the assays were conducted as previously reported.^24^ Regarding hepatocytes, the compounds were incubated with 1.0×10^6^ cells/mL in suspension in the William’s Medium E (Thermo Fisher Scientific Inc.) at 37 °C. Incubations were initiated by adding the compounds and terminated by adding the organic solvent (MeCN/MeOH = 1:1) after 0, 1, and 2 h of incubation at 37 °C. The precipitation protein was removed by centrifugation. The supernatants were analyzed by liquid chromatography tandem mass spectrometry (LC/MS/MS). LC/MS/MS was performed using LCMS-8060 (Shimadzu Corporation, Kyoto). All incubations were conducted in duplicate, and the percentage of compound remaining at the end of the incubation was determined from the LC/MS/MS peak area ratio.

### PK Study in Rat

The rat PK study was conducted in accordance with the guidelines provided by AAALAC and approved by the director of the institute after reviewing the protocol by the Institutional Animal Care and Use Committee in terms of the 3R(Replacement/Reduction/Refinement) principles (Approved No. S20086C).

Rat PK studies were done at Shionogi Pharmaceutical Research Center (Osaka, Japan). Male Sprague−Dawley rats were purchased from The Jackson laboratory Japan Inc and used for 8 weeks. For oral administration, the dosing vehicle was dimethyl sulfoxide/0.5% methylcellulose (400 cP) = 1:4. The compound was orally administered at 1 μmol/5 mL/kg (n = 2) under nonfasted conditions. Blood samples (0.2 mL) were collected with 1 mL syringes containing anticoagulants (EDTA-2K and heparin) at 0.5, 1,2, 4, 8, and 24 h after dosing. For intravenous administration, compounds were formulated as solutions in dimethyl sulfoxide/propylene glycol (1:1, v/v) and intravenously administered *via* the tail vein at 1 μmol/mL/kg (n = 2) under isoflurane anesthesia under nonfasted conditions. Blood samples (0.2 mL) were collected with 1 mL syringes containing anticoagulants (EDTA-2K and heparin) at 5, 30, 60, 120, 240, and 360 min after dosing. Blood samples were centrifuged to obtain plasma samples, which were transferred to each tube and stored in a freezer until analysis. Plasma concentrations were determined by LC/MS/MS after protein precipitation with MeCN. LC/MS/MS analysis was performed using a SCIEX Triple Quad 6500+ or SCIEX Triple Quad 6500 (Sciex, Framingham, MA). PK parameters were calculated by non-compartmental analysis.

### PK Study in Dog and Monkey

PK experiments in dogs and monkeys were conducted in accordance with the guidelines provided by AAALAC. The animal study protocol was approved by the director of the institute after reviewing the protocol by the Institutional Animal Care and Use Committee in terms of the 3R (Replacement/Reduction/Refinement) principles (Approved No. S15015C and S15013D).

Dog and Monkey PK studies were performed at Shionogi Aburahi Research Center (Shiga, Japan). Male beagles were purchased from Marshall BioResources Japan Inc. Female cynomolgus monkeys were purchased from Shin Nippon Biomedical Laboratories, Ltd. or Hamri Co., Ltd. For intravenous administration, compounds were formulated as solutions in dimethyl acetamide/ethanol/20% HP-β-CD in carbonate buffer (pH 9.0) (2:3:5, by volume) and intravenously administered *via* a forelimb or tail vein at 0.1 mg/0.2 mL/kg (n = 2) under nonfasted conditions. Blood samples (0.2 mL) were collected with 1 mL syringes containing anticoagulants (EDTA-2K and heparin) at 5, 15, 30, 60, 120, 240, 480, and 1440 min after dosing. Blood samples were centrifuged to obtain plasma samples, which were transferred to each tube and stored in a freezer until analysis. Plasma concentrations were determined by LC/MS/MS after protein precipitation with MeCN. LC/MS/MS analysis was performed using a SCIEX Triple Quad 6500+ (Sciex, Framingham, MA). PK parameters were calculated by non-compartmental analysis.

### Serum Protein Binding

Human, rat, dog, and monkey serum protein binding was evaluated by equilibrium dialysis method except for the human, rat, and dog serum protein bindings of Compound **17** (**S-892216**) where the evaluation was performed by ultrafiltration method. The equilibrium dialysis was performed at 2-4 μM using HTDialysis (Gales Ferry, CT), and the detail of procedures were described in the previous report.^38^ Concentration of test compounds were determined by LC/MS/MS (SCIEX Triple Quad 6500+, Sciex). For the ultrafiltration, [^14^C]-**S-892216** was spiked in each serum at the concentration of 0.1, 1, and 10 μg/mL. The serum containing [^14^C]-**S-892216** was applied to an ultrafiltration devise (Centrifree, Merck millipore) and centrifuged. The radioactivity in the serum and filtrate samples were determined by Liquid scintillation counter (2500TR or 3110TR, PerkinElmer).

### Permeability Assessment Using LLC-PK1 Cells

LLC-PK1 cells (BD Biosciences) were used for permeability assessment. The culture and assay conditions were described in the previous reports.^39, 40^ The permeability coefficient of the apical-to-basolateral was treated as the P_app_ value. The P_app_ values were calculated by dividing the transport amount on the receiver (basolateral) side by the initial concentration on the donor (apical) side. Test compounds were determined by LC/MS/MS (SCIEX Triple Quad 6500, Sciex).

### Expression and Purification of SARS-CoV-2 3CL^pro^ Protein for X-ray Crystal Structure Analysis

The SARS-CoV-2 3CL^pro^ (1-306) containing an *N*-terminal 10-histidine tag followed by a thrombin cleavage site was cloned into pET15b vectors. The 3CL^pro^ construct was expressed and purified in the same manner as below. E. coli strain BL21 Star (DE3) (Thermo Fisher Scientific) was transformed by the expression plasmid and then precultivated in a LB medium containing 100 μg/mL ampicillin sodium salt. Six milliliters of preculture was inoculated into 600 mL of fresh TB medium supplemented by 100 μg/mL ampicillin sodium salt in a 2-L flask with baffles. After vigorous shaking at 37 °C, 1 mM IPTG was added for the induction when the optical density (OD)600 reached 1.0. After induction for 16 h at 16 °C, the cells were harvested by centrifugation.

Cells expressing SARS-CoV-2 3CLpro were resuspended and sonicated. The clarified lysate was subjected to HisTrap FF 5 mL (Cytiva) equilibrated with 20 mM Tris-HCl (pH 8.0), 300 mM NaCl, 1 mM DTT, and 20 mM imidazole, and the proteins were eluted with a linear concentration gradient of imidazole (20–500 mM). Fractions containing SARS-CoV-2 3CL^pro^ were collected and mixed with thrombin at 4 °C overnight to remove the *N*-terminus His-tag. Thrombin-treated SARS-CoV-2 3CL^pro^ was applied to HisTrap FF 5 mL (Cytiva) to remove proteins with uncleaved His-tags. The flow-through fraction was applied to a HiLoad 16/60 Superdex 200 prep grade (Cytiva) equilibrated with 20 mM HEPES (pH 7.5), 150 mM NaCl, and 1 mM DTT, and the fraction containing the major peak was collected.

### Co-crystallization of SARS-CoV-2 3CL^pro^ with Compounds 1, 4 and 17 (S-892216), Diffraction Data Collection, and Structure Determination

*N*-terminal His-tag-free SARS-CoV-2 3CL^pro^ protein (4.6 mg/mL) was incubated with 250-500 μM compound for 1 h at room temperature, and the complexes were crystallized by sitting-drop vapor diffusion at 20 °C. The compound **1** complex crystal was grown with buffer containing 0.02 M Sodium/potassium phosphate 0.1 M Bis-Tris propane 7.5 20 % w/v PEG 3350. The compound **4** complex crystal was grown with buffer containing 0.2 M Sodium chloride, 0.1 M HEPES pH 7.5, 25% w/v Polyethylene glycol 3,350. The compound **17** (**S-892216**) complex crystal was grown with buffer containing 0.2 M Lithium sulfate monohydrate, 20% w/v Polyethylene glycol 3,350.

X-ray diffraction data were collected using a Rigaku HyPix6000C detector mounted on a Rigaku FR-X rotating anode generator. Data were processed by CrysAlis Pro.^41^ The structures were determined by molecular replacement using MOLREP^42^ with the SARS-CoV-2 3CLpro-inhibitor complex (PDB code: 6LU7) as a search model.^43^ Iterative model-building cycles were performed with COOT^44^ and refined using REFMAC^45^. The data collection and structure refinement statistics are summarized in Table S8.

## ASSOCIATED CONTENT

Supporting figures and tables for crystal structures of 3CL^pro^ complex, *in vitro* antiviral activities, *in vitro* protease selectivity experiments, antiviral profiling and crystallography data collection and refinement statistics; synthesis of [^14^C]-**S-892216**; ^1^H NMR and ^13^C NMR spectra of synthetic compounds; HPLC traces of compounds; physicochemical properties and HRMS data of **17** (**S-892216**); supplementary reference (PDF)

## Author contributions

Conceptualization: Y.U., J.S., H. Shibayama, T.K., T.S., Y.T.; Methodology: R.D., S.K.; Formal Analysis: H.N., R.W.; Investigation: Y.U., S.U., S.K., H.N., K.H., K.N., J.S., H. Shibayama, A.H., K.K., M. Takamatsu, S.Y., Q.Z., M. Tanimura, R.D., Y.M., R.W., T.M.; Data Curation: Y.U., S.K., H.N., K.H.; Writing – Original Draft Preparation: Y.U., S.U., H.N., K.H., R.W., T.M.; Writing –Review & Editing: Y.U., S.U., S.K., H.N., K.H., R.W., T.M.; Visualization: Y.U., S.U., H.N., R.D., R.W., T.M.; Project Administration: Y.U., R.W., T.K., T.S., Y.T.; Supervision: H.Sawa, T.K., T.S., Y.T.; Funding Acquisition: H. Sawa, T.S.

## Competing interests

SHIONOGI has applied for a patent covering **4-18**. Y.U., S.U., S.K., H.N., K.H., K.N., J.S., H. Shibayama, A.H., K.K., M. Takamatsu, S.Y., Q.Z., M. Tanimura, R.D., Y.M., R.W., T.M., T.K., T.S., Y.T. are employees of SHIONOGI & Co., Ltd. S.U., H.N., J.S., M. Tanimura, R.D., Y.M., R.W., T.M., T.K., T.S., Y.T. are shareholders in SHIONOGI & Co., Ltd. H. Sawa is financially supported by the joint research fund from SHIONOGI & Co., Ltd.

The manuscript was written through contributions of all authors. All authors have given approval to the final version of the manuscript.

## Acknowledgments

We acknowledge the National Institute of Infectious Diseases (NIID), the Repository of Data and Biospecimen of Infectious Disease (REBIND) project, and BEI resources, Dr. Kouichi Morita (Nagasaki University), and Dr. Bart Haagmans (Erasmus University Medical Center) for providing the SARS-CoV-2 strains, SARS-CoV, and MERS-CoV, respectively. We are also grateful to Osaka University, which through a Material Transfer Agreement provided the technology used to create the recombinant SARS-CoV-2 used in this study. We would like to thank all our colleagues who participated in the COVID-19 antiviral program at SHIONOGI. We are also grateful to Shionogi Technoadvance Research Co., Ltd. for the compound supplies and technical support in the pharmacological studies. We would like to thank Editage (www.editage.jp) for English language editing.

## Table of Contents graphic

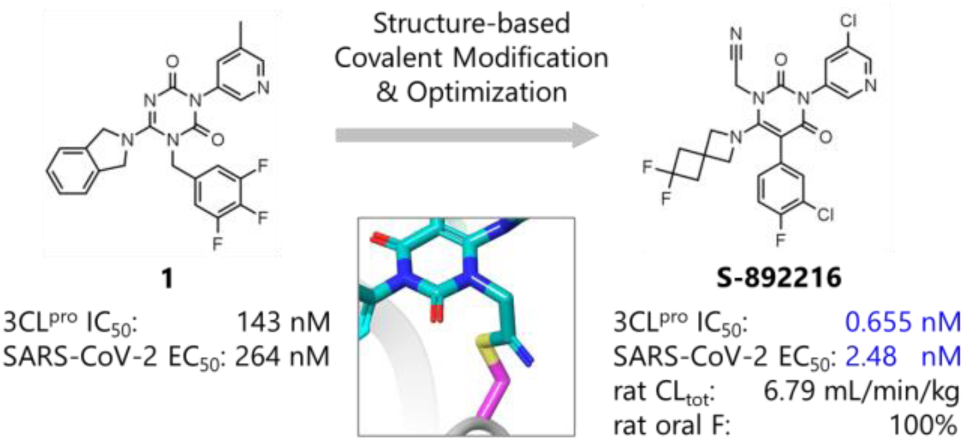

