## Supporting information for "Discovery of The Clinical Candidate S-892216: A Second-Generation of SARS-CoV-2 3CL Protease Inhibitor for Treating COVID-19"

**Table of contents**


- Figure S1. Crystal structure of 3CL$^{pro}$ in complex with compound **1**......S3
- Figure S2. **S-892216** resistant SARS-CoV-2 isolation study......S4
- Table S1. Selectivity of **S-892216** against human and HIV protease......S5
- Table S2. Antiviral activity of **S-892216** against various SARS-CoV-2 variants......S5
- Table S3. A list of 3CL$^{pro}$ residues that directly contact or are located within 5Å of **S-892216**......S6
- Table S4. Antiviral activity of **S-892216**, ensitrelvir, and nirmatrelvir against SARS-CoV-2 using VeroE6/TMPRSS2 cells with P-gp inhibitor and A549/ACE2-TMPRSS2 cells......S7
- Table S5. Potency shift of antiviral activity of **S-892216** and ensitrelvir in the presence of human or mouse serum......S7
- Table S6. Antiviral activity of **S-892216**, ensitrelvir, and nirmatrelvir against beta-coronavirus and alpha-coronavirus......S7
- Table S7. Drug susceptibility of rgSARS-CoV-2 with 3CL$^{pro}$ mutations associated with reduced susceptibility to ensitrelvir and nirmatrelvir......S8
- Table S8. Diffraction data and refinement statistics for SARS-CoV-2 3CL$^{pro}$ complexed with compound **1**, **4** and **17** (**S-892216**) ......S9
- Experimental Procedures for *in vitro* pharmacological assays ......S10
- Synthesis of [$^{14}$C]-**S-892216**......S12
- $^{1}$H-NMR and $^{13}$C-NMR spectra for synthesized compounds......S16
- HPLC chromatogram of compound **17** (**S-892216**)......S34
- HPLC chromatogram of compound **18**......S35
- Physicochemical Properties of **17** (**S-892216**) ......S36
- HRMS Spectra for compound **17** (**S-892216**)......S37
- References......S38

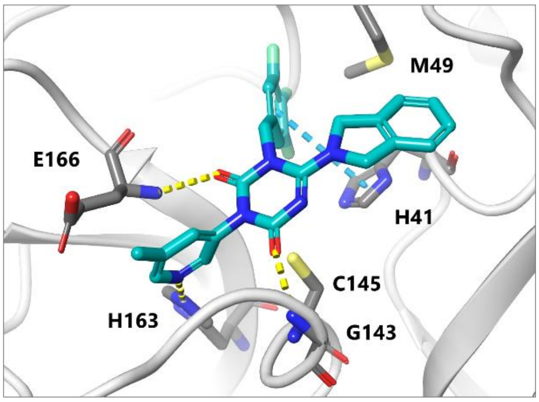


**Figure S1.** Crystal structure of 3CL$^{pro}$ in complex with compound **1**. The two carbonyl groups interacted with G143 and E166. The 3-methylpyridine group in S1 pocket formed a hydrogen bond with H163.

**A**

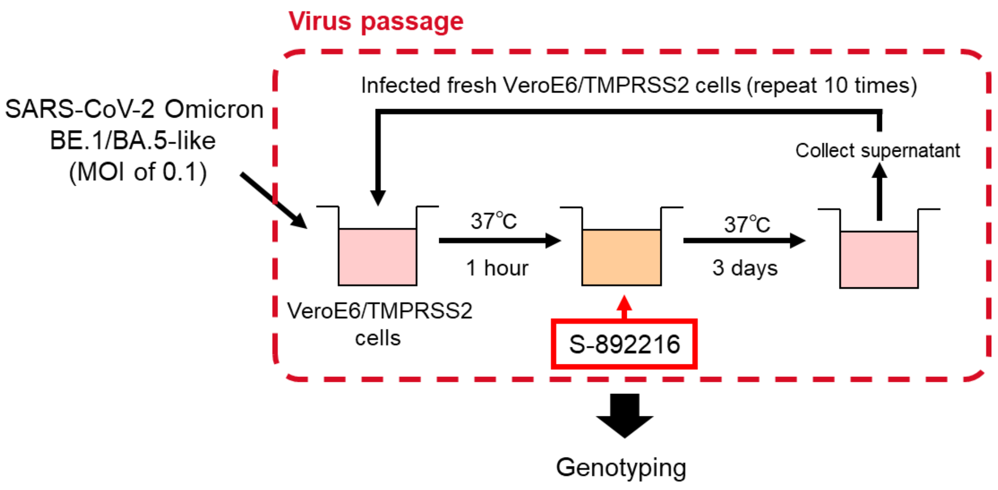


**B**

| **S-892216** (nM) | | No. | Amino acid substitutions |
|---|---|---|---|
| Passage 1-5 | Passage 6-10 | | in 3CL$^{pro}$ region |
| 0 | 0 | 1 | - |
| | | 2 | - |
| | | 3 | - |
| 5.56 | 5.56 | 1 | P252L |
| | | 2 | L50F, P252L |
| | | 3 | P252L |
| 5.56 | 16.7 | 1 | L50F, P252L |
| | | 2 | L50F, P252L |
| | | 3 | L50F, P252L |
| 16.7 | 16.7 | 1 | M49K, N221K, P252L |
| | | 2 | M49K, P252L |
| | | 3 | D48E, L50F, P252L |
| 16.7 | 50.0 | 1 | M49K, P252L |
| | | 2 | M49K, P252L |
| | | 3 | D48E, L50F, P252L |
| 50.0 | 50.0 | 1 | No virus |
| | | 2 | No virus |
| | | 3 | No virus |

**Figure S2. S-892216** resistant SARS-CoV-2 isolation study. (A) Passaging scheme for *in vitro* **S-892216** resistant isolation study. VeroE6/TMPRSS2 cells were infected with SARS-CoV-2 Omicron BE.1/BA.5-like strain and passaged to fresh VeroE6/TMPRSS2 cells every 3 days with **S-892216** for three independent wells. This subculture was repeated 10 times, and when CPE was observed at the fifth subculture, **S-892216** concentration was increased three-fold. (B) Mutations detected in 3CL$^{pro}$ during the *in vitro* passage study using **S-892216**.

**Table S1**. Selectivity of **S-892216** against human and HIV protease

| Protease | $IC_{50}$ (nM) |
|---|---|
| Human Caspase 2 | >10,000 |
| Human Chymotrypsin | >10,000 |
| Human Cathepsin B/D/G/L | >10,000 |
| Human Thrombin | >10,000 |
| HIV-1 Protease | >10,000 |

**Table S2.** Antiviral activity of **S-892216** against various SARS-CoV-2 variants in a cytopathic effect (CPE) inhibition assay using VeroE6/TMPRSS2 cells

| SARS-CoV-2 strains | WHO Label | Pango Lineage | $EC_{50}$ (nM) | |
|---|---|---|---|---|
| | | | **S-892216** | Ensitrelvir |
| hCoV-19/Japan/TY/WK-521/2020 | Ancestral | A | 8.77 ± 1.92 | 345 ± 102 |
| hCoV-19/Japan/QHN002/2020 | Alpha | B.1.1.7 | 3.85 ± 1.41 | 316 ± 38 |
| hCoV-19/Japan/TY8-612/2021 | Beta | B.1.351 | 9.64 ± 1.20 | 393 ± 109 |
| hCoV-19/Japan/TY7-501/2021 | Gamma | P.1 | 8.91 ± 1.62 | 622 ± 113 |
| hCoV-19/Japan/TY11-927/2021 | Delta | B.1.617.2 | 4.91 ± 1.29 | 338 ± 46 |
| hCoV-19/Japan/TY38-871/2021 | Omicron | BA.1.1 | 2.63 ± 0.18 | 136 ± 16 |
| hCoV-19/Japan/TY40-385/2022 | Omicron | BA.2 | 7.38 ± 3.82 | 257 ± 133 |
| hCoV-19/Japan/TY41-721/2022 | Omicron | BA.2.12.1 | 2.27 ± 0.26 | 198 ± 75 |
| hCoV-19/Japan/TY41-716/2022 | Omicron | BA.2.75 | 8.44 ± 1.25 | 217 ± 80 |
| hCoV-19/Japan/TY41-703/2022 | Omicron | BA.4.1 | 2.45 ± 0.40 | 178 ± 60 |
| hCoV-19/Japan/TY41-763/2022 | Omicron | BA.4.6 | 6.91 ± 1.21 | 302 ± 74 |
| hCoV-19/Japan/TY41-704/2022 | Omicron | BA.5.2.1 | 6.80 ± 1.38 | 366 ± 18 |
| hCoV-19/Japan/TY41-702/2022 | Omicron | BE.1/BA.5-like | 6.54 ± 3.49 | 259 ± 66 |
| hCoV-19/Japan/TY41-820/2022 | Omicron | BF.7 | 9.49 ± 0.56 | 512 ± 74 |
| hCoV-19/Japan/TY41-828/2022 | Omicron | BF.7.4.1 | 7.39 ± 1.97 | 552 ± 81 |
| hCoV-19/Japan/TY41-796/2022 | Omicron | BQ.1.1 | 11.0 ± 1.7 | 531 ± 84 |
| hCoV-19/Japan/TY41-832/2022 | Omicron | CH.1.1.11 | 5.69 ± 2.44 | 375 ± 91 |
| hCoV-19/Japan/TY41-795/2022 | Omicron | XBB.1 | 5.11 ± 2.15 | 259 ± 41 |
| hCoV-19/Japan/23-018/2022 | Omicron | XBB.1.5.19 | 7.40 ± 1.08 | 570 ± 74 |
| hCoV-19/Japan/TY41-951/2023 | Omicron | XBB.1.9.1 | 12.5 ± 0.9 | 986 ± 111 |
| hCoV-19/Japan/TY41-984/2023 | Omicron | XBB.1.16 | 7.06 ± 2.00 | 331 ± 26 |
| hCoV-19/Japan/TY41-831/2022 | Omicron | XBF | 6.27 ± 3.30 | 289 ± 15 |
| hCoV-19/Japan/TY41-686/2022 | Omicron | XE | 7.10 ± 2.59 | 209 ± 88 |

Data are the means ± SD; n=3 biological replicates. Beta variant carries amino acid substitution of K90R and Omicron variants carry amino acid substitution of P132H in $3CL^{pro}$.

**Table S3** A list of $3CL^{pro}$ residues that directly contact or are located within 5Å of **S-892216**

| $3CL^{pro}$ residue within 5Å of **S-892216** | Interaction between $3CL^{pro}$ and **S-892216** |
|---|---|
| T25 | - |
| T26 | - |
| L27 | - |
| H41 | Edge to face π-π interaction between side chain |
| C44 | - |
| T45 | - |
| S46 | - |
| M49 | - |
| Y54 | - |
| F140 | - |
| L141 | - |
| N142 | - |
| G143 | Hydrogen bond with main chain NH donor |
| S144 | - |
| C145 | Covalent bond with side chain S atom |
| H163 | Hydrogen bond with side chain NH donor |
| H164 | - |
| M165 | - |
| E166 | Hydrogen bond to main chain NH donor<br>CH-π interaction between side chain β-carbon |
| H172 | - |
| V186 | - |
| D187 | - |
| R188 | - |
| Q189 | - |

**Table S4.** Antiviral activity of **S-892216**, ensitrelvir, and nirmatrelvir against SARS-CoV-2 in a cytopathic effect (CPE) inhibition assay using VeroE6/TMPRSS2 cells with P-gp inhibitor and A549/ACE2-TMPRSS2 cells

| | | $EC_{50}$ (nM) | | |
|---|---|---|---|---|
| Cells | SARS-CoV-2 | **S-892216** | Ensitrelvir | Nirmatrelvir |
| VeroE6/TMPRSS2 cells with P-gp inhibitor | Ancestral, A | 3.36 ± 0.42 | 94.9 ± 38.6 | 68.1 ± 16.2 |
| A549/ACE2-TMPRSS2 cells | Ancestral, A | 2.21 ± 0.63 | 73.1 ± 16.4 | 47.5 ± 12.3 |

Data are the means ± SD; n=3 biological replicates (n=12 biological replicates for **S-892216** in A549/ACE2-TMPRSS2 cells).

**Table S5.** Potency shift of antiviral activity of **S-892216** and ensitrelvir in the presence of human or mouse serum using VeroE6/TMPRSS2 cells

| | Potency shift (-fold) | |
|---|---|---|
| Serum | **S-892216** | Ensitrelvir |
| Human | 4.78 ± 1.05 | 14.9 ± 6.1 |
| Mouse | 11.7 ± 4.6 | 27.8 ± 10.9 |

Data are the means ± SD; n=3 biological replicates.

**Table S6.** Antiviral activity of **S-892216**, ensitrelvir, and nirmatrelvir against beta-coronavirus and alpha-coronavirus in a cytopathic effect (CPE) inhibition assay using VeroE6/TMPRSS2 cells and MRC5 cells

| | | | $EC_{50}$ (nM) | | |
|---|---|---|---|---|---|
| Cells | Virus | | **S-892216** | Ensitrelvir | Nirmatrelvir |
| VeroE6/TMPRSS2 cells | Beta-coronavirus | SARS-CoV | 6.61 ± 5.36 | 210 [S1] | NT |
| VeroE6/TMPRSS2 cells | | MERS-CoV | 57.3 ± 15.1 | 1400 [S1] | NT |
| MRC5 cells | | HCoV-OC43 | 92.0 ± 21.0 | 142 ± 9 | 58.8 ± 12.2 |
| MRC5 cells | Alpha-coronavirus | HCoV-229E | 3570 ± 510 | 5500 [S1] | 696 ± 210 |

Data are the means ± SD; n=3 biological replicates.

**Table S7.** Drug susceptibility of rgSARS-CoV-2 with $3CL^{pro}$ mutations associated with reduced susceptibility to ensitrelvir and nirmatrelvir

| **rgSARS-CoV-2 with $3CL^{pro}$ mutations** | Fold change* v.s. wild-type | | | |
|---|---|---|---|---|
| | **S-892216** | Ensitrelvir | Nirmatrelvir | Remdesivir |
| T25A | 0.513 | 14.4 [S2] | 1.06 [S2] | 1.31 [S2] |
| M49I | 0.137 | 5.25 [S2] | 0.248 [S2] | 0.731 [S2] |
| M49L | 0.420 | 33.0 [S2] | 0.833 [S2] | 0.996 [S2] |
| S144A | 0.697 | 8.24 [S2] | 1.40 [S2] | 1.49 [S2] |
| T21I+E166V | 0.276 | 2.76 [S2] | 36.0 [S2] | 0.876 [S2] |
| L50F+E166V | 0.405 | 2.81 [S2] | 43.4 [S2] | 0.672 [S2] |

*Calculated by dividing the mean $EC_{50}$ of each tested mutant by that of the cognate wild-type.

**Table S8.** Diffraction data and refinement statistics for SARS-CoV-2 3CL$^{pro}$ complexed with compound **1**, **4** and **17 (S-892216)**

| | 3CL$^{pro}$ -Compound **1** | 3CL$^{pro}$ -Compound **4** | 3CL$^{pro}$-Compound **17 (S-892216)** |
|---|---|---|---|
| **PDB code** | 9LVR | 9LVT | 9LVV |
| **Data collection** | | | |
| Space group | P212121 | P21 | I2 |
| Cell dimensions | | | |
| *a, b, c* (Å) | 67.83 101.25 104.18 | 56.01 99.05 58.76 | 44.46 53.73 114.31 |
| *α, β, γ* (° ) | 90.00 90.00 90.00 | 90.00 108.16 90.00 | 90.00 98.88 90.00 |
| Wavelength (Å) | 1.54178 | 1.54178 | 1.54178 |
| Resolution (Å) | 30.22-2.20 (2.27-2.20) | 28.42-1.90(1.94–1.90) | 27.70-1.90(1.95-1.90) |
| Completeness (%) | 99.9(100.0) | 99.6(99.1) | 99.6(97.8) |
| $R_{merge}$ (%) * | 17.1(51.3) | 5.1(33.8) | 3.9(18.4) |
| $I/σ(I)$ | 9.1(3.7) | 21.3(3.6) | 18.3(5.2) |
| **Refinement** | | | |
| Resolution (Å) | 30.23 – 2.20 | 28.44 - 1.90 | 27.72 - 1.90 |
| No. of reflections | 35092 | 45336 | 19970 |
| $R_{work}/R_{free}$ | 0.2226/0.2684 | 0.1952/0.2371 | 0.1912/0.2417 |
| *B factor* (Å$^2$) | | | |
| Protein | 26.4 | 18.9 | 18.0 |
| Ligand | 38.9 | 17.3 | 18.7 |
| Water | 21.2 | 22.0 | 21.5 |
| RMS deviations | | | |
| Bond length (Å) | 0.0089 | 0.0073 | 0.0085 |
| Bond angles (° ) | 1.6676 | 1.6470 | 1.7725 |
| Ramachandran plot (%) | | | |
| Favored | 94.4 | 96.9 | 95.2 |
| Allowed | 4.9 | 2.6 | 4.1 |
| Outliers | 0.7 | 0.5 | 0.7 |

Values in parentheses are for the highest resolution shell.

* $R_{merge} = \Sigma| I - \langle I \rangle |/\Sigma I$, where *I* is the observed intensity and $\langle I \rangle$ is the average intensity.

**Human and HIV Protease Enzyme Assay**

Selectivity tests against a variety of human and HIV protease activity were conducted by Eurofins Panlabs Discovery Services Taiwan, Ltd., on behalf of Shionogi Co. & Ltd. as per established protocols. **S-892216** was tested on a set of seven human proteases (caspase-2, chymotrypsin, cathepsin B/D/G/L, and thrombin) and HIV-1 protease at 10 μM.

**Cellular Antiviral Activity using VeroE6/TMPRSS2 cells with P-gp inhibitor**

Antiviral activity of compounds against SARS-CoV-2 in VeroE6/TMPSS2 cells with P-gp inhibitor was assessed by same method as **Cellular Antiviral Activity using VeroE6/TMPRSS2 cells**. 0.75 μmol/L CP-100356 monohydrochloride (Sigma-Aldrich) was added as a P-gp inhibitor. Cells were infected with SARS-CoV-2 ancestral strain (hCoV-19/Japan/TY/WK-521/2020) at 100 $TCID_{50}$/well and incubated for 3 days.

**Cellular Antiviral Activity using A549-Dual hACE2-TMPRSS2 cells**

Antiviral activity of compounds against SARS-CoV-2 in A549-Dual hACE2-TMPRSS2 cells was assessed by monitoring the cell viability. A549-Dual hACE2-TMPRSS2 cells ($1.5 \times 10^4$ cells/well) suspended in MEM supplemented with 2% FBS and 1% P/S were seeded into 96-well plates with diluted compounds in each well. Cells were infected with SARS-CoV-2 Delta strain (hCoV-19/Japan/TY11-927/2021) at 100 $TCID_{50}$/well and cultured at 37 °C with 5% $CO_2$ for 2 days. Cell viability was assessed using a CellTiter-Glo 2.0 assay (Promega). $EC_{50}$ values were determined by plotting the compound concentration versus inhibition and fitting data with a four-parameter logistical fit (Model 205, XLfit). The mean and SD values were calculated based on three (twelve for **S-892216**) independent experiments.

**Cellular Antiviral Activity in the Presence of Human and Mouse Serum**

The inhibitory effect of **S-892216** on SARS-CoV-2 replication in cultured cells in the presence of human and mouse serum was evaluated using viral replication inhibition assays as previously reported.[S3] Briefly, VeroE6/TMPRSS2 cells ($1.5 \times 10^4$ cells/well) were seeded on 96-well plates in MEM supplemental with 2% FBS and 1% P/S 1 day prior to infection. After the removal of supernatant, the cells were infected with 1000 $TCID_{50}$ SARS-CoV-2 Gamma strain (hCoV-19/Japan/TY7-501/2021) and incubated for about 1 hour at 37°C, 5% $CO_2$. The supernatant including unadsorbed viruses were removed, and then added fresh medium with diluted compound and human serum (Sigma) or mouse serum (BALB/cAJcl mice, CLEA Japan, Inc.) (final 12.5, 25, or 50%). The plates were incubated for 1 day at 37°C, 5% $CO_2$, followed by extraction of the viral RNA from the cell lysate using TRIzol LS (Thermo Fisher Scientific Inc.) and a Direct-zol-96 RNA Kit (Zymo Research, Irvine, CA, USA). The viral RNA was quantified by real-time PCR using EXPRESS One-Step SuperScript qRT-PCR kit (Thermo Fisher Scientific Inc.) on an Applied Biosystems QuantStudio 5 system (Thermo Fisher Scientific) with SARS-CoV-2-specific primers and probes.[S4] $EC_{90}$ values in the presence of each concentration of serum were calculated with two points sandwiching one-tenth reduction. $EC_{90}$ values extrapolated to 100% serum were calculated by linear regression using the $EC_{90}$ values of each serum concentration. Potency shifts were calculated by dividing $EC_{90}$ extrapolated to 100% seurm by $EC_{90}$ in the absence of human and mouse serum. The mean and SD values were calculated based on three independent experiments.

**Cellular Antiviral Activity Against Coronaviruses**

Antiviral activity of compounds against SARS-CoV, MERS-CoV, HCoV-OC43, and HCoV-229E was assessed by monitoring the cell viability. Antiviral activities against SARS-CoV and MERS-CoV were evaluated at Hokkaido University using VeroE6/TMPRSS2 cells as previously reported.[S1] Briefly, VeroE6/TMPRSS2 cells ($1.5 \times 10^4$ cells/well) suspended in MEM supplemented with 2% FBS were seeded into 96-well plates with diluted compounds in each well. Cells were infected with each SARS-CoV at 1000 $TCID_{50}$/well or MERS-CoV 2500 $TCID_{50}$/well and cultured at 37 °C with 5% $CO_2$ for 3 days. Cell viability was assessed via (3-[4,5-dimethyl-2-thiazolyl]-2,5-diphenyl-2H-tetrazolium bromide (MTT) assay (Nacalai Tesque, Japan). Antiviral activity against HCoV-OC43 and HCoV-229E was evaluated using MRC-5 cells. MRC-5 cells ($2.0 \times 10^4$ cells/well) suspended in MEM supplemented with 2% FBS and 1% P/S were seeded into 96-well plates and incubated at 37 °C with 5% $CO_2$ overnight. The next day, the cells were infected with HCoV-OC43 or HCoV-229E at 300 or 1,000 $TCID_{50}$/well, respectively, and incubated at 37 °C with 5% $CO_2$ for 1 h, followed by removal of the inoculum and addition of MEM supplemented with 2% FBS and 1% P/S with the diluted compounds. The plates were incubated at 37 °C with 5% $CO_2$ for 3 days. Cell viability was assessed using a CellTiter-Glo 2.0 assay and $EC_{50}$ values were determined by plotting the compound concentration versus inhibition and fitting data with a four-parameter logistical fit (Model 205, XLfit). The mean and SD values were calculated based on three independent experiments.

**Synthesis of [$^{14}$C]-S-892216**

Synthesis of **[$^{14}$C]-S-892216** via an alternative ring synthesis approach is summarised in Scheme S1. For the detail discussion of **S-892216** synthesis routes, see references[S5,S6].

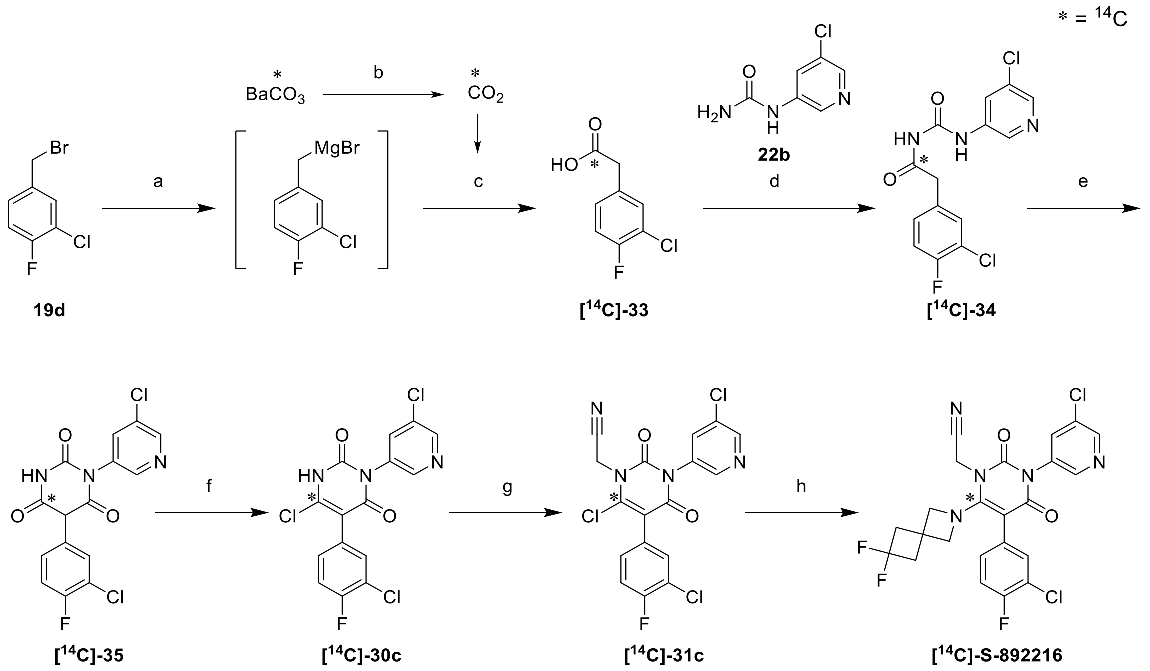


**Scheme S1.** Synthesis of **[$^{14}$C]-S-892216**.

(a) Mg, $I_2$, $Et_2O$, 35°C; (b) $HClO_4$; (c) -196°C to -78°C to rt; (d) **22b,** $T_3P$, $Et_3N$, DMA, rt; (e) CDI, DBU, THF, rt; (f) $POCl_3$, $H_2O$. 90°C; (g) 2-bromoacetonitrile, DIPEA, DMF, 40°C; (h) 6,6-difluoro-2-azaspiro[3.3]heptane trifluoroacetic acid salt, DIPEA, DMF, 60°C

**2-(3-Chloro-4-fluorophenyl)acetic-1-$^{14}$C acid ([$^{14}$C]-33)**

To a stirring suspension of Mg (55 mg, 1.98 mmol) in $Et_2O$ (2 mL) was added a piece of $I_2$ and dropwise **19d** (418 mg, 1.98 mmol) in $Et_2O$ (1 mL) and rinse out with $Et_2O$ (1 mL) at r.t.. The resulting suspension was stirred at 35°C for 1.5 hours. The resulting mixture was cooled with liquid nitrogen and introduced [$^{14}$C]-carbon dioxide generated from barium [$^{14}$C]-carbonate (299 mg, Total activity (TA) = 86.3 mCi, Specific activity (SA) = 56.8 mCi/mmol, 1.52 mmol) and 60% perchloric acid by vacuum line. After stirring at about -70°C for 10 min, the reaction mixture was warmed slowly to 0°C for 2 hours to complete the reaction. The reaction mixture was quenched with water (2 mL) at 0°C and acidified (pH = 1) with 2M HCl aq. (2.5 mL) and extracted with $Et_2O$. The organic layer was extracted with 1M $NaHCO_3$ aq. (6 mL). The aqueous layer was acidified (pH = 2) with 2M HCl aq. (4 mL) and extracted with $Et_2O$. The organic layer was washed with brine, dried over sodium sulfate, filtered and evaporated under reduced pressure. The crude **[$^{14}$C]-33** (278 mg, 1.46 mmol, as white crystals) was used in next reaction without further purification. HPLC purity 92%

**2-(3-Chloro-4-fluorophenyl)-*N*-((5-chloropyridin-3-yl)carbamoyl)acetamide-1-$^{14}$C ([$^{14}$C]-34)**

To a stirring suspension of **[$^{14}$C]-33** (Theoretical value: 286 mg, 1.52 mmol) and **22b** (286 mg, 1.67 mmol) in DMA (2 mL) was added $NEt_3$ (0.242 mL, 1.74 mmol). After 10 minutes, the mixture was clear solution. To the reaction mixture was added $T_3P$ (50% in AcOEt, 1.4 mL, 2.35 mmol) and stirred at 50°C for 2 hours. To the reaction mixture was added $T_3P$ (total 4 mL) every hour. To the reaction mixture was added $NEt_3$ (total 1.3 mL) every 30 minutes and stirred at 50°C for 2 hours. The reaction mixture was evaporated under reduced pressure to remove AcOEt. Then the reaction mixture was cooled 0°C, $H_2O$ (3.8 mL) was added and stirred at 0°C for 30 minutes. The precipitated solid were collected by filtration and washed with 50% MeCN/$H_2O$ (3 mL) to obtain **[$^{14}$C]-34** (432 mg, 1.26 mmol). HPLC purity 92%

**5-(3-Chloro-4-fluorophenyl)-1-(5-chloropyridin-3-yl)pyrimidine-2,4,6(1*H*,3*H*,5*H*)-trione-4-$^{14}$C ([$^{14}$C]-35)**

To a stirring suspension of **[$^{14}$C]-34** (432 mg, 1.26 mmol) in 2-Me-THF (4 mL) was added 1,1'-carbonyldiimidazole (328 mg, 2.02 mmol) and dropwise 1,8-diazabicyclo[5.4.0]-7-undecene (264 μL, 1.77 mmol) for 2 minutes. To a reaction mixture was added 2-Me-THF (1 mL) and stirred at r.t. for 1 h. The reaction mixture was poured into ice (ca 2 g), water 2 mL and 2M HCl aq. (4.5 mL), and rinsed out with 2-Me-THF (5 mL). The organic layer was separated. The aqueous layer was re-extracted twice with 2-Me-THF (10 mL). The organic layers were combined and evaporated in reduced pressure to ca.2.5g. To a stirring solution of the residue was added toluene (7 mL) and acetic acid, and cooled under ice-bath. The suspension was stirred at 0°C for 1 h. The precipitated solid were collected by filtration and washed with cold toluene (1.86 mL) to obtain **[$^{14}$C]-35** (381 mg, 1.04 mmol) as yellow solid. Mother liquor was purified by column chromatography ($CHCl_3$/MeOH : 19/1 to 3/2) to obtain **[$^{14}$C]-35** (60 mg, 0.164 mmol) as yellow solid. HPLC purity 99%

**6-Chloro-5-(3-chloro-4-fluorophenyl)-3-(5-chloropyridin-3-yl)pyrimidine-2,4(1*H*,3*H*)-dione-6-$^{14}$C ([$^{14}$C]-30c)**

To a combined starting material **[$^{14}$C]-35** (441 mg, 1.20 mmol) was added $POCl_3$ (3 mL) at r.t. and stirred at 50°C for 10 minutes. To a reaction mixture (yellow slurry) was added water (65 μL) and stirred at 90°C for 12 hours and allowed to stand overnight at r.t.. The reaction mixture was stirred again at 90°C for 12 hours 40 minutes and allowed to stand overnight at r.t. An ice-cooled mixture of 30% KOHaq. (120 mmol) and THF (15 mL) was added dropwise to reaction mixture over 1 h under ice bath cooling. To the reaction mixture was added THF (10 mL) and toluene (12.5 mL) and stirred at r.t. for 15 minutes and rinse out with THF:toluene=2:1 (10 mL) and water (3 mL). The organic layer was separated. The aqueous layer was re-extracted twice with THF:toluene=2:1 (45 mL). The organic layers were washed with 20% KCl aq. (35 mL), combined and evaporated in reduced pressure. To a stirring solution of the residue in DMA (7 mL) was added acetic acid (510 μL) and warmed at 50°C. To a solution was added dropwise water (4.64 mL) over 2 minutes under stirring. After 13 minutes, the oil bath was removed. The suspension was stirred at r.t for 20 minutes and 0°C for 1 h. The precipitated crystals were collected and washed with 70% DMA aq. (2 mL) and water (5 mL) to give **[$^{14}$C]-30c** (345.3 mg, 0.893 mmol, as white crystals). To the mother liquor was added water (2 mL) under ice bath cooling. The precipitated crystals were collected and washed with 70% DMA aq. (1 mL) and water (5 mL) to give **[$^{14}$C]-30c** (57.3 mg, 0.148 mmol, as brown crystals). HPLC purity 97%.

**2-(6-Chloro-5-(3-chloro-4-fluorophenyl)-3-(5-chloropyridin-3-yl)-2,4-dioxo-3,4-dihydropyrimidin-1(2*H*)-yl-6-$^{14}$C)acetonitrile ([$^{14}$C]-31c)**

To a stirring solution of **[$^{14}$C]-30c** (402.6 mg, 1.036 mmol) in DMF (4 mL) was added 2-bromoacetonitrile (207 µL, 3.11 mmol) and DIPEA (543 µL, 3.11 mmol). The resulting mixture was stirred at 40°C for 3 h. To a reaction mixture was added EtOAc, 2N-HCl and water (4 mL) under ice bath cooling. The organic layer was separated and washed twice with water (4 mL). The aqueous layer was re-extracted with EtOAc. The organic layers were combined and washed with brine, dried over sodium sulfate, filtered and evaporated under reduced pressure. The residue was purified by silica gel column chromatography (*n*-hexane/EtOAc = 4/1 to 2/1) to give crude product. The crude product was re-purified by amino silica gel column chromatography (*n*-hexane/EtOAc = 1/1) followed by trituation and filtlation in IPE to give **[$^{14}$C]-31c** (366.8 mg, 0.858 mmol, as pale beige powder). HPLC purity 98%.

**[$^{14}$C]-S-892216**

To a stirring solution of **[$^{14}$C]-31c** (366 mg, 0.856 mmol) in DMF (3.7 mL) was added 6,6-difluoro-2-azaspiro[3.3]heptane 2,2,2-trifluoroacetate (254 mg, 1.027 mmol) and DIPEA (179 µL, 1.025 mmol), and stirred at 60°C. After 2 hours 15 minutes, to a reaction mixture was added azaspiro[3.3]heptane 2,2,2-trifluoroacetate (66.5 mg, 0.269 mmol) and DIPEA (47 µL, 0.257 mmol) and stirred at 60°C for 2 h. To a reaction mixture was added DIPEA (74 µL, 0.428 mmol) and stirred at 60°C for 1.5 h. After cooling to r.t., EtOAc (12 mL) and water (12 mL) were added. The organic layer was separated and washed twice with water. The aqueous layer was re-extracted with EtOAc. The organic layers were combined and washed with brine, dried over sodium sulfate, filtered and evaporated under reduced pressure. The crude product was re-purified by amino silica gel column chromatography (EtOAc only). The obtained eluate was roughly evaporated and crystallized. To an obtained suspension was added dropwise IPE (1.5 mL). The supernatant was removed by decanting. The residue was triturated with IPE (1 mL) and the supernatant was removed by decanting. The residue was dried under reduced pressure to give **[$^{14}$C]-S-892216** (412 mg, 0.673 mmol). HPLC purity 98.7%

Analytical HPLC was performed on a YMC-Triart ($C_{18}$, 5 µm, 3.0 × 150 mm, mobile phase [A] = 0.1% $HCOONH_4$ / $H_2O$, [B] = 0.1% $HCOONH_4$ / MeCN, flow rate 0.4 mL/min, at 40°C) using a Shimadzu HPLC system equipped with a Radiomatic 625TR (PerkinElmer Inc.), Cocktail: ULTIMA-FLO AP, flow rate 1 mL/min.

Time-program for gradient elution

| Time (min) | Mobile phase B (%) |
|---|---|
| 0 | 50 |
| 20 | 85 |
| 21 | 85 |
| 21.01 | 50 |
| 25 | 50 |

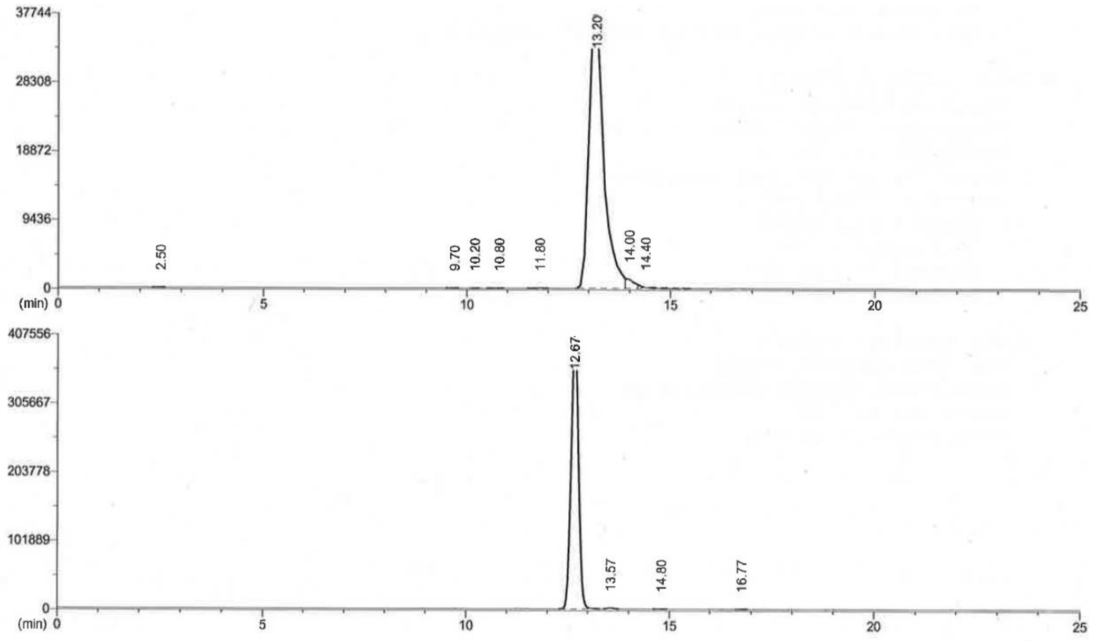


Channel 1: $^{14}C$ CPM

| peak# | Rt (min) | Area | % Pks |
|---|---|---|---|
| 1 | 2.50 | 19 | 0.01 |
| 2 | 9.70 | 9 | 0.01 |
| 3 | 10.20 | 9 | 0.01 |
| 4 | 10.80 | 28 | 0.02 |
| 5 | 11.80 | 56 | 0.03 |
| 6 | 13.20 | 162843 | 99.83 |
| 7 | 14.00 | 154 | 0.09 |
| 8 | 14.40 | 0 | 0 |
| Total peak area | | 161118 | |

Channel 2: UV

| peak# | Rt (min) | Area | % Pks |
|---|---|---|---|
| 1 | 12.67 | 5.25E+06 | 99.72 |
| 2 | 13.57 | 12910 | 0.25 |
| 3 | 14.80 | 1040 | 0.02 |
| 4 | 16.77 | 878 | 0.02 |
| Total peak area | | 5262702 | |

### $^{1}H$ NMR and $^{13}C$ NMR spectra for synthesized compounds

$^{1}H$ NMR spectra of **1** in $CDCl_3$

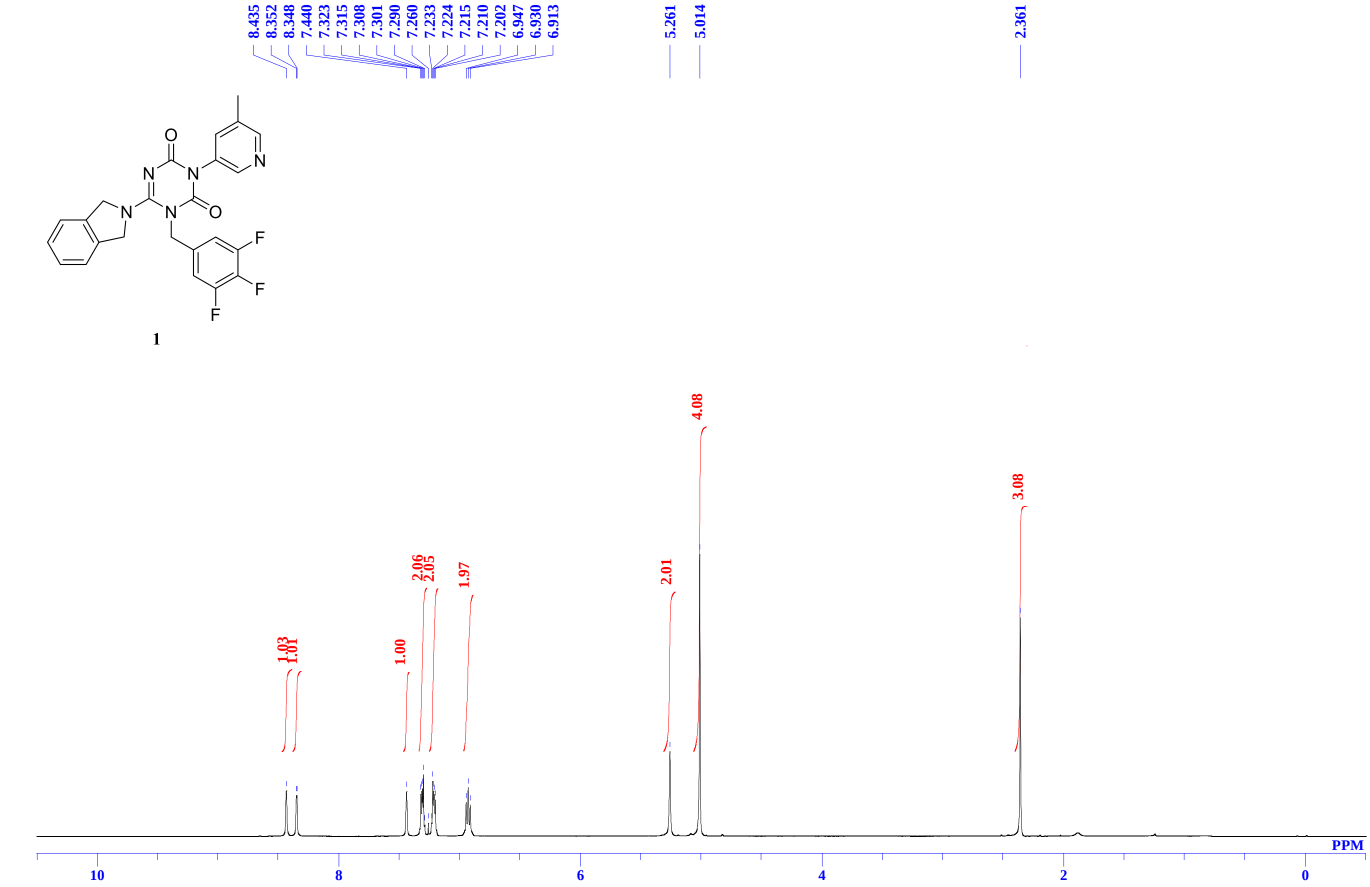


$^{13}C$ NMR spectra of **1** in $CDCl_3$

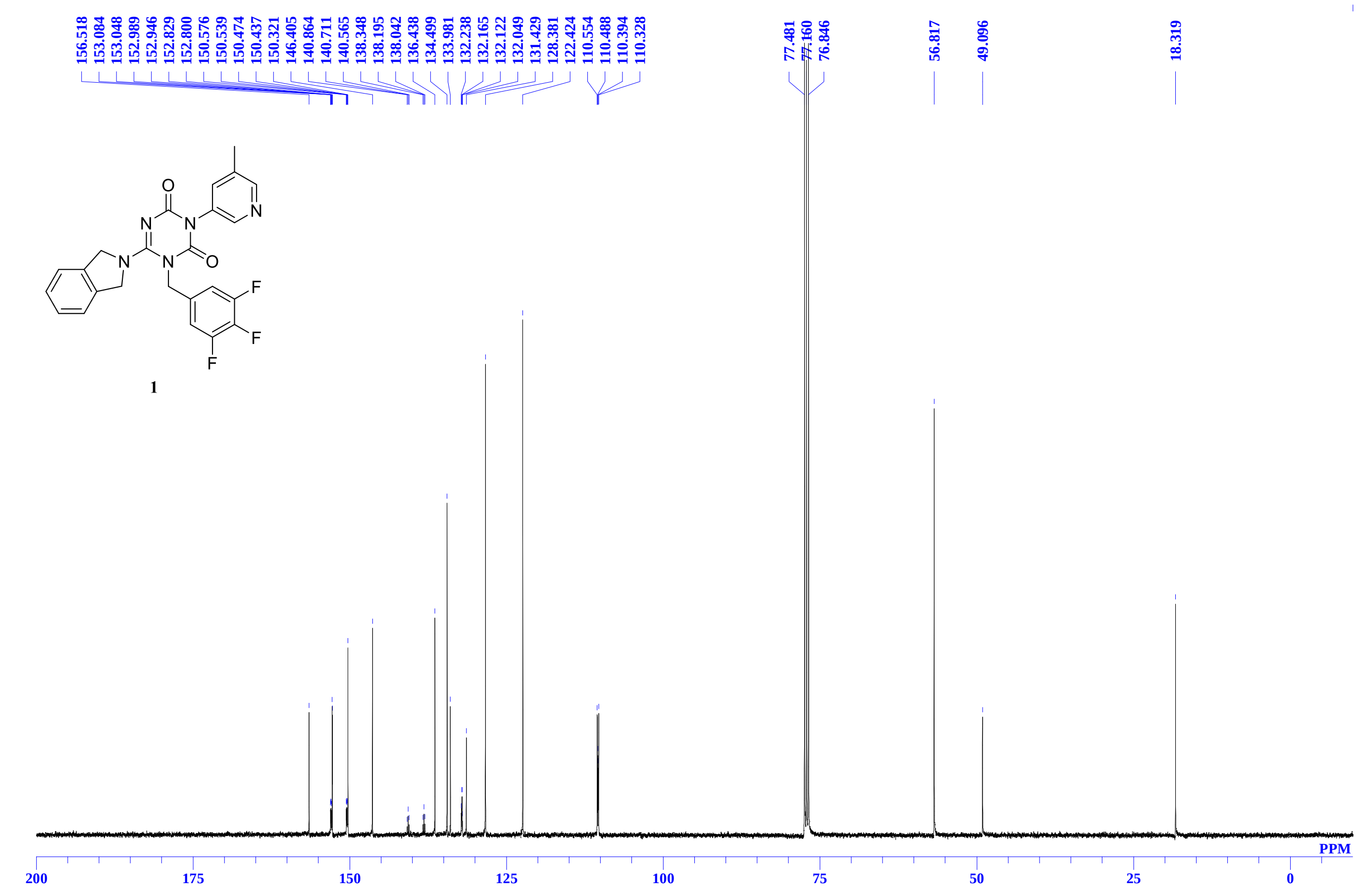

[1]H NMR spectra of **2** in DMSO-$d_6$

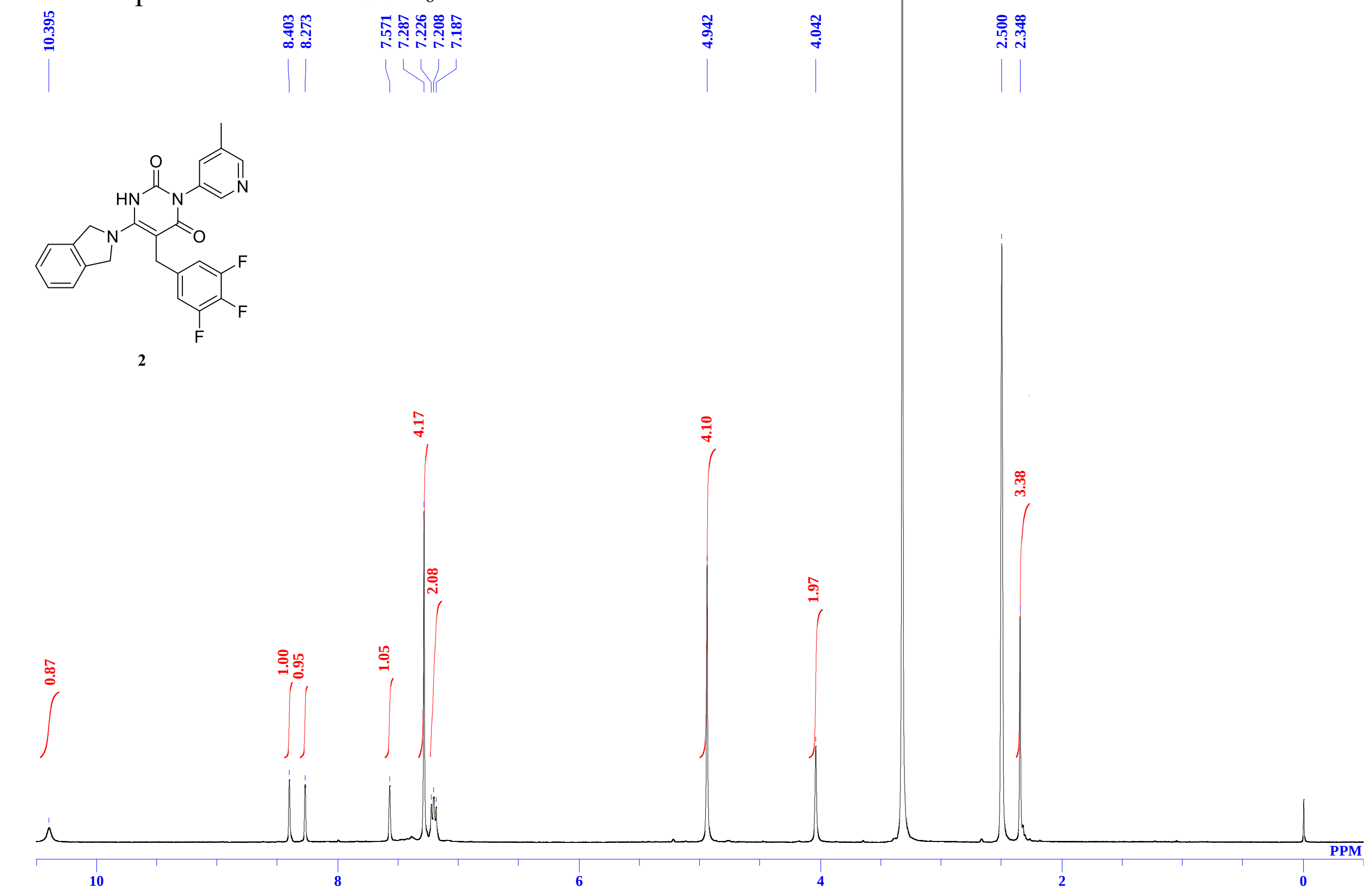


**13C** NMR spectra of **2** in DMSO-$d_6$

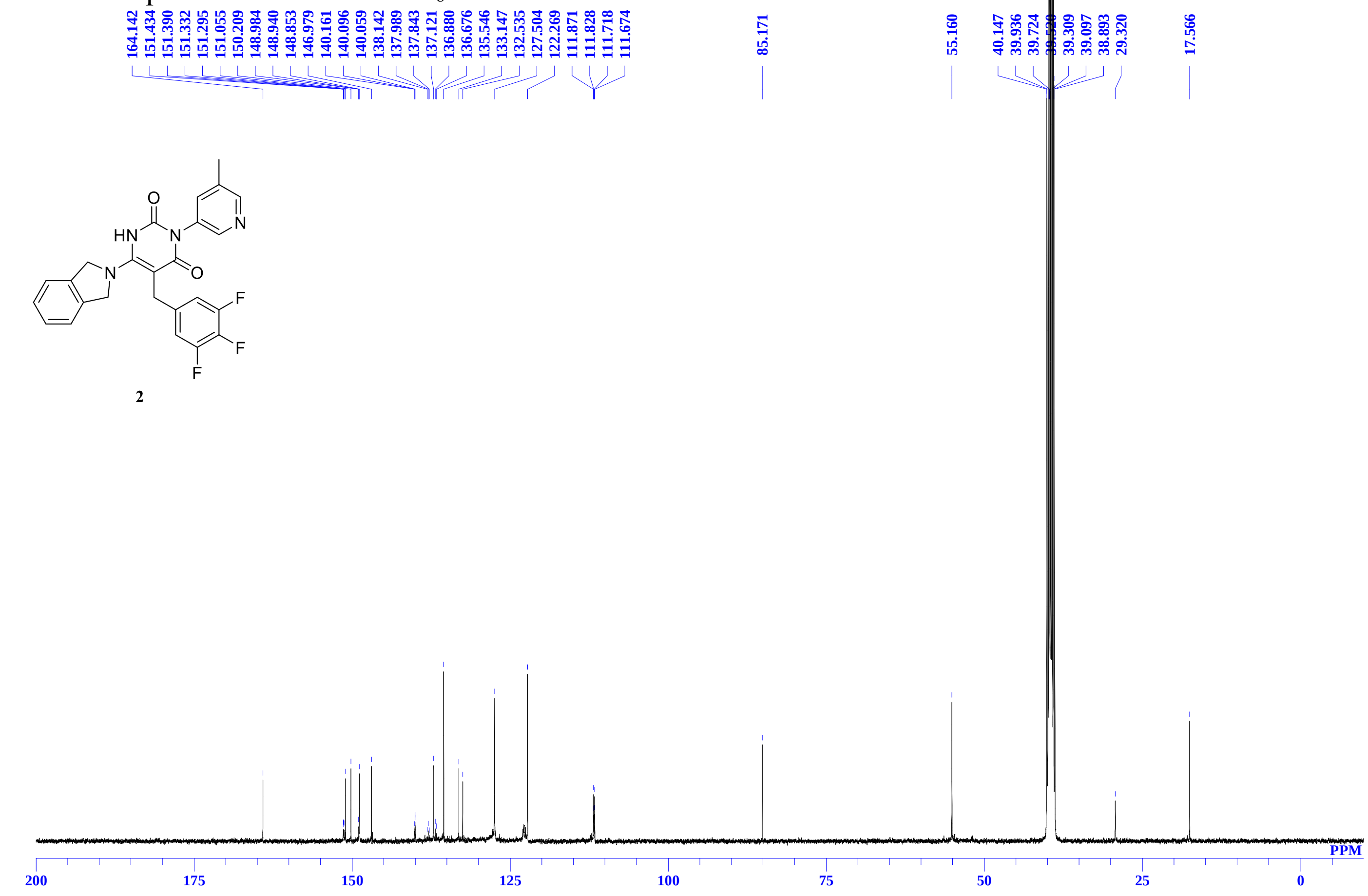

$^{1}$H NMR spectra of **3** in $CDCl_3$

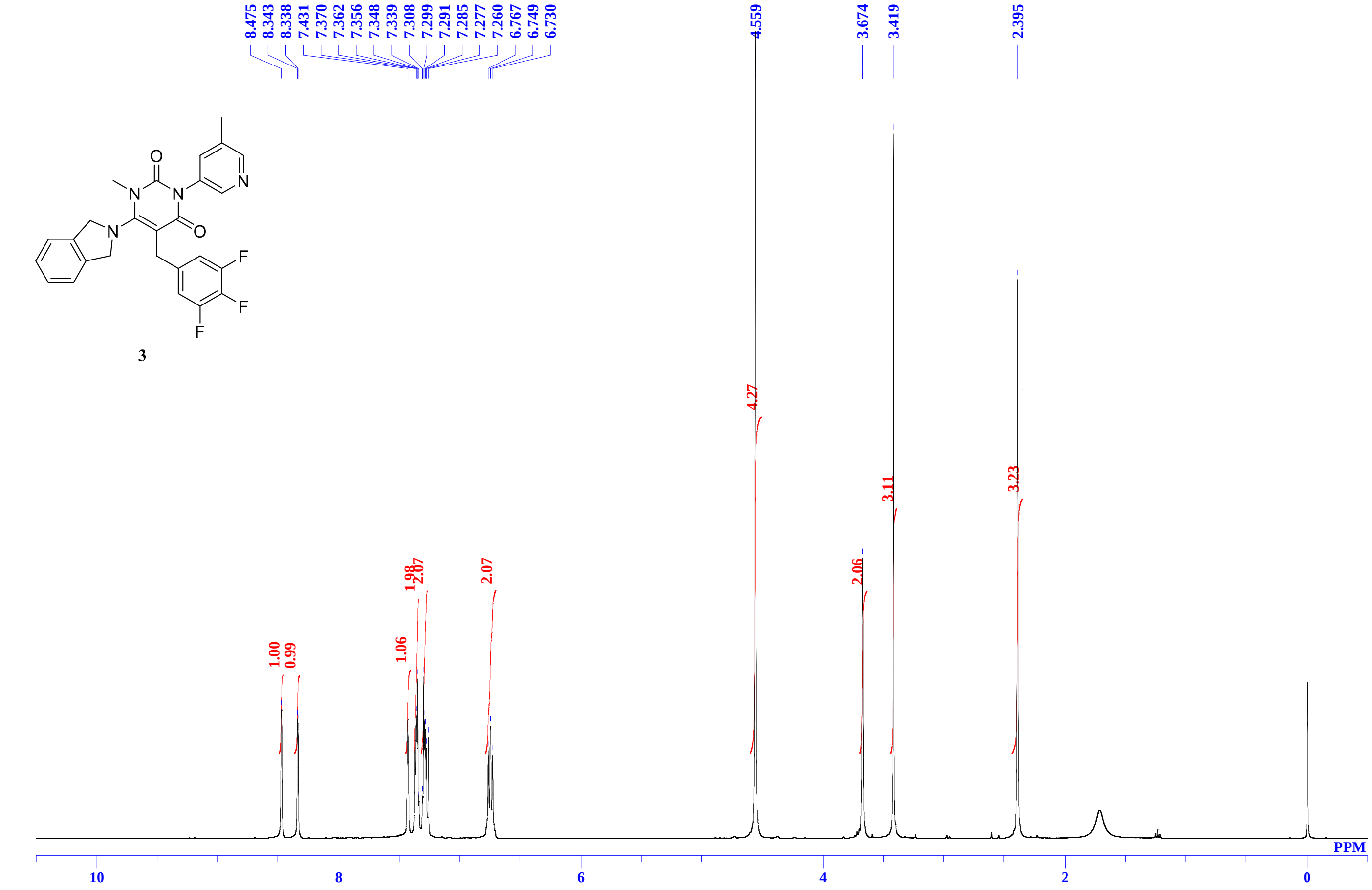


$^{13}$C NMR spectra of **3** in $CDCl_3$

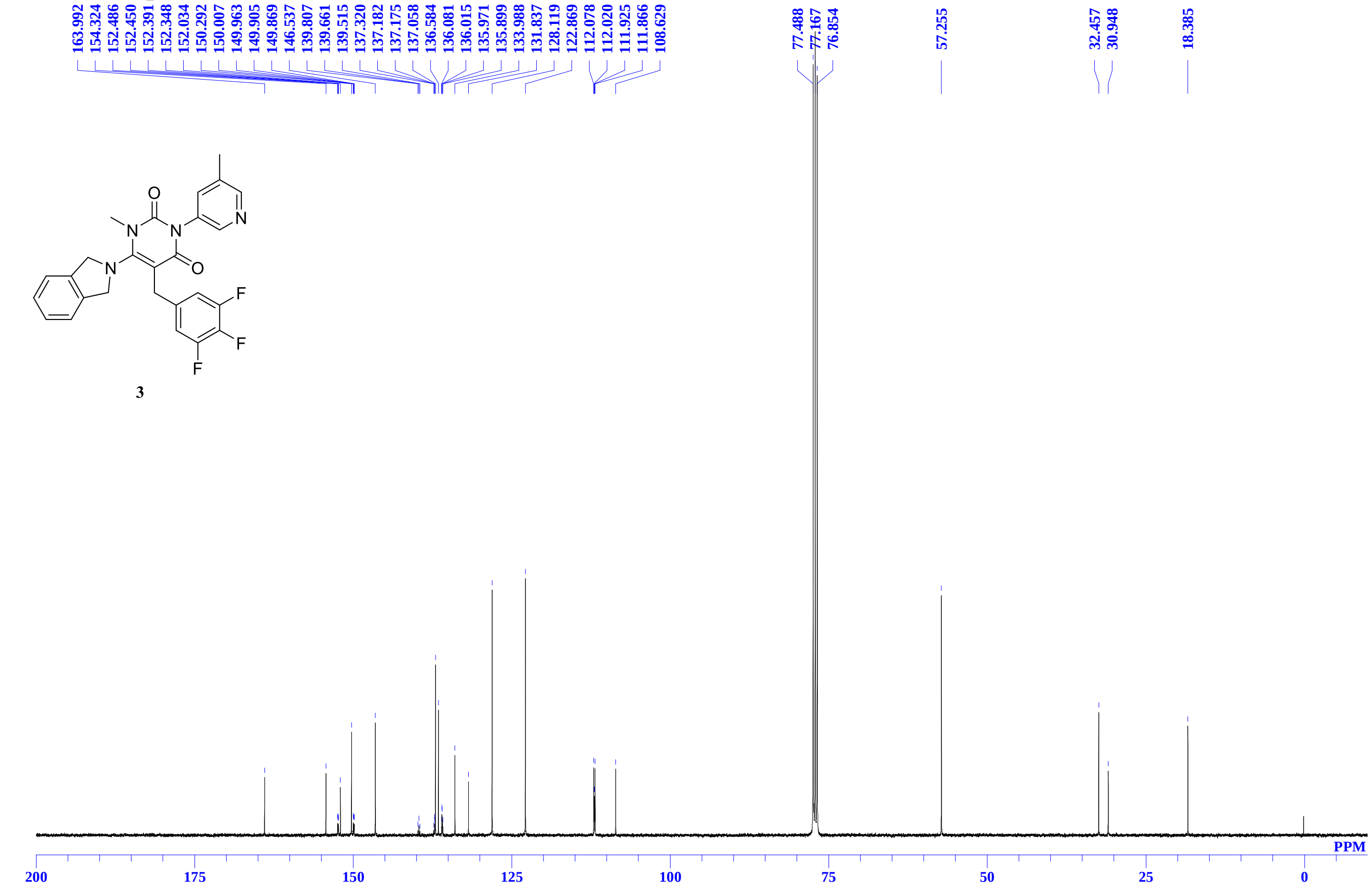

$^{1}$H NMR spectra of **4** in $CDCl_3$

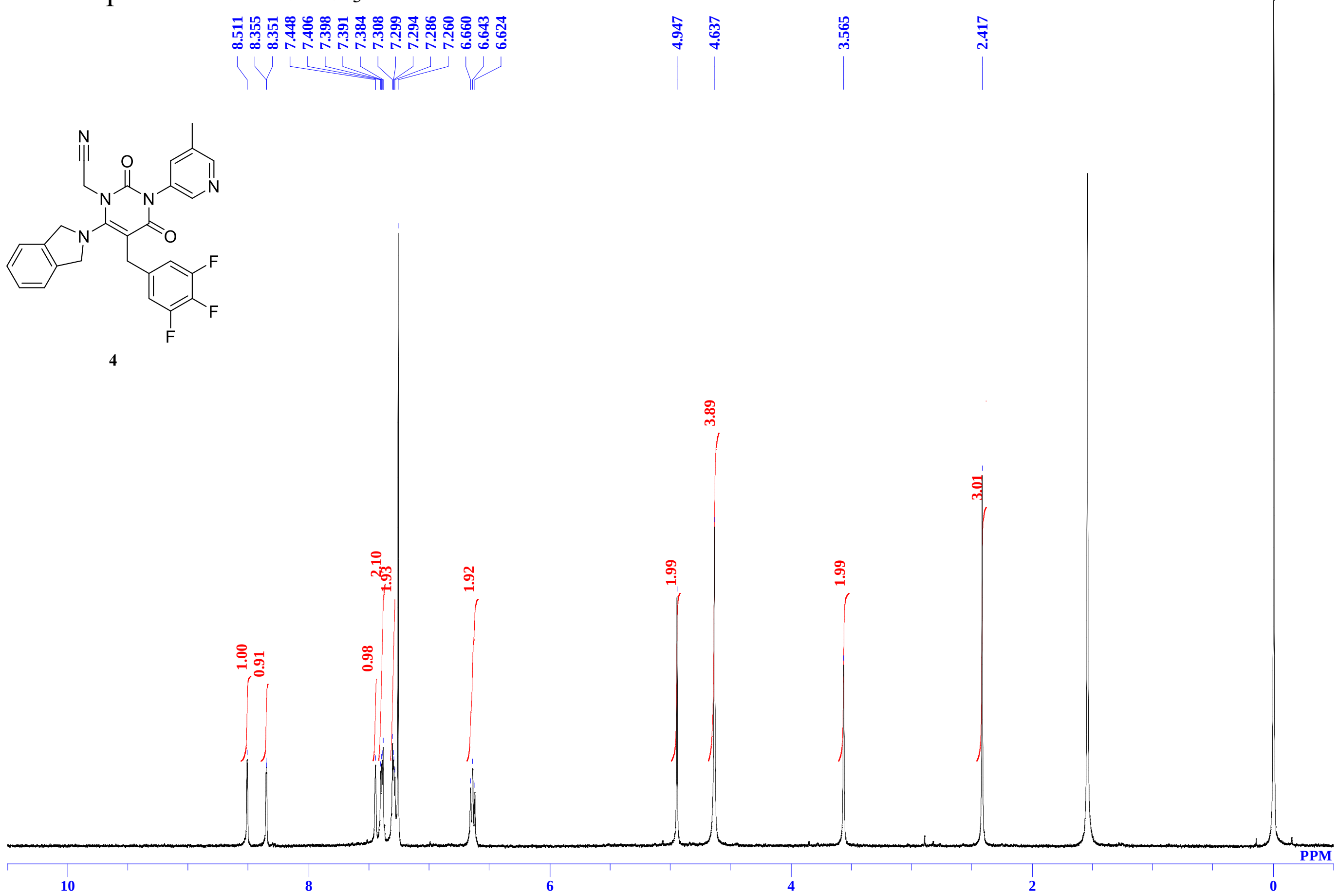


**$^{13}$C** NMR spectra of **4** in $CDCl_3$

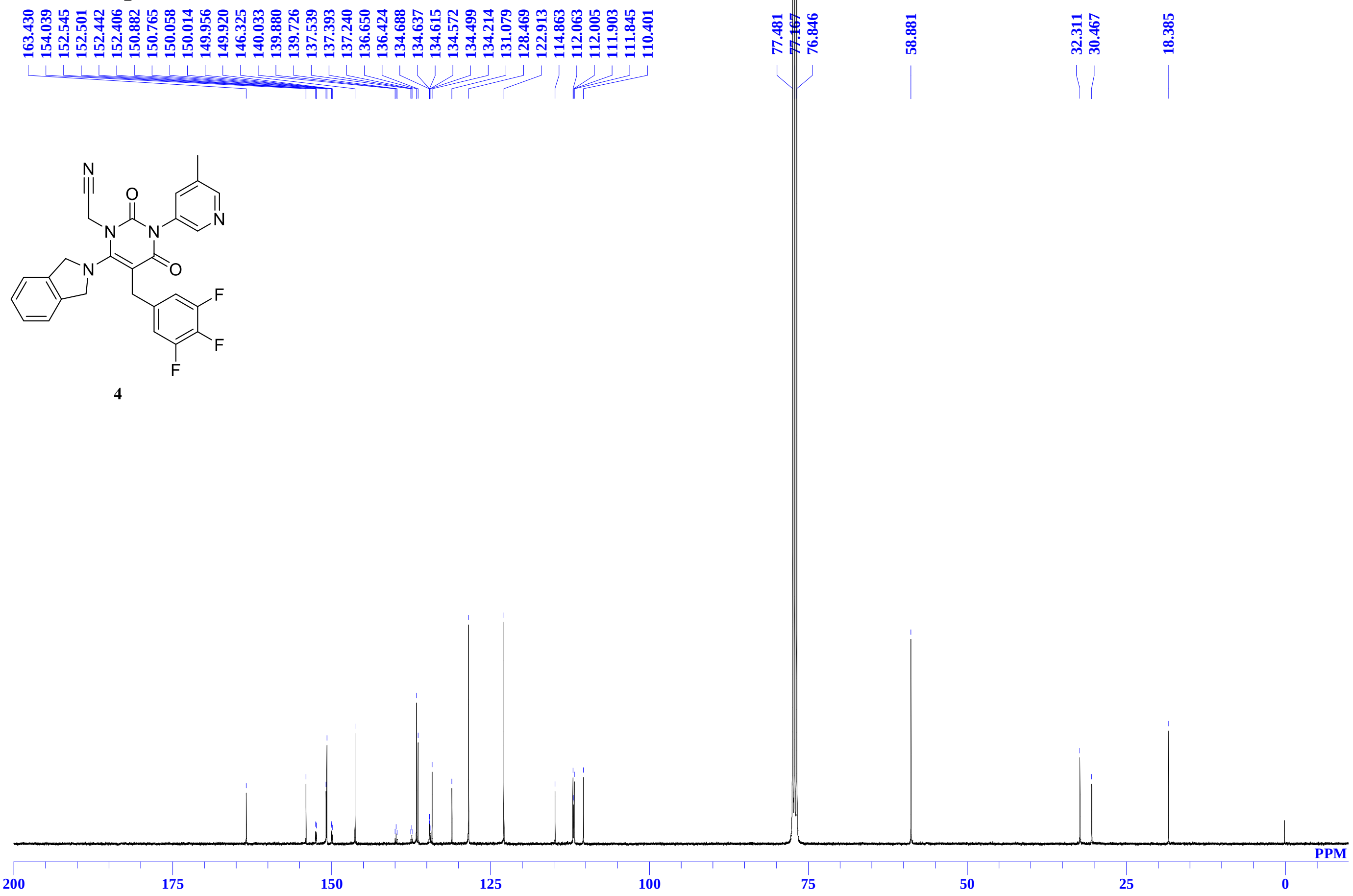

### $^1$H NMR spectra of **5** in $CDCl_3$

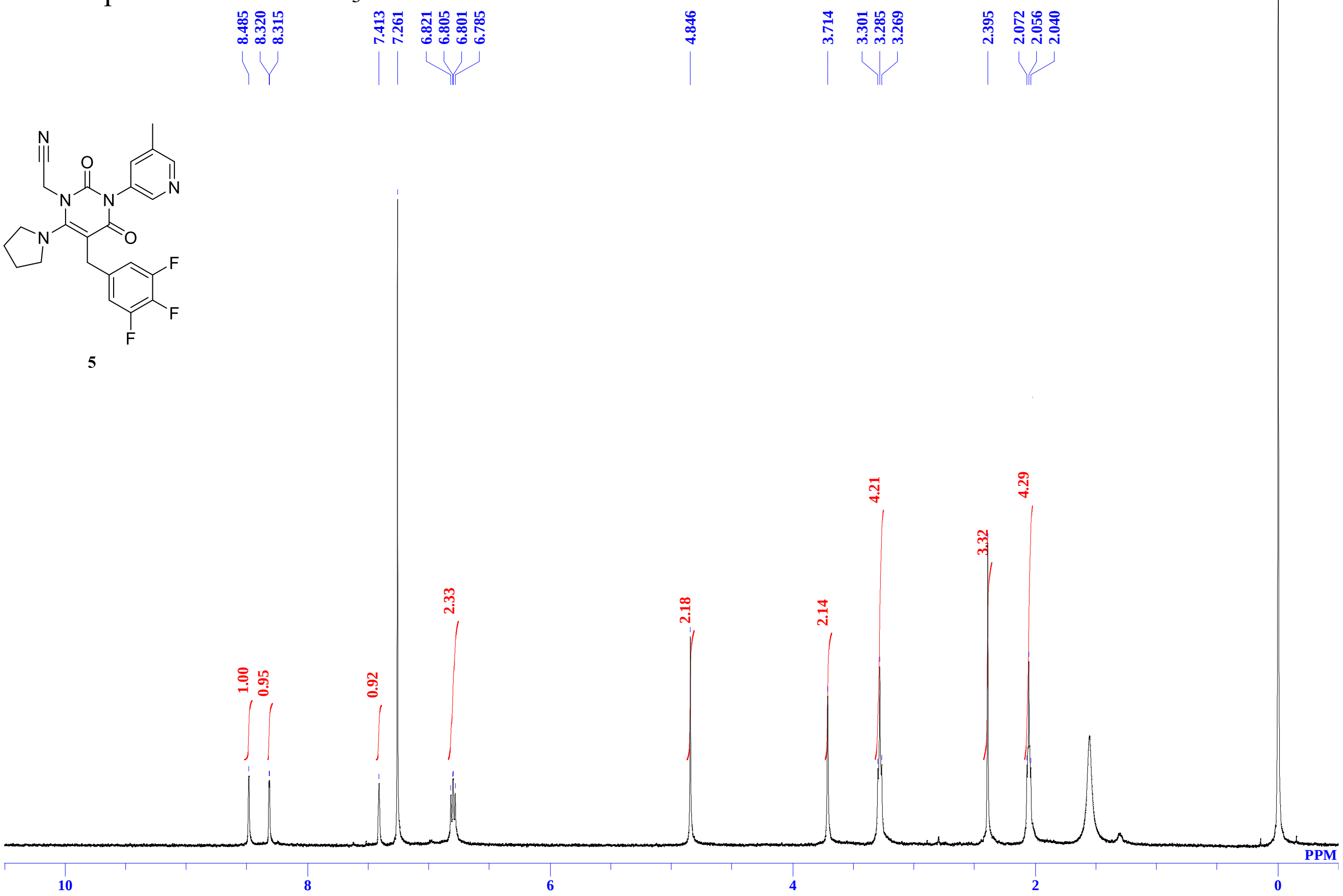


**5**

### $^{13}$C NMR spectra of **5** in DMSO-$d_6$

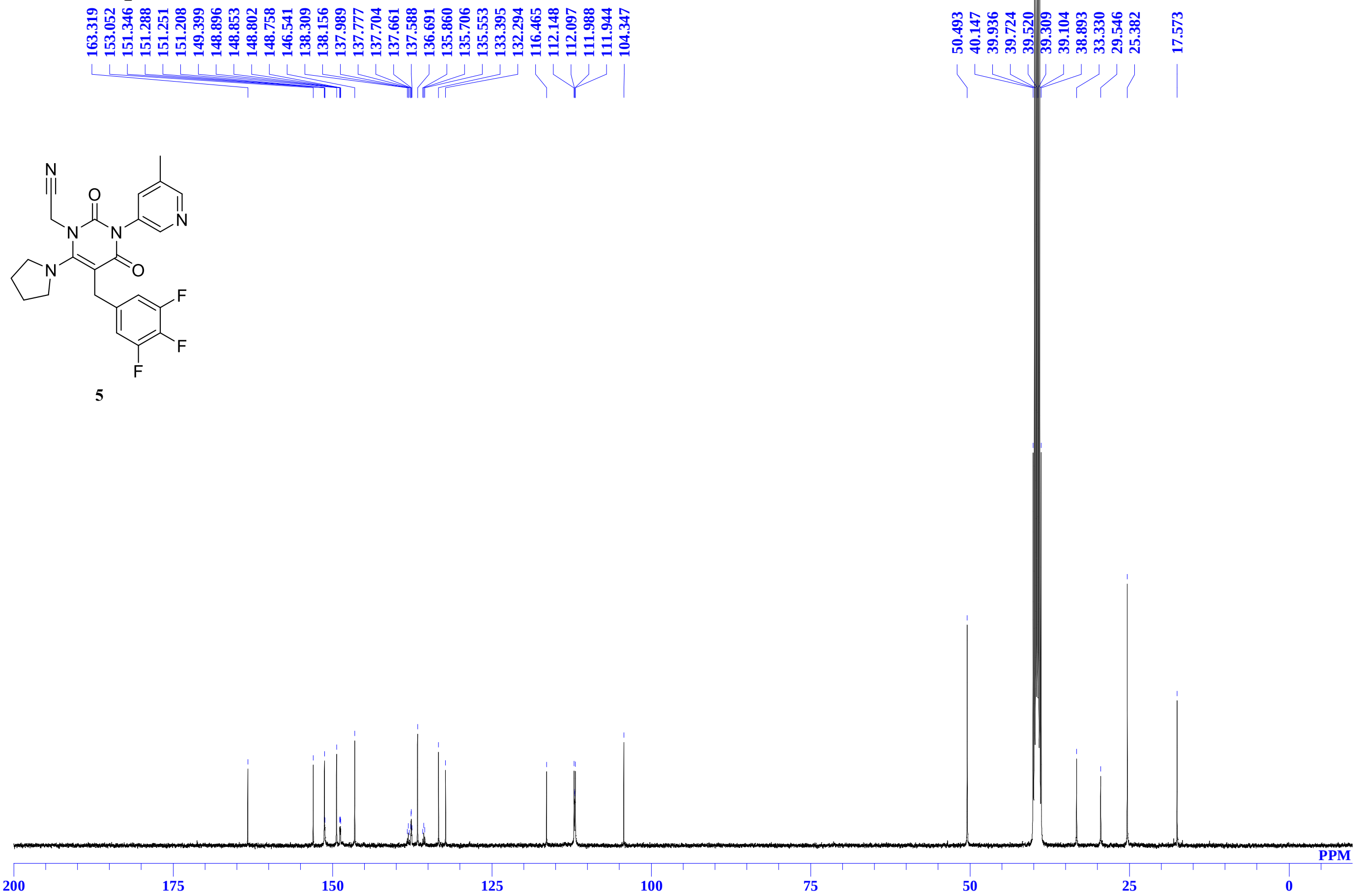


**5**

### $^{1}$H NMR spectra of **6** in DMSO-$d_6$

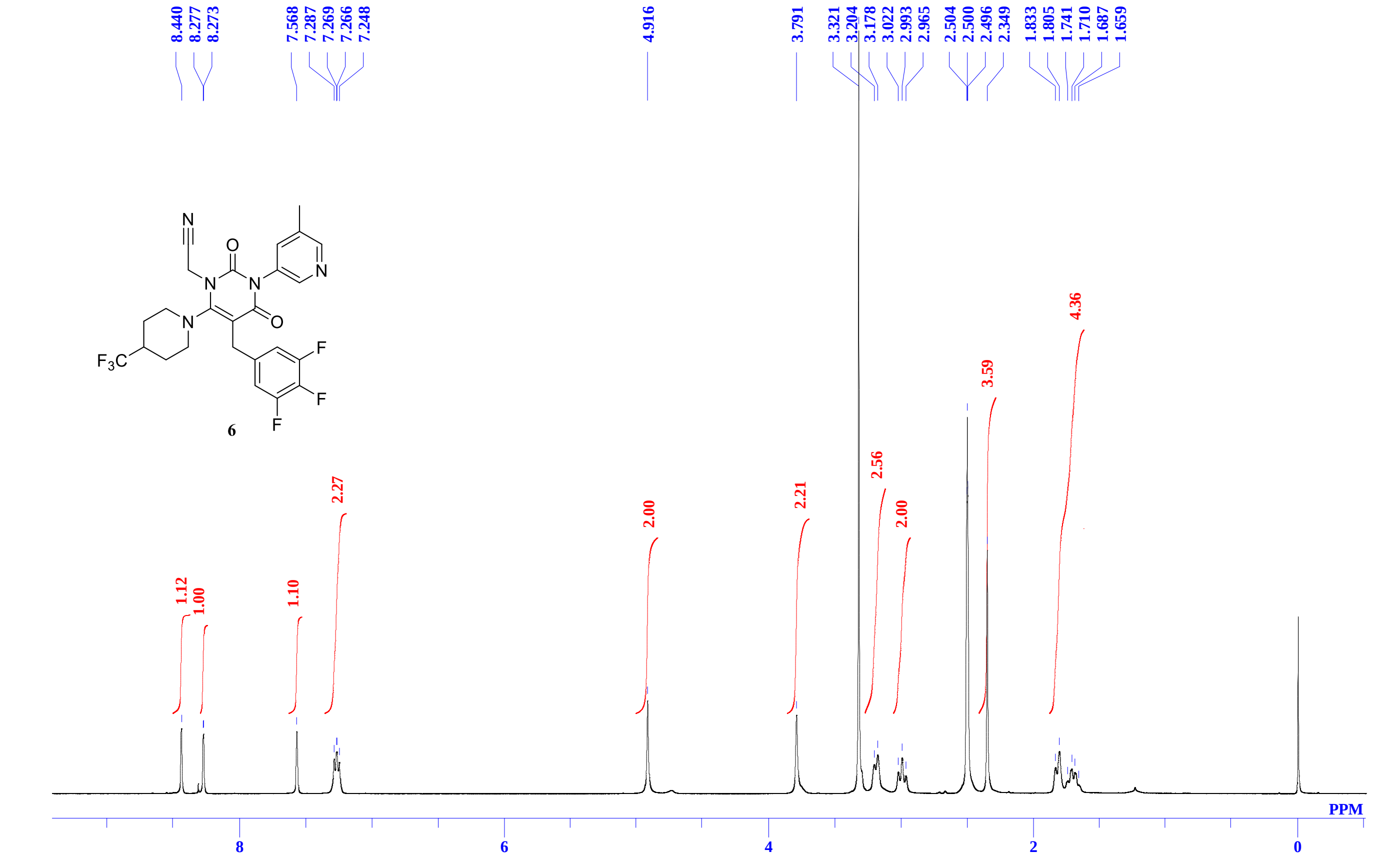


### $^{13}$C NMR spectra of **6** in DMSO-$d_6$

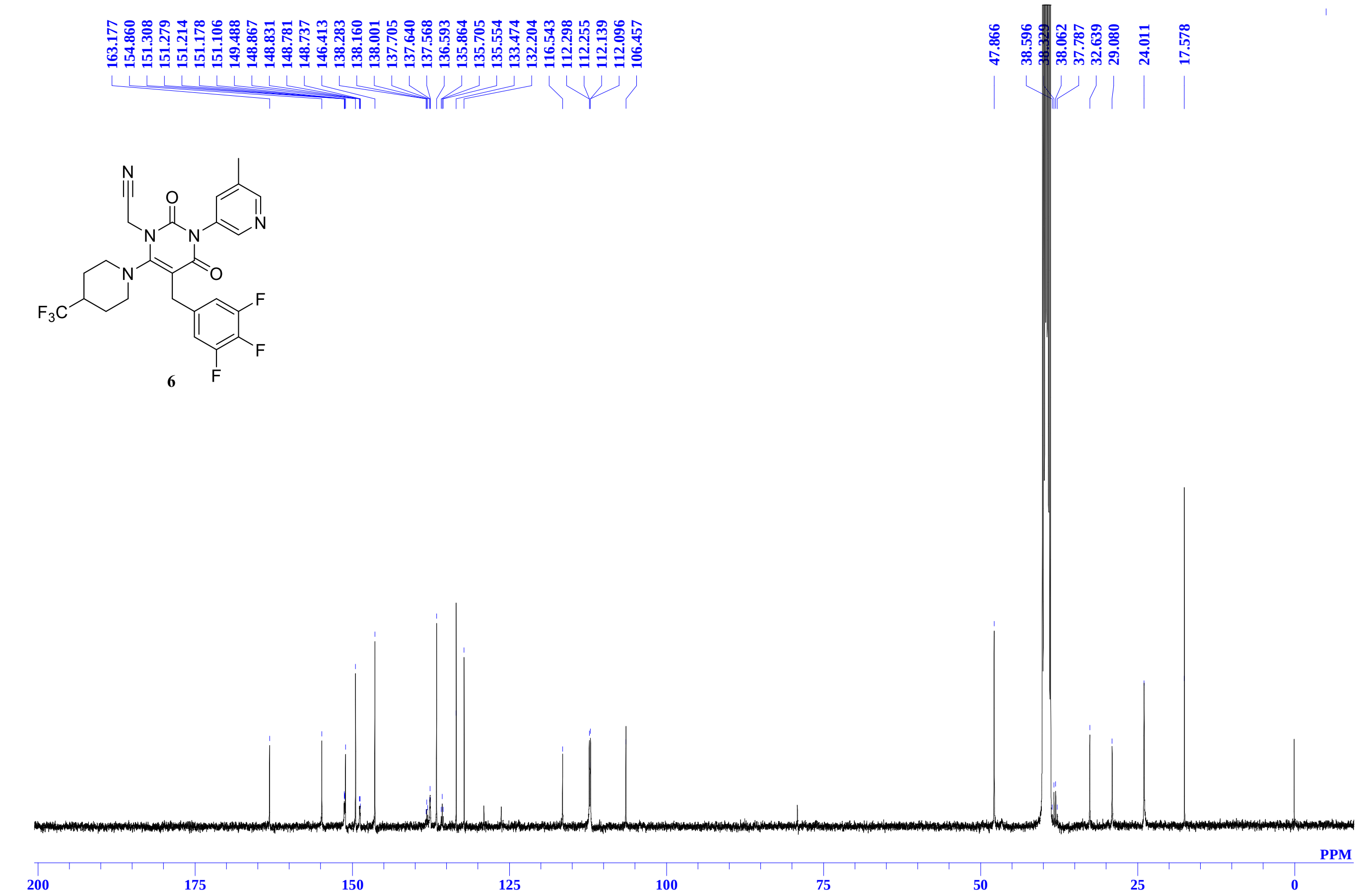

$^{1}H$ NMR spectra of **7** in DMSO-$d_6$

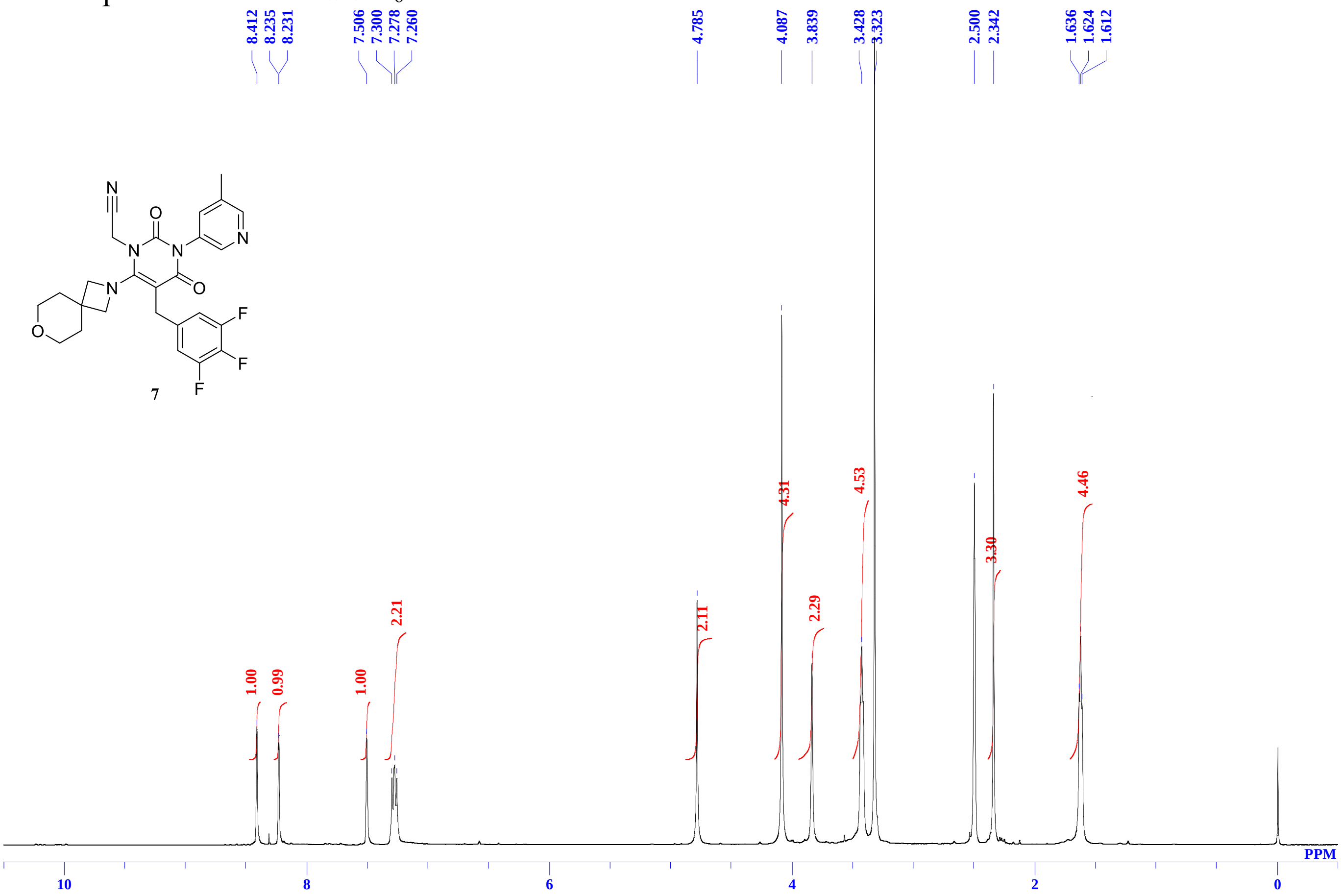


$^{13}C$ NMR spectra of **7** in DMSO-$d_6$

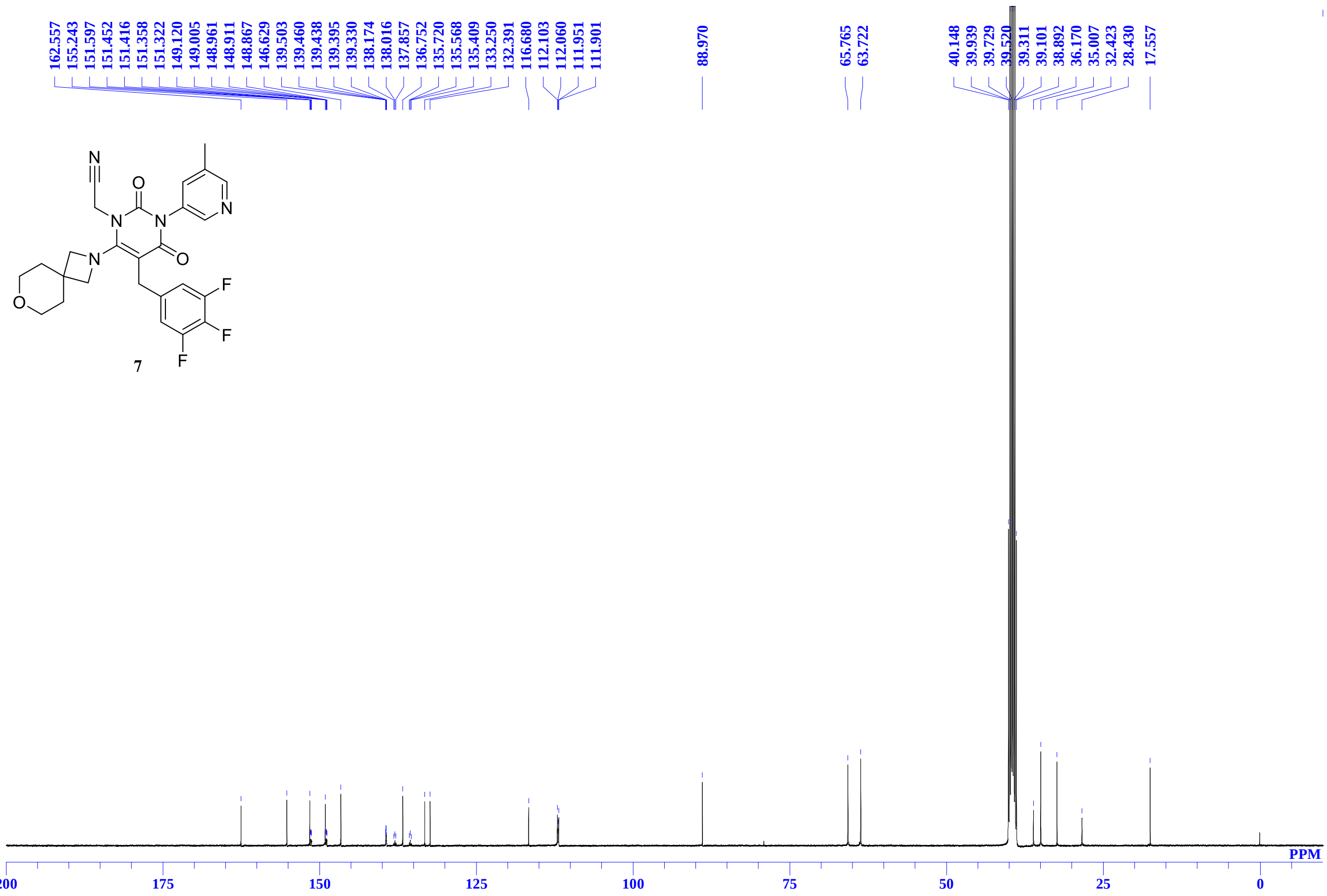

[1]H NMR spectra of **8** in DMSO-$d_6$

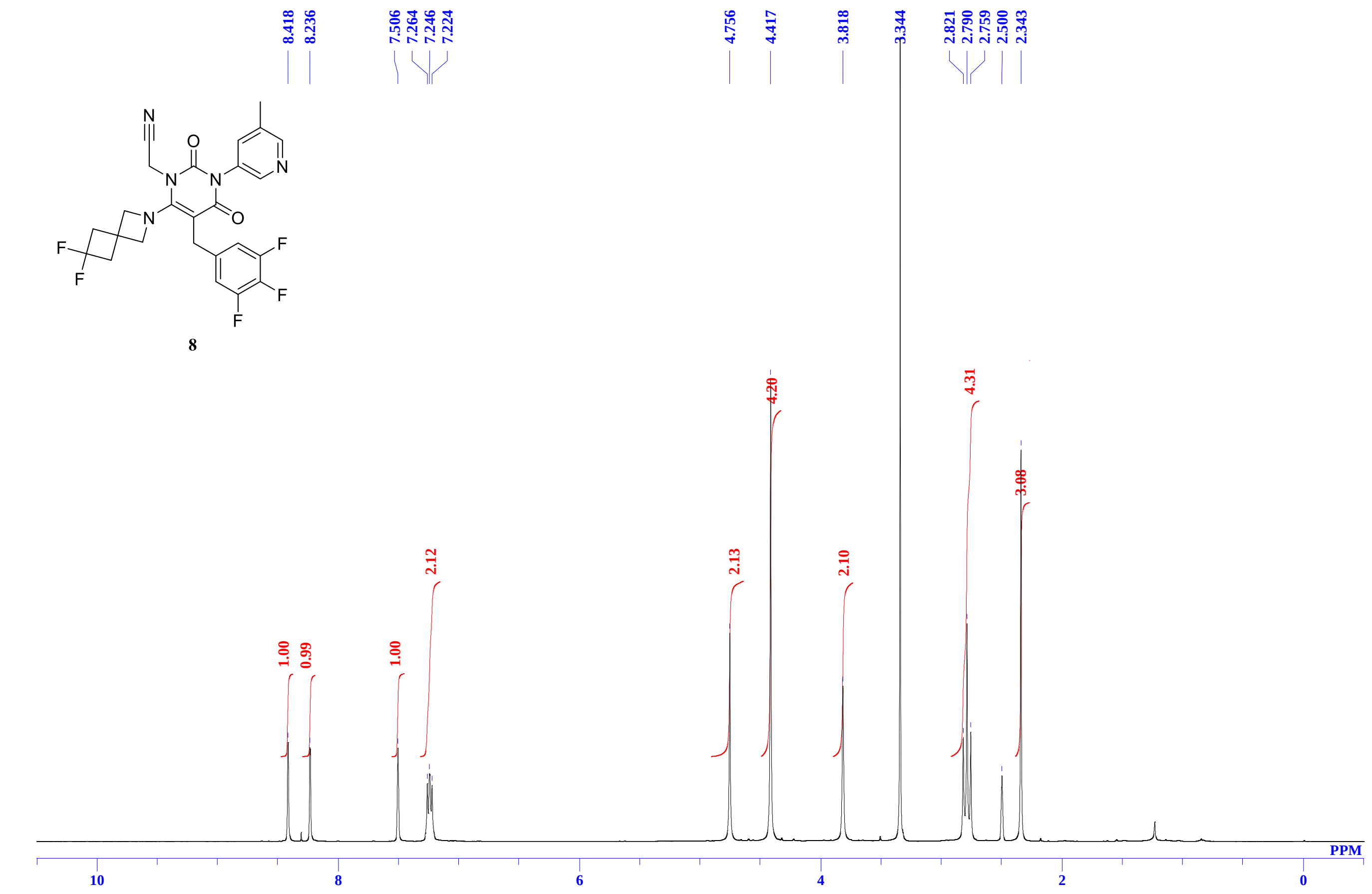


**13C** NMR spectra of **8** in DMSO-$d_6$

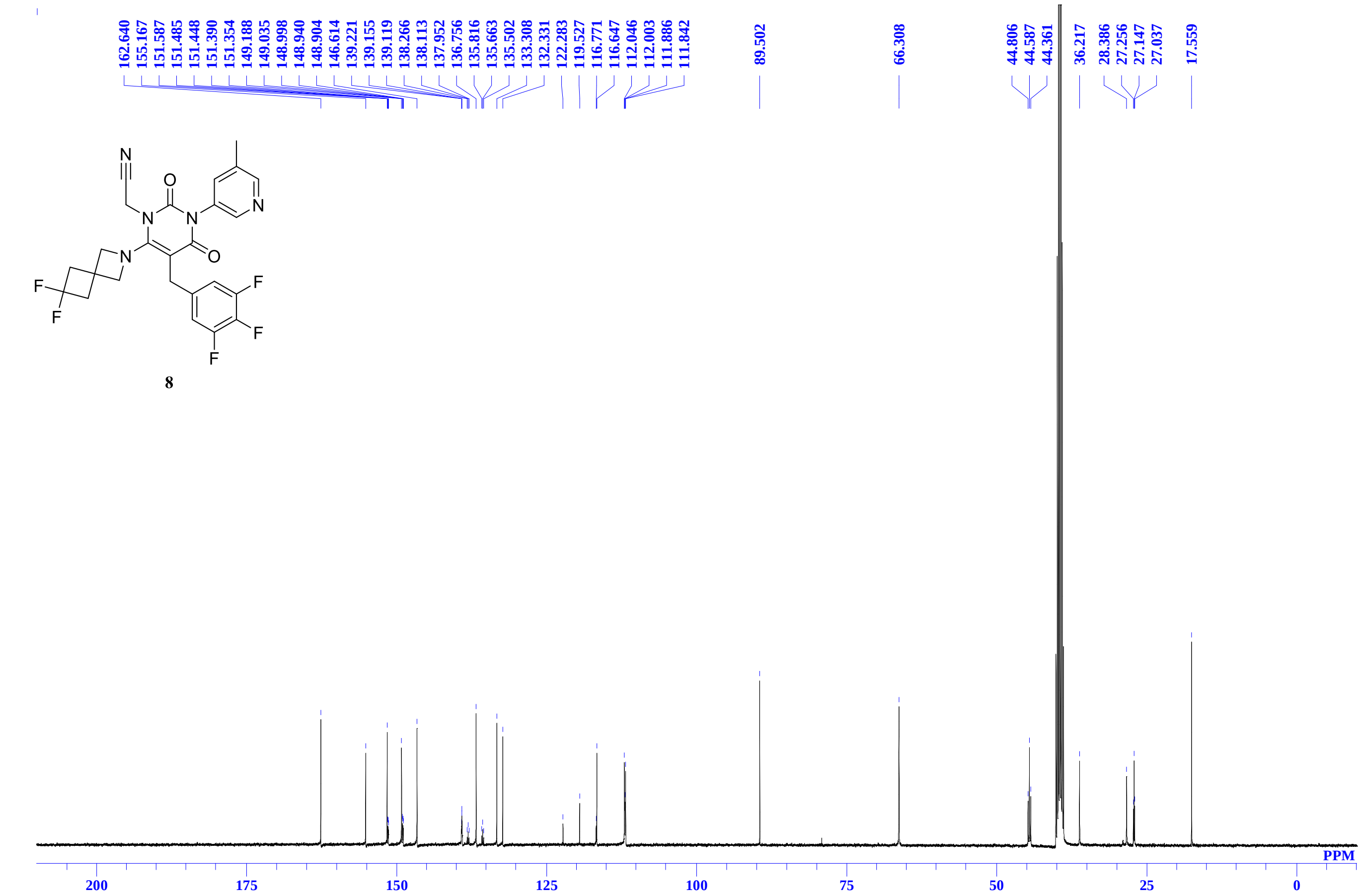

[1]H NMR spectra of **9** in DMSO-$d_6$

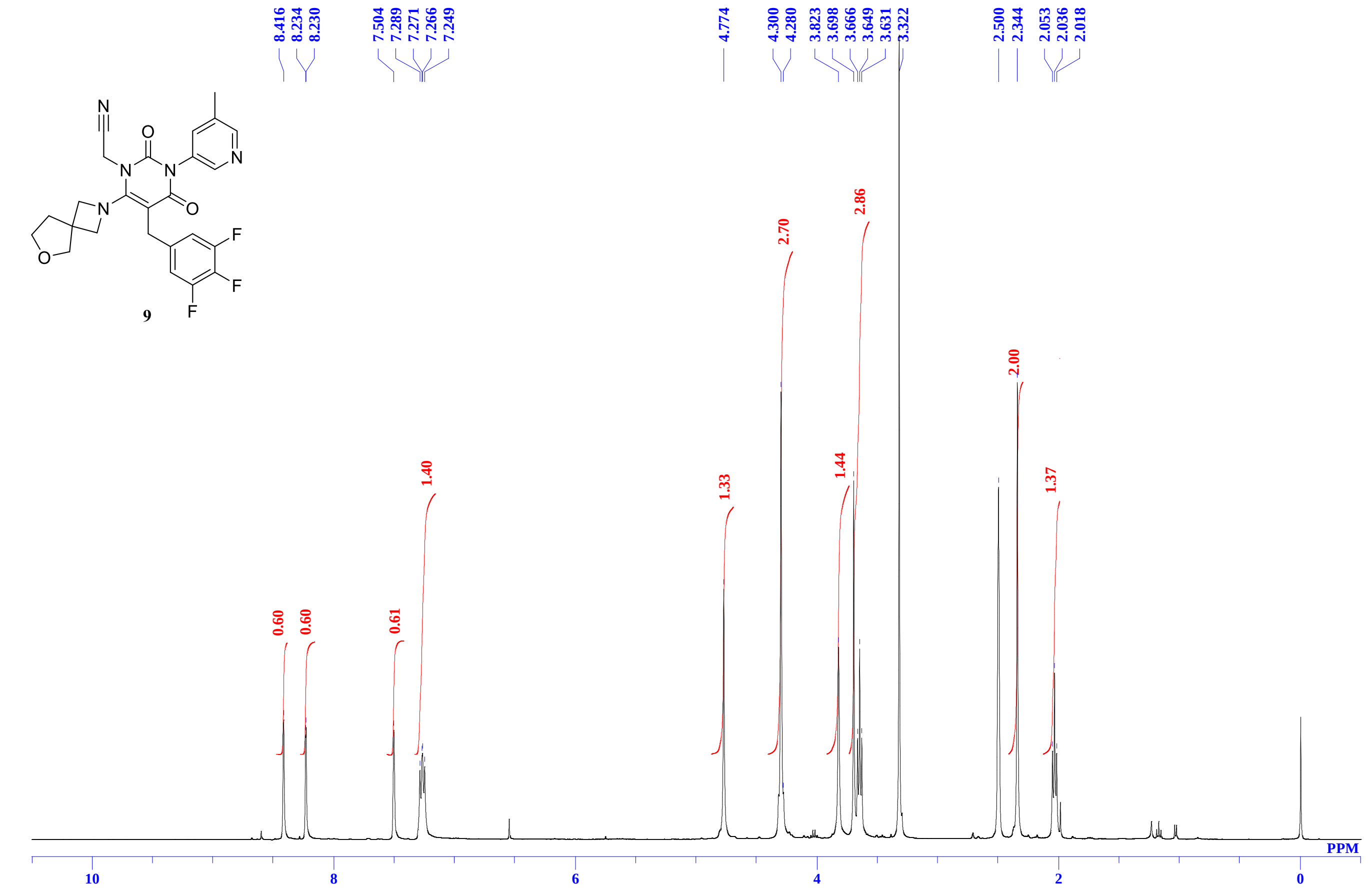


**[13]C** NMR spectra of **9** in DMSO-$d_6$

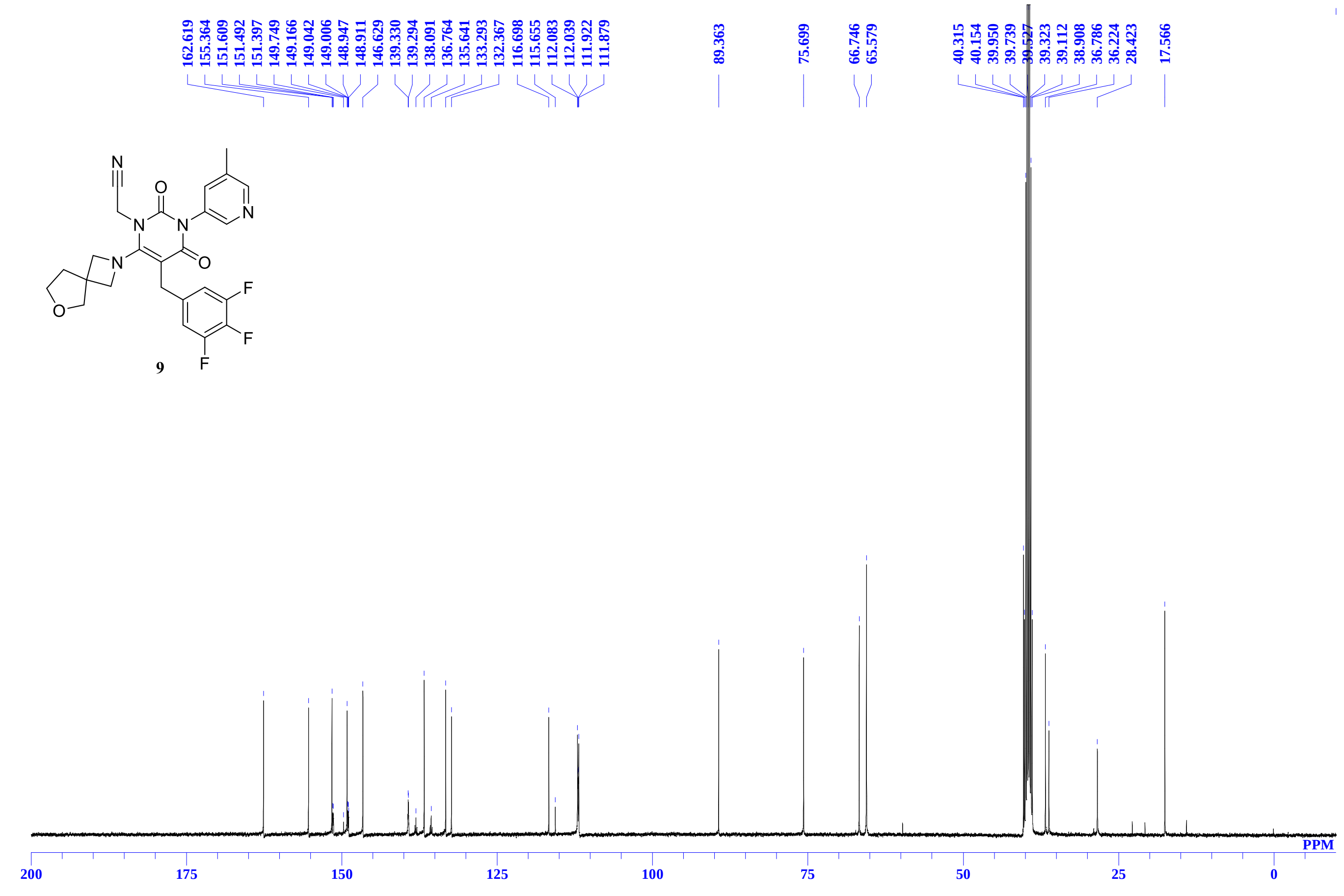

$^1$H NMR spectra of **10** in DMSO-$d_6$

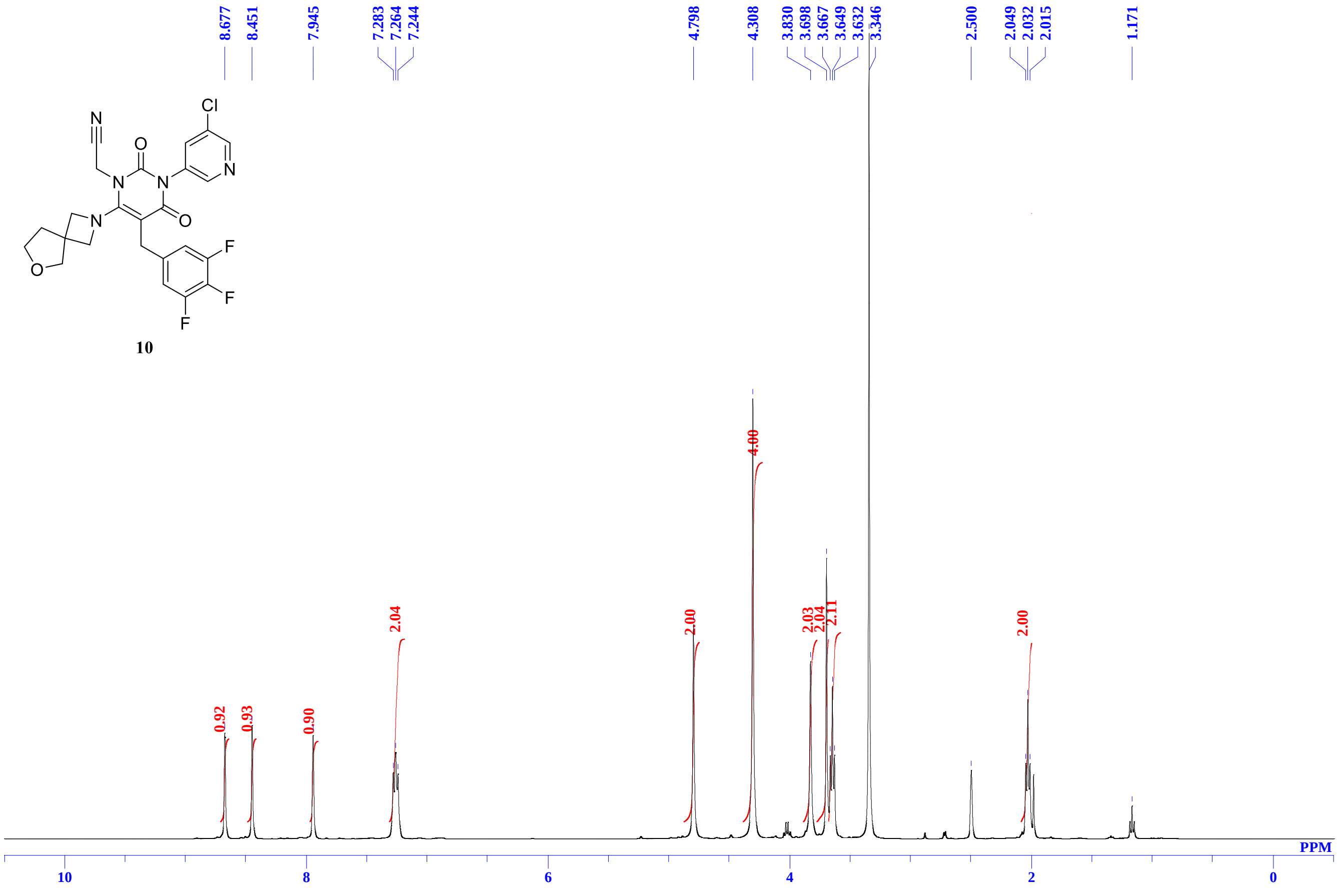


$^{13}$C NMR spectra of **10** in DMSO-$d_6$

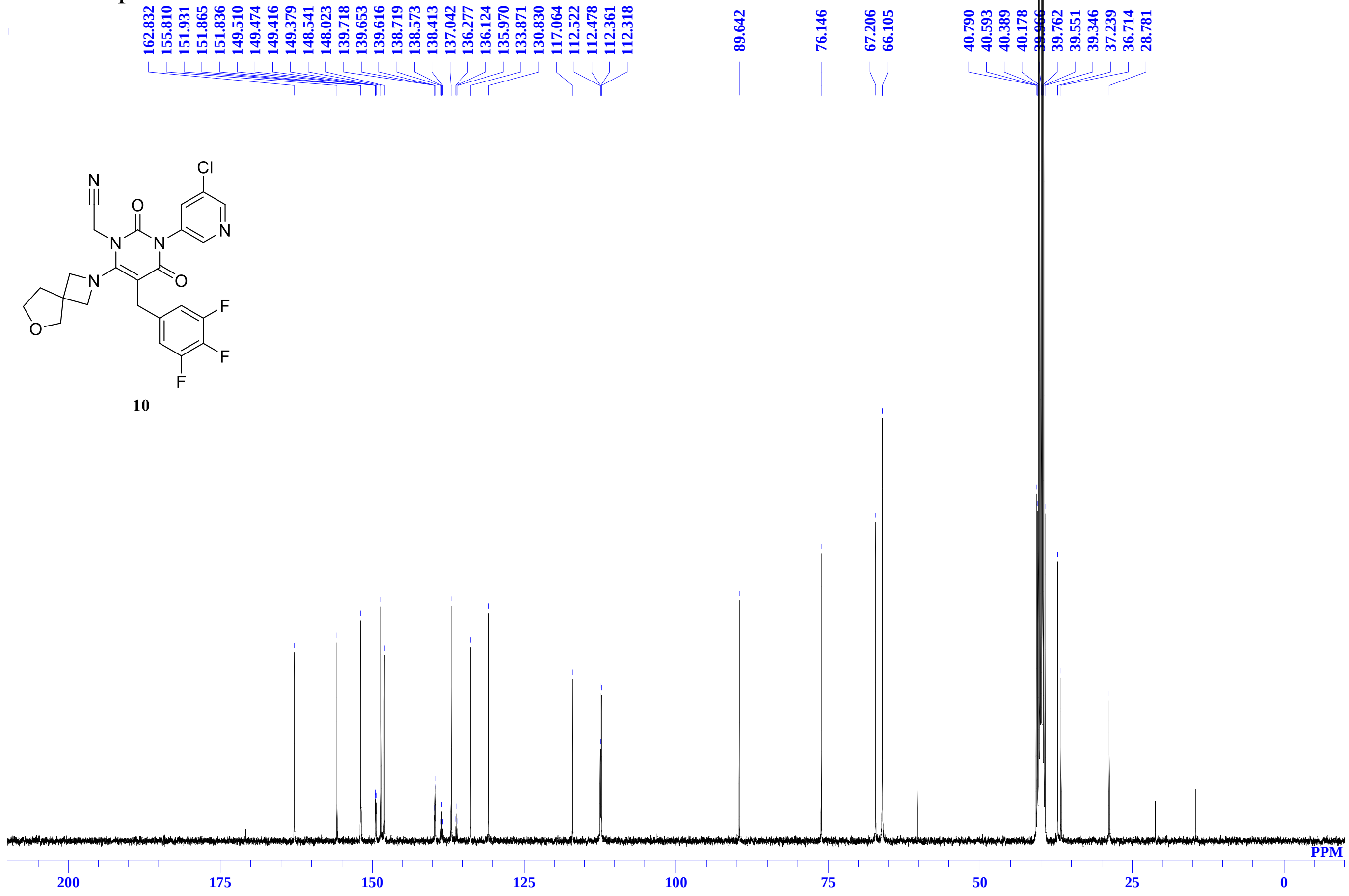

[1]H NMR spectra of **11** in $CDCl_3$

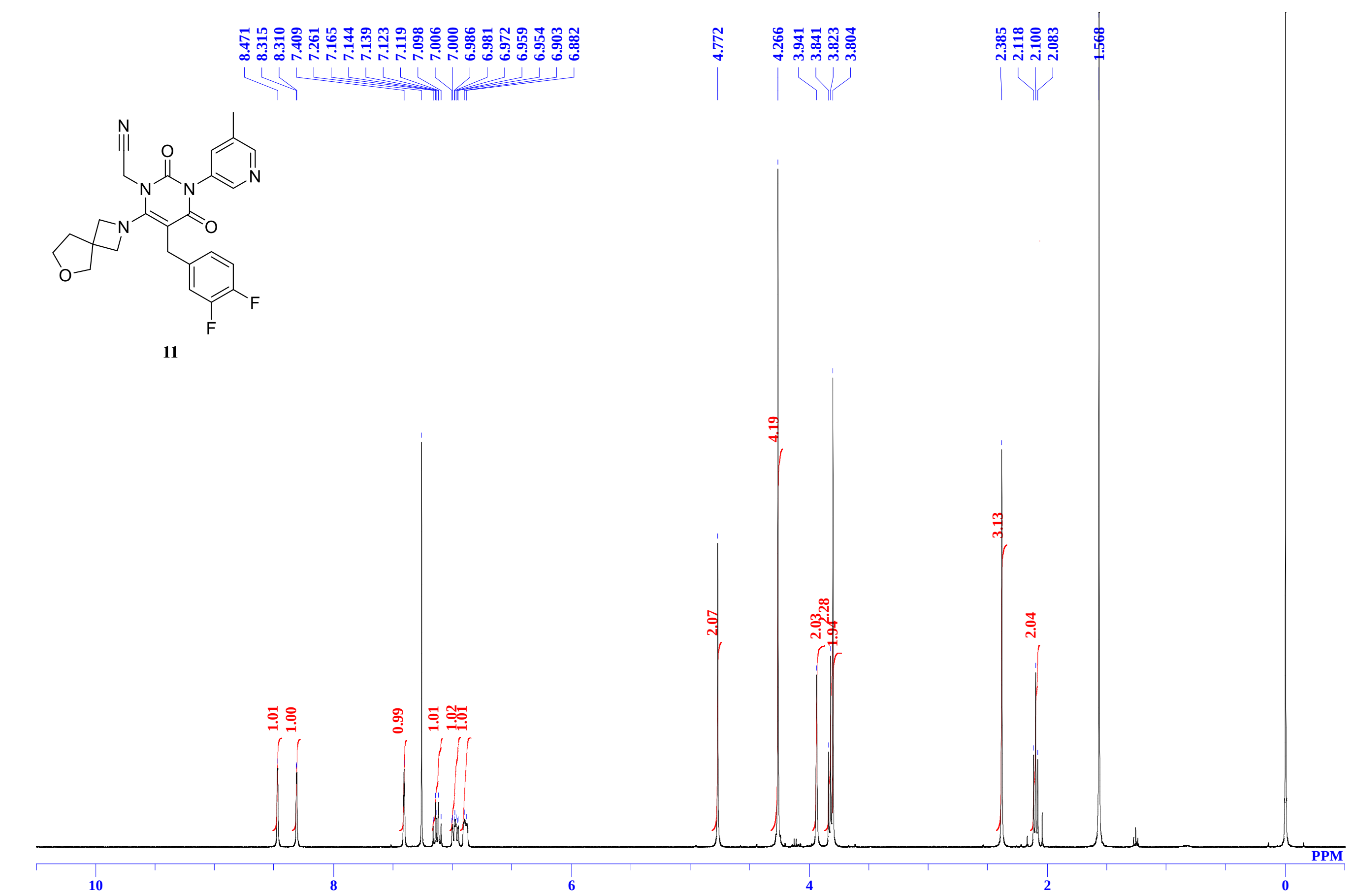


[13]C NMR spectra of **11** in DMSO-$d_6$

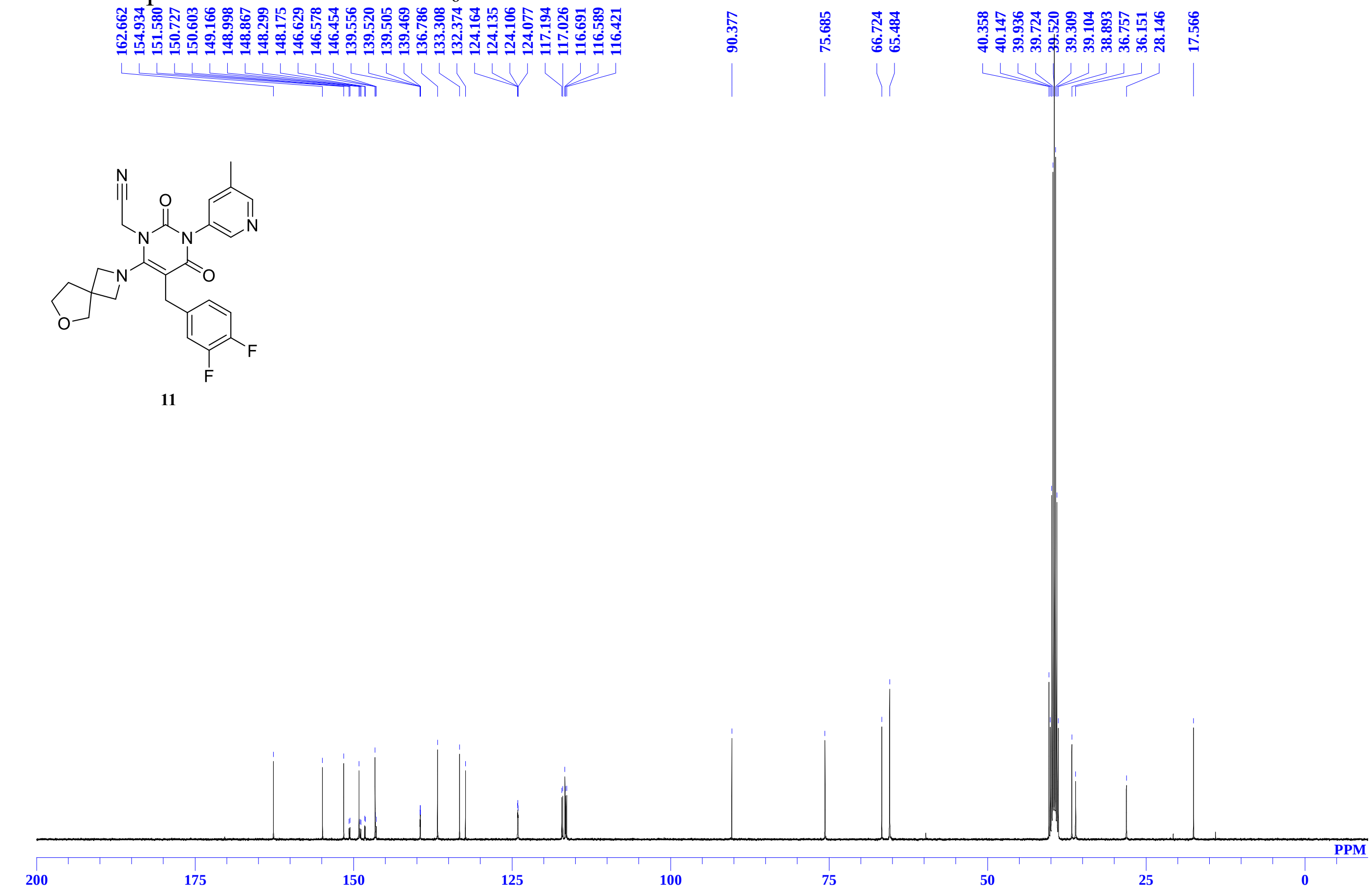

$^{1}H$ NMR spectra of **12** in $CDCl_3$

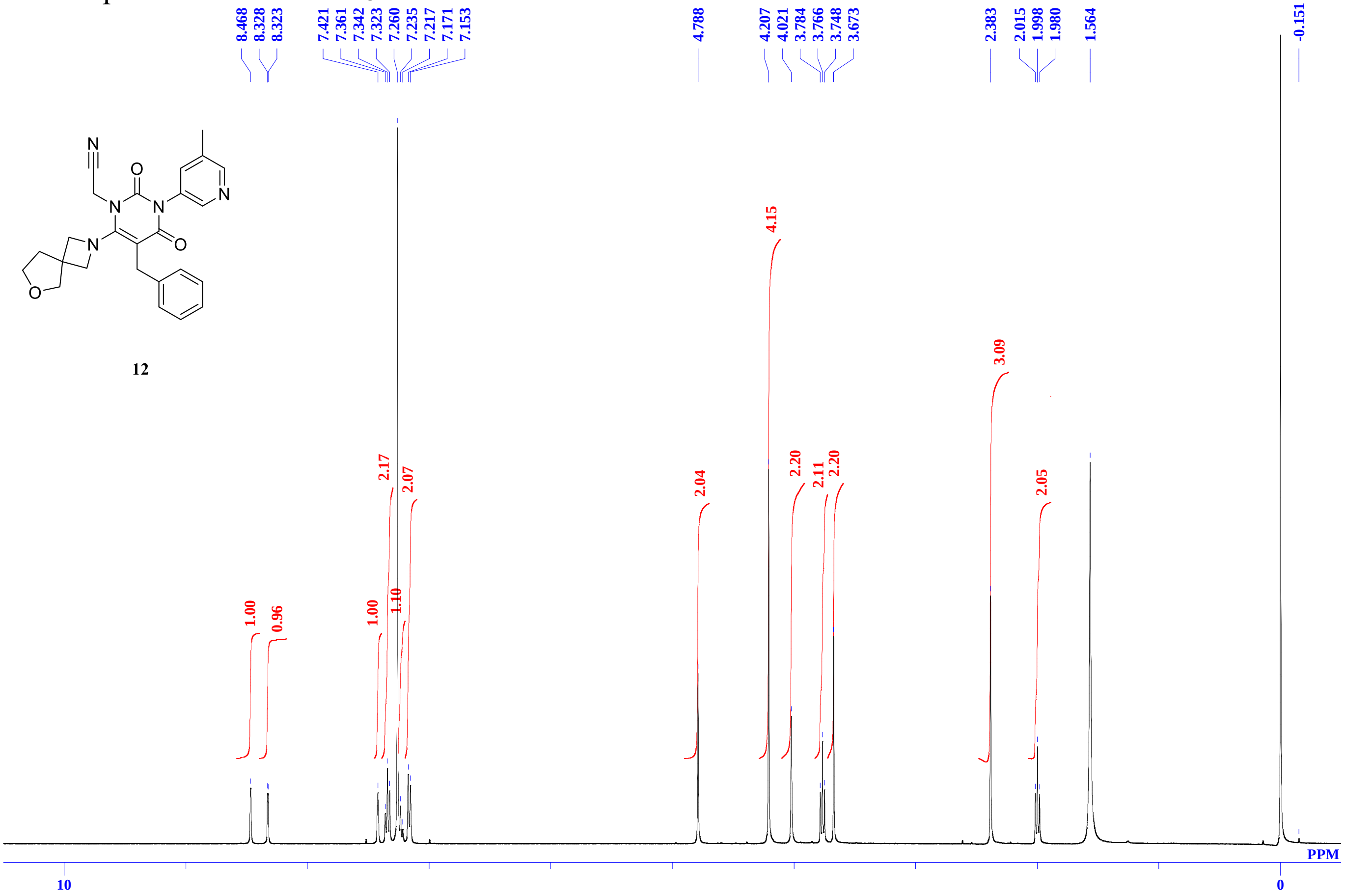


**$^{13}C$** NMR spectra of **12** in DMSO-$d_6$

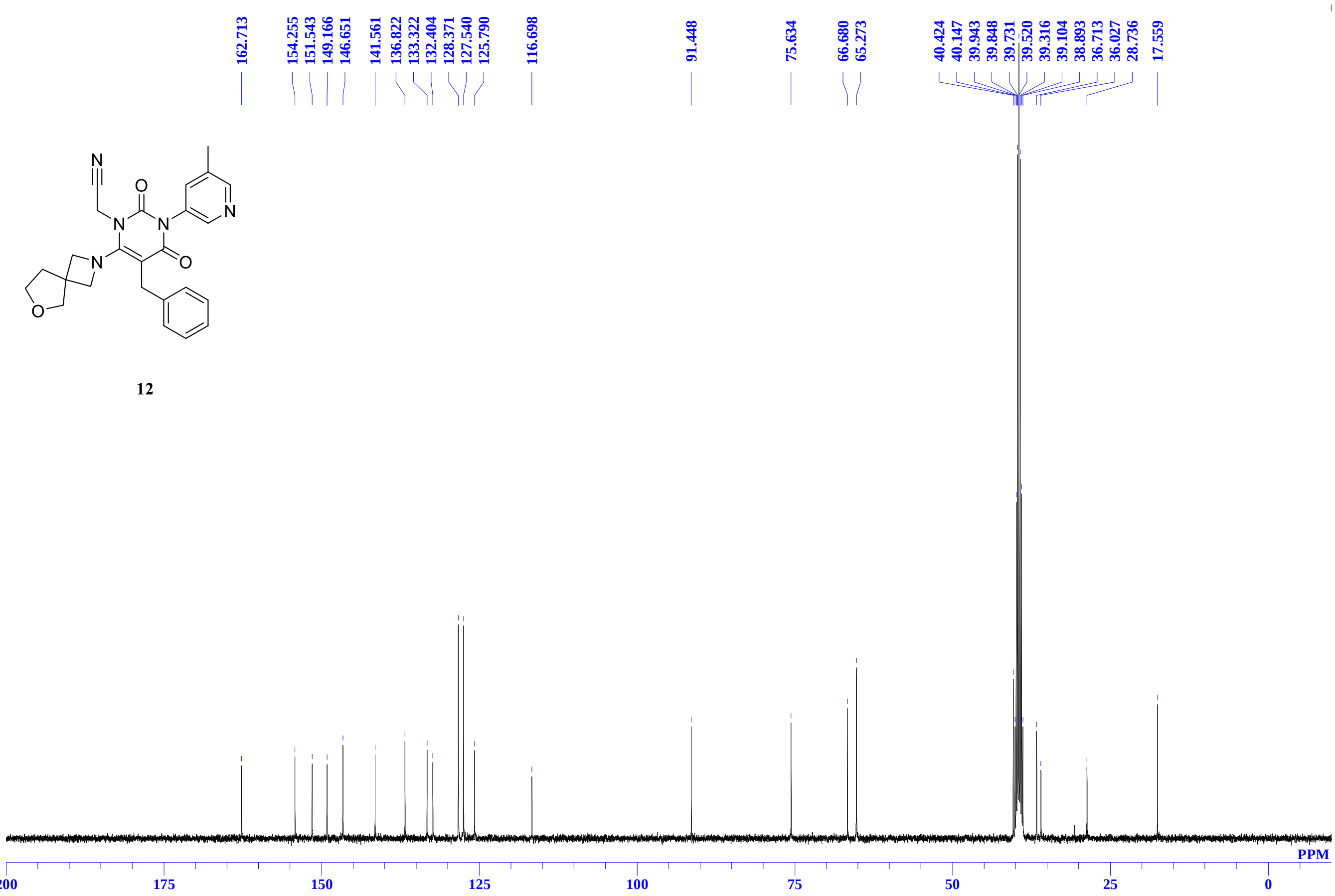

$^{1}H$ NMR spectra of **13** in $CDCl_3$

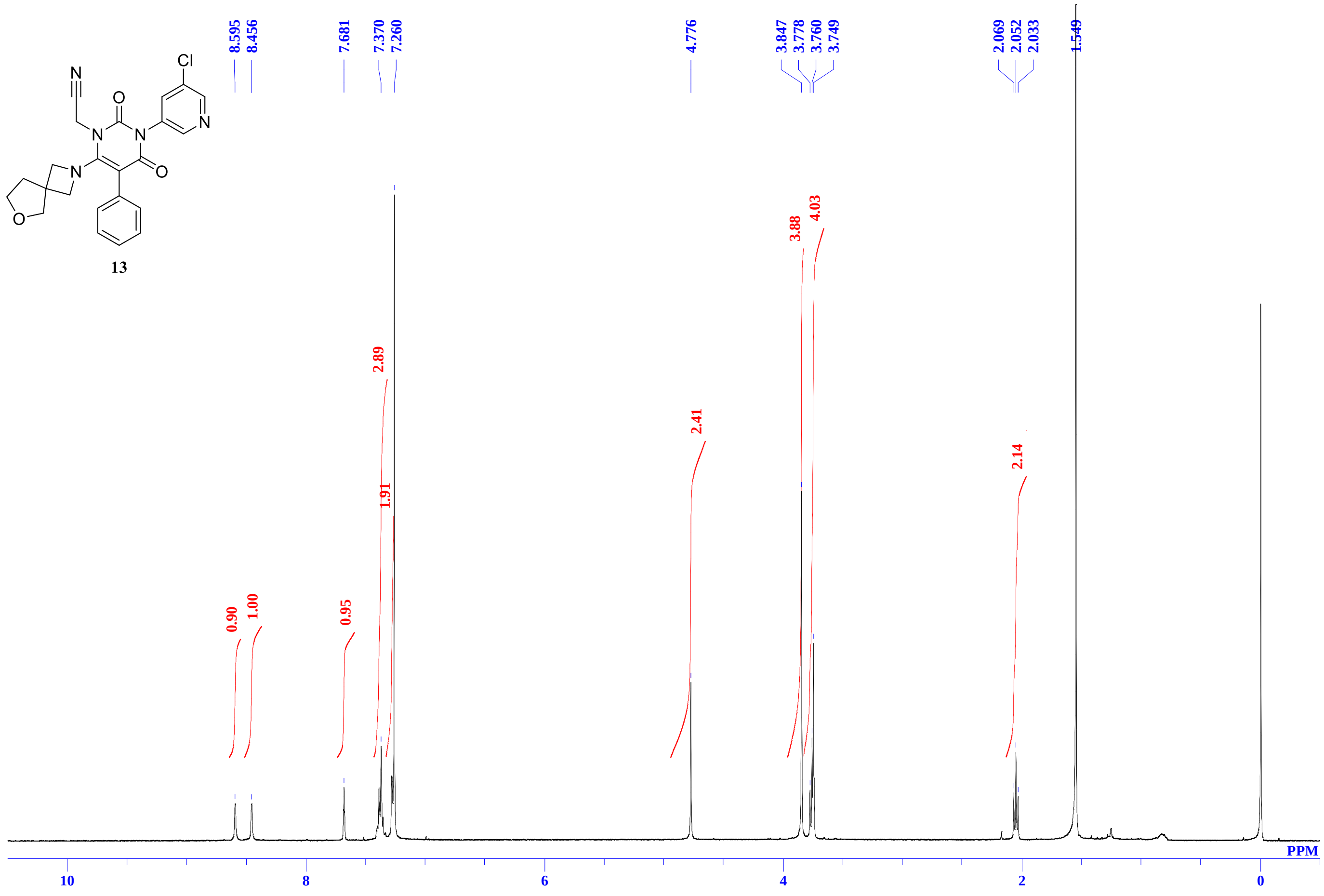


$^{13}C$ NMR spectra of **13** in $CDCl_3$

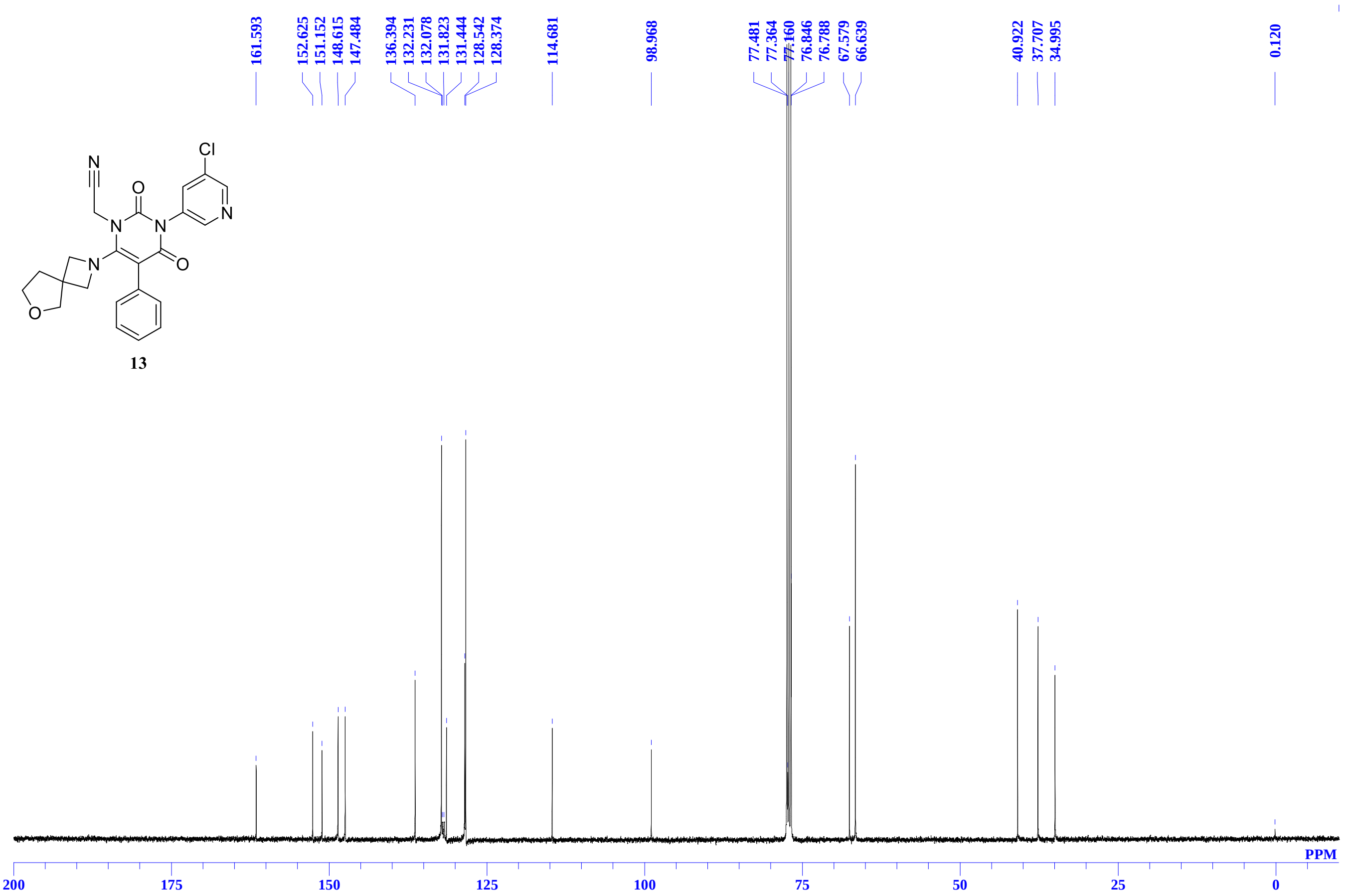

$^{1}H$ NMR spectra of **14** in $CDCl_3$

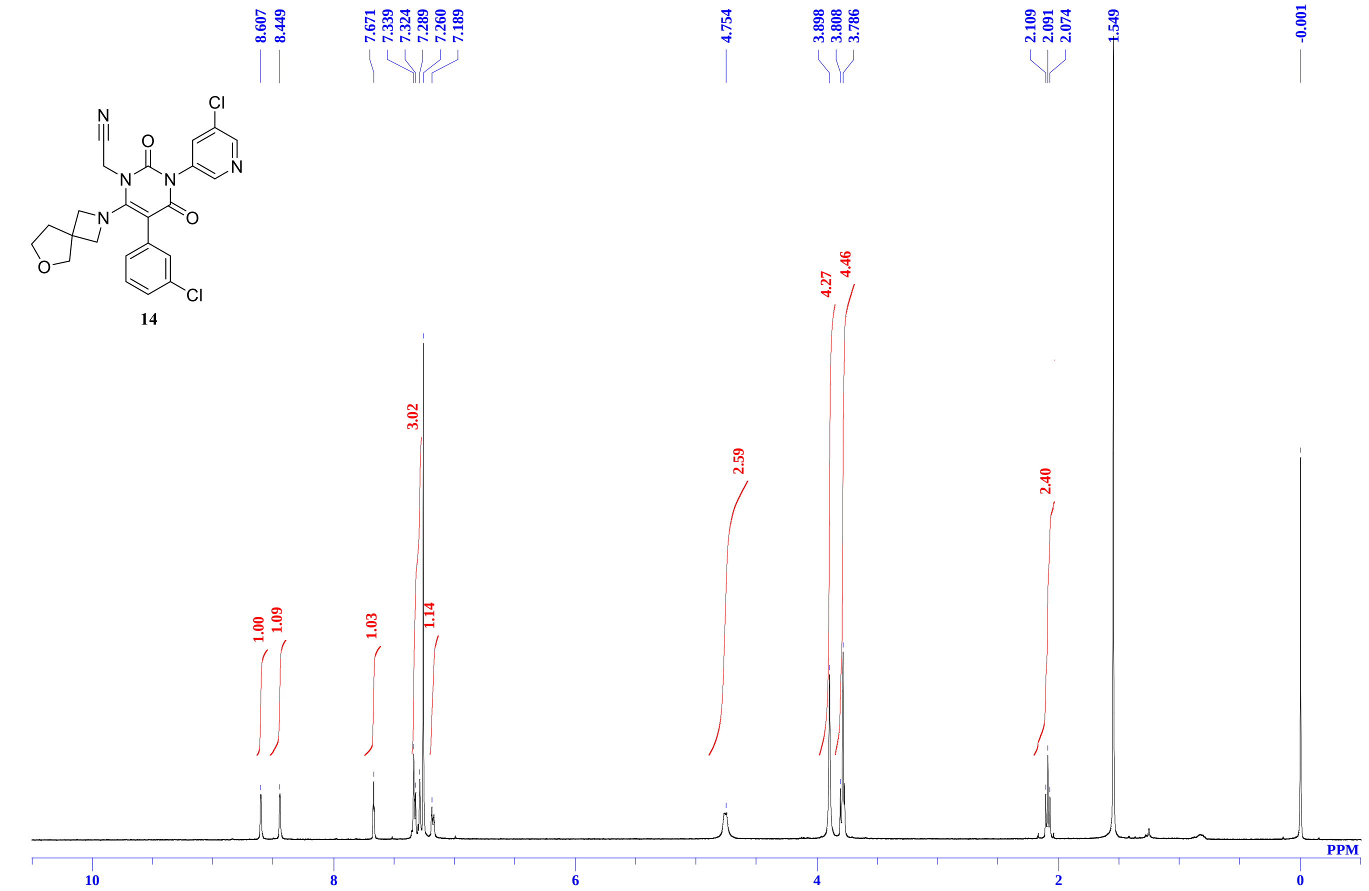


$^{13}C$ NMR spectra of **14** in $CDCl_3$

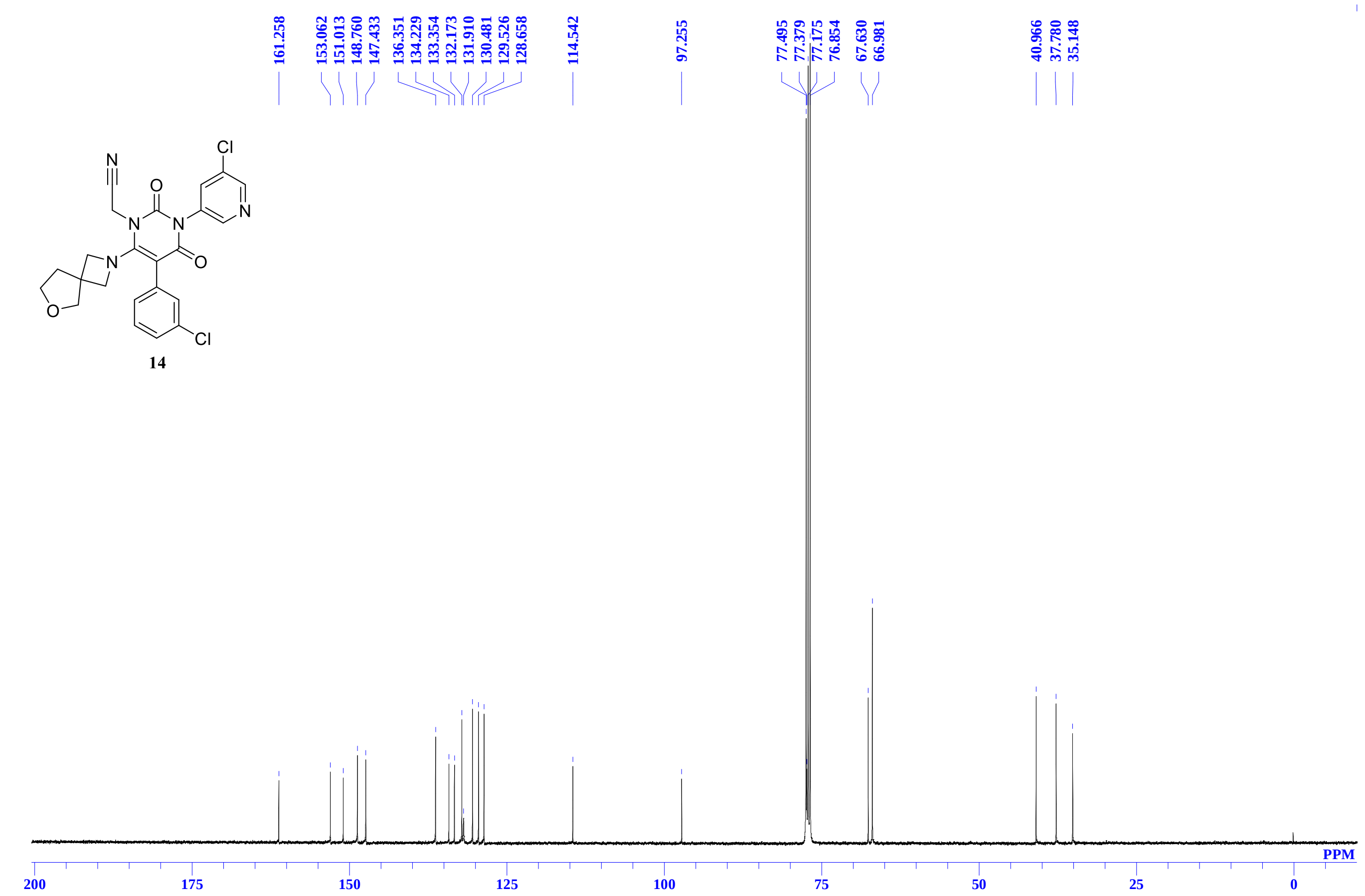

[1]H NMR spectra of **15** in DMSO-$d_6$

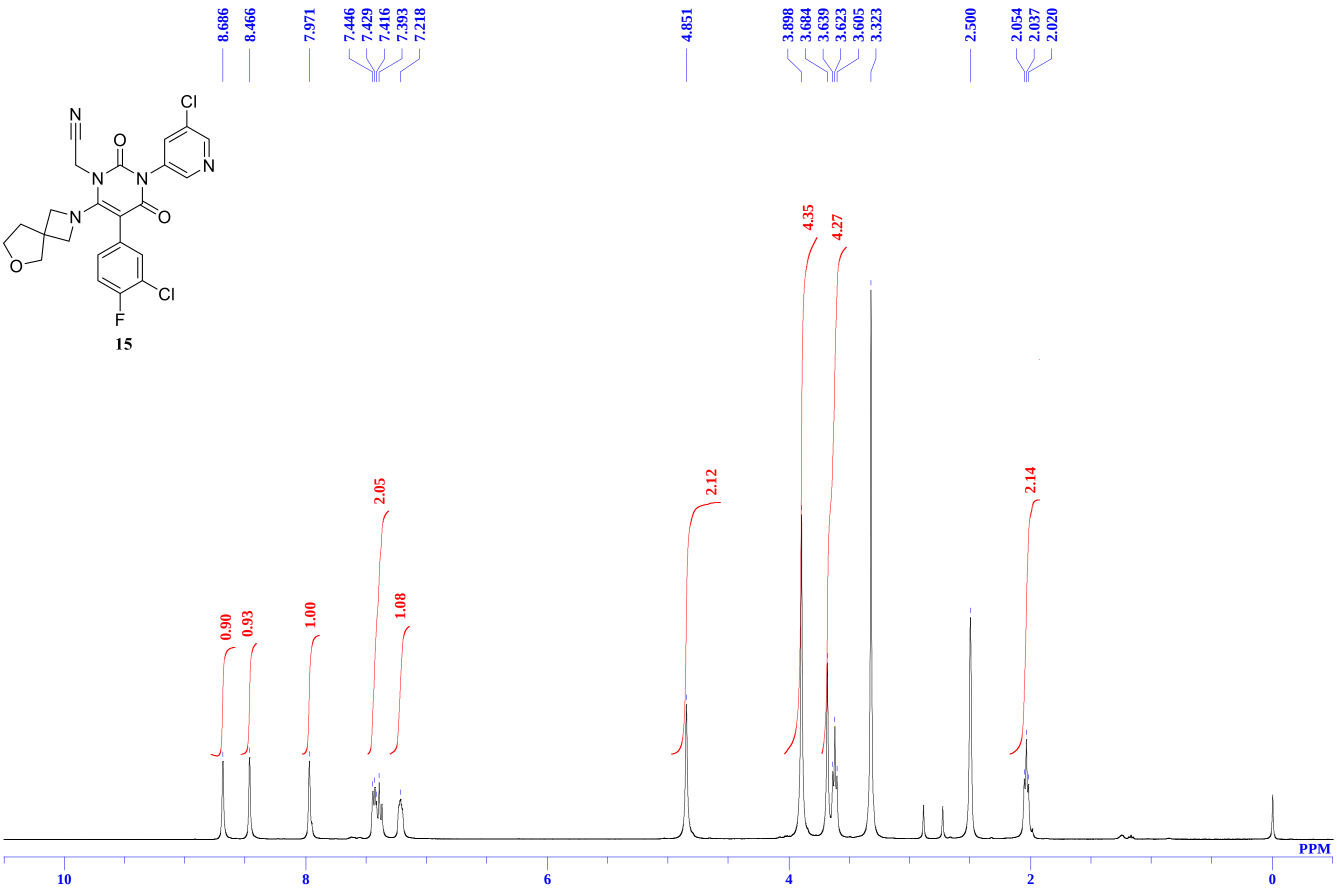


**[13]C** NMR spectra of **15** in DMSO-$d_6$

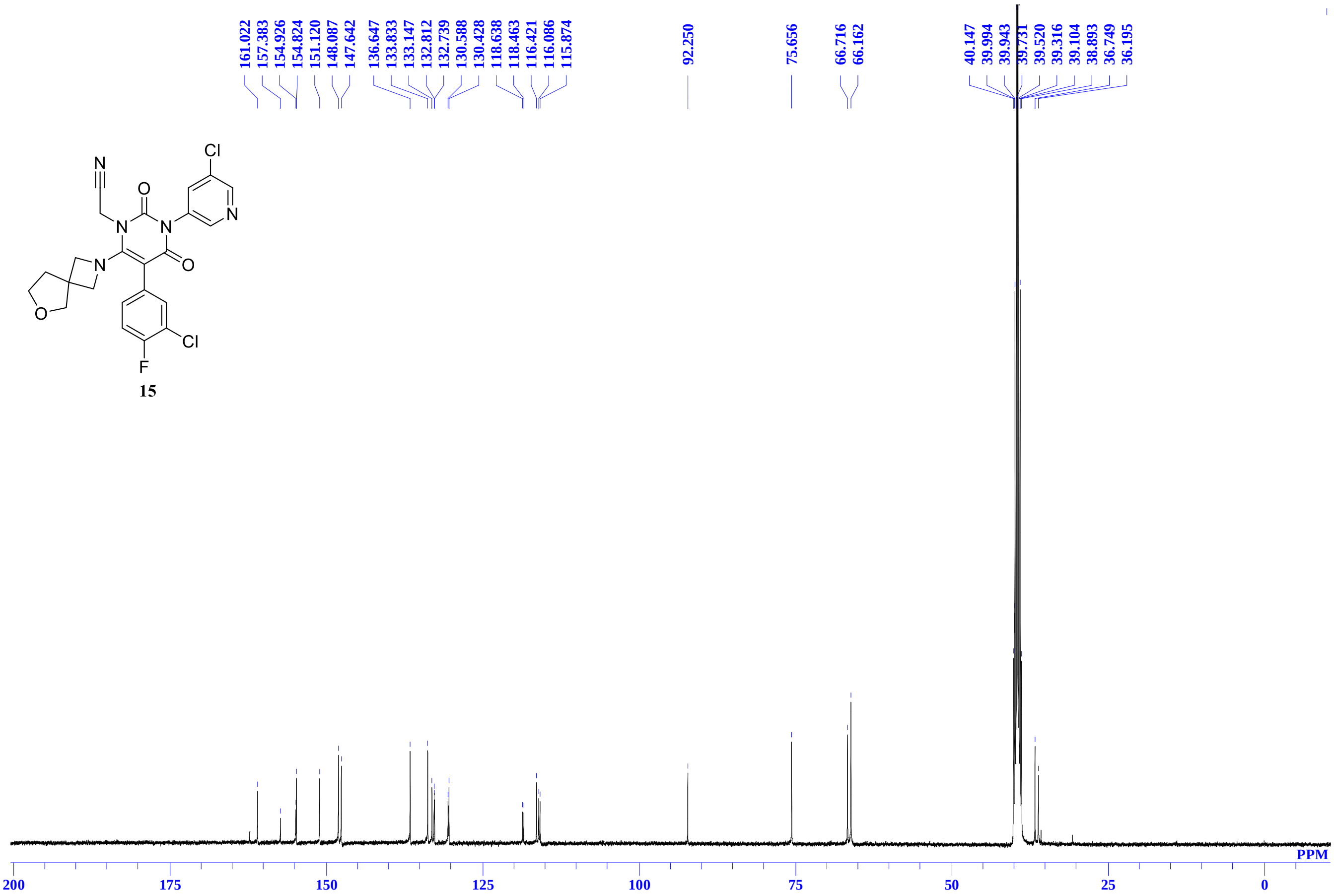

### $^{1}$H NMR spectra of **16** in DMSO-$d_6$

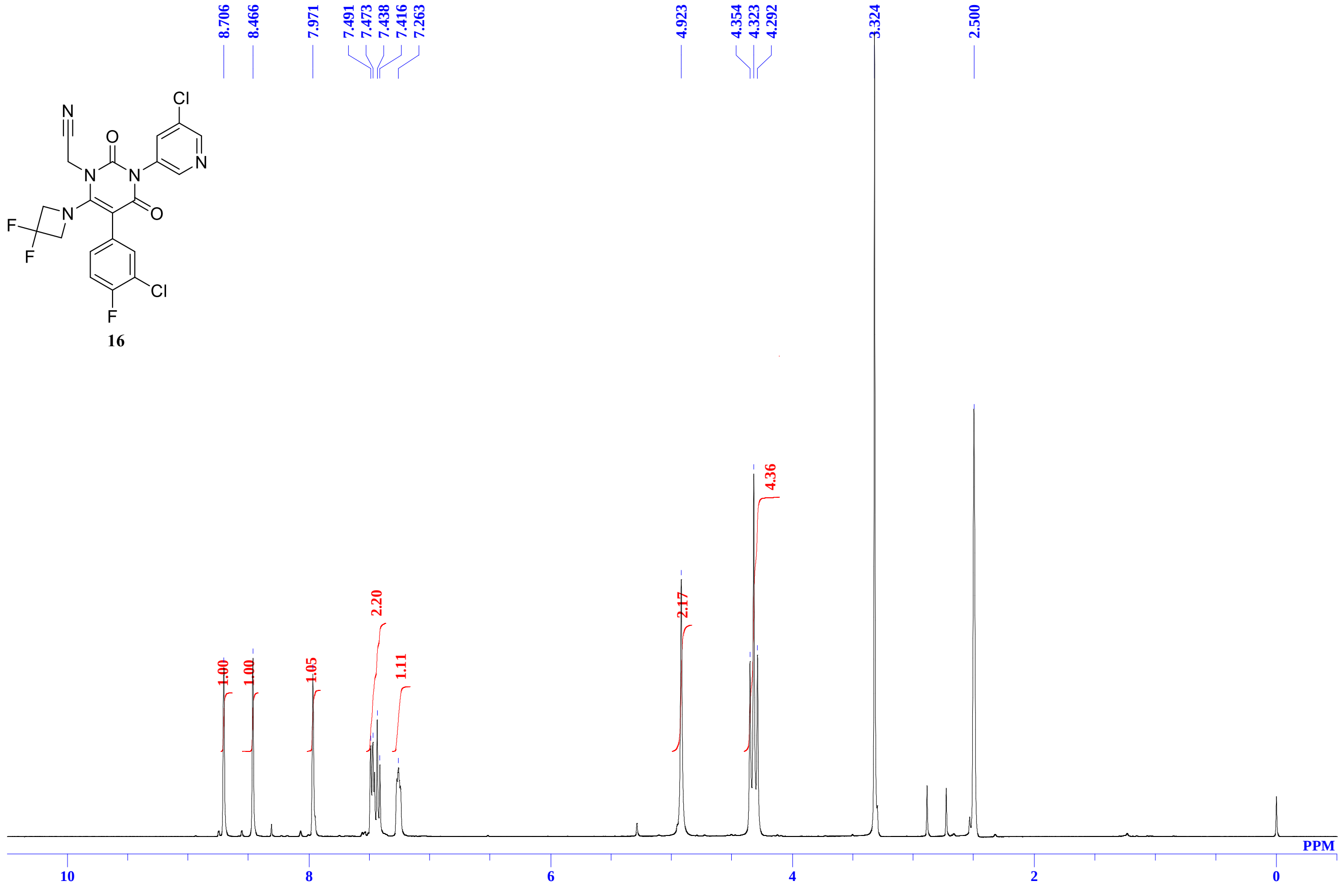


### $^{13}$C NMR spectra of **16** in DMSO-$d_6$

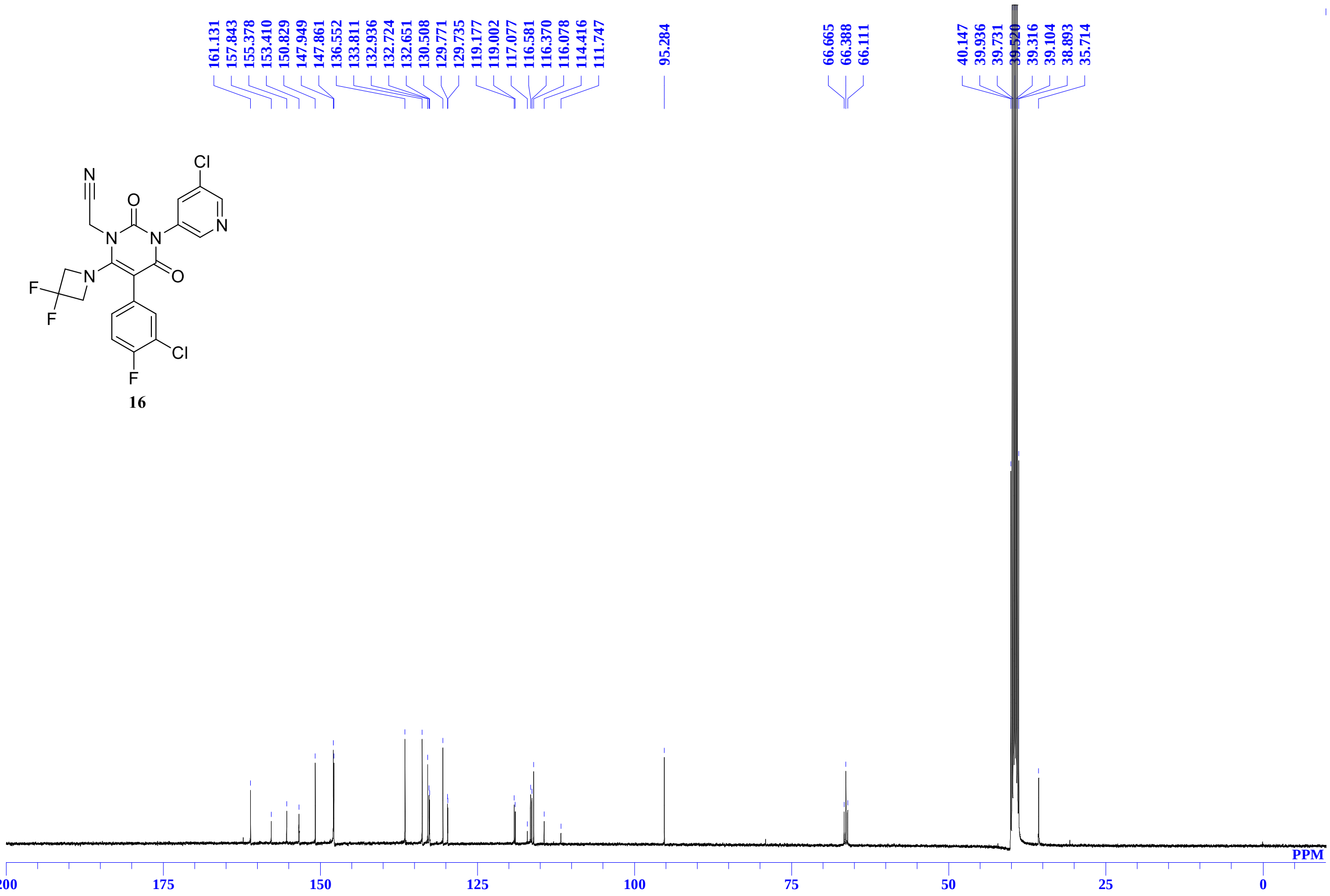

$^{1}$H NMR spectra of **17** in DMSO-$d_6$

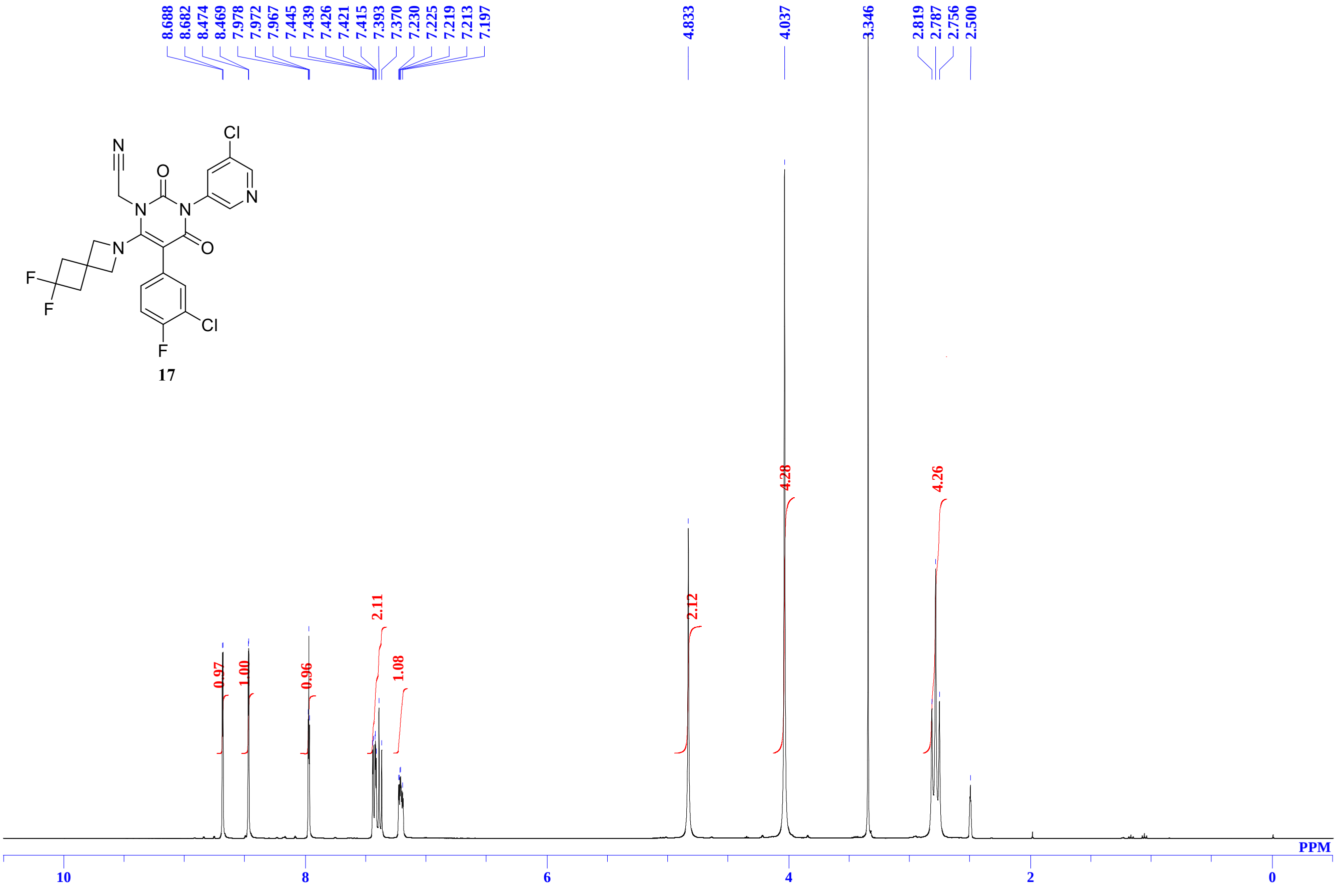


$^{13}$C NMR spectra of **17** in DMSO-$d_6$

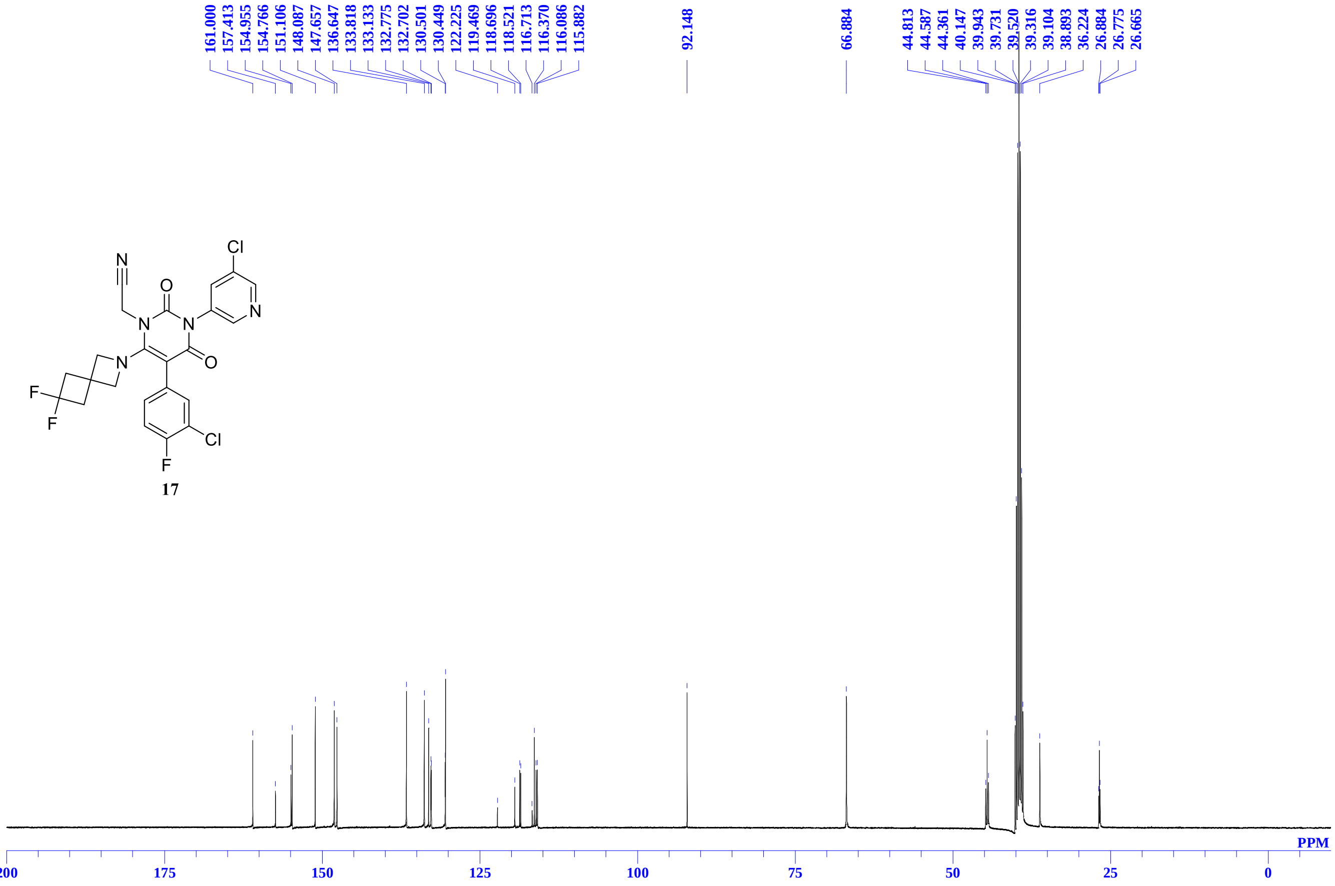

[1]H NMR spectra of **18** in DMSO-$d_6$

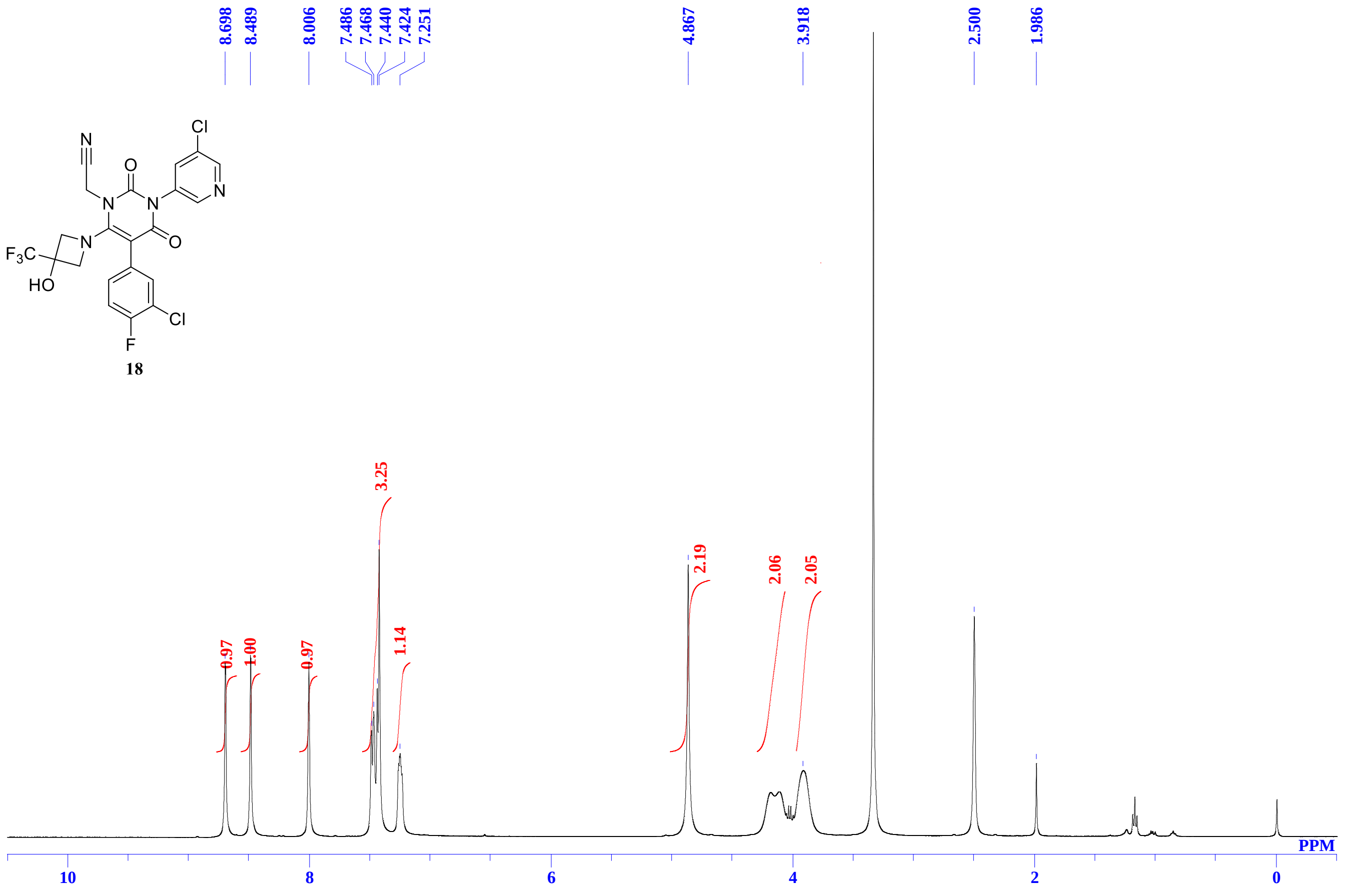


[13]C NMR spectra of **18** in DMSO-$d_6$

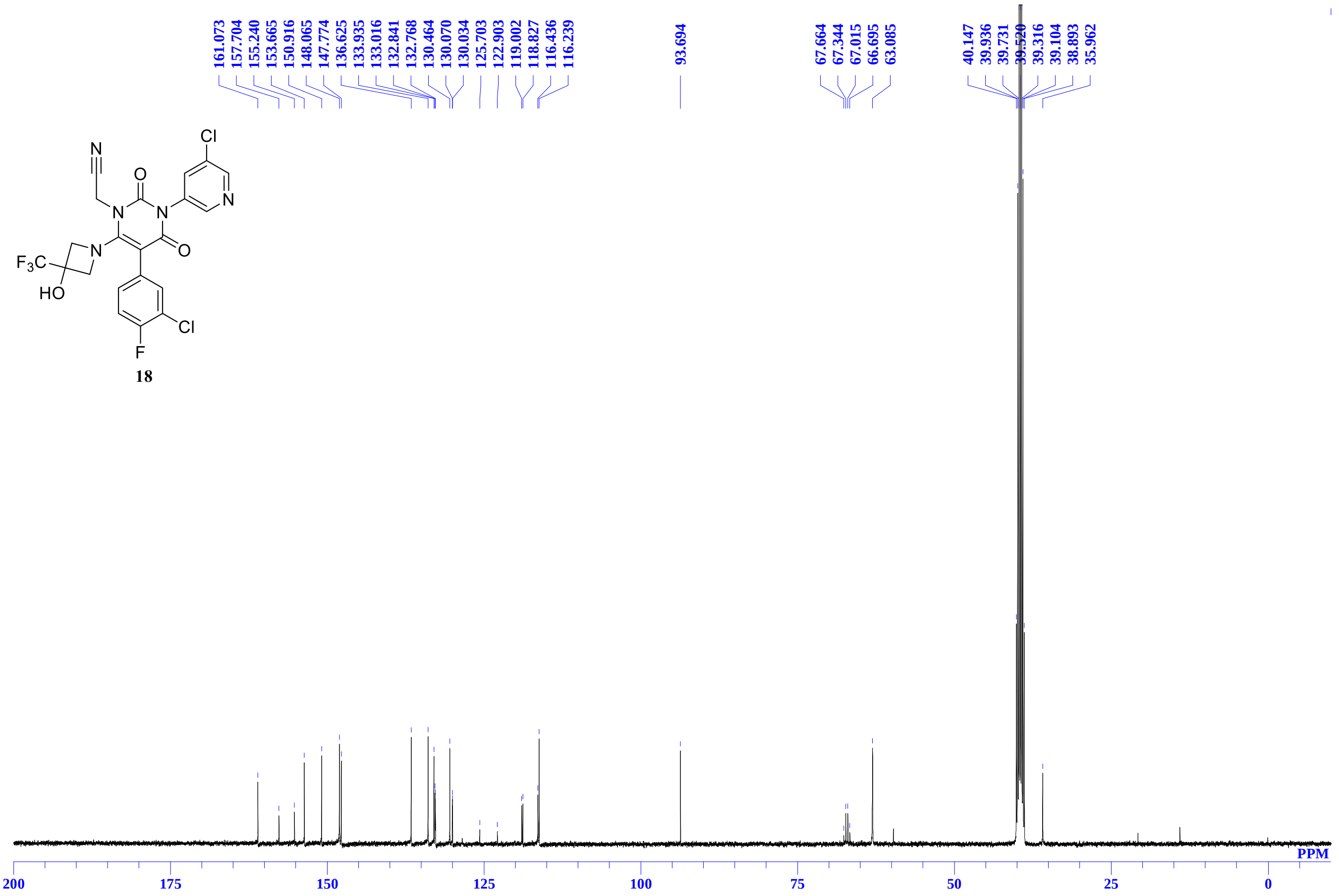

**HPLC chromatogram of compound 17 (S-892216)**

Analytical liquid chromatography of compound **17** (**S-892216**) was performed on a YMC-UltraHT Pro C18 Column (2.0 µm, 2.0 mm I.D.x100 mm, gradient from 10% to 90% B [A = 10 mmol/L $HCOONH_4$ / $H_2O$, B = 10 mmol/L $HCOONH_4$ / MeCN], flow rate: 0.4 mL/min) using a Shimadzu Nexera system equipped with LC-30AD binary gradient module and SPD-20AV detector (detection at 255 nm).

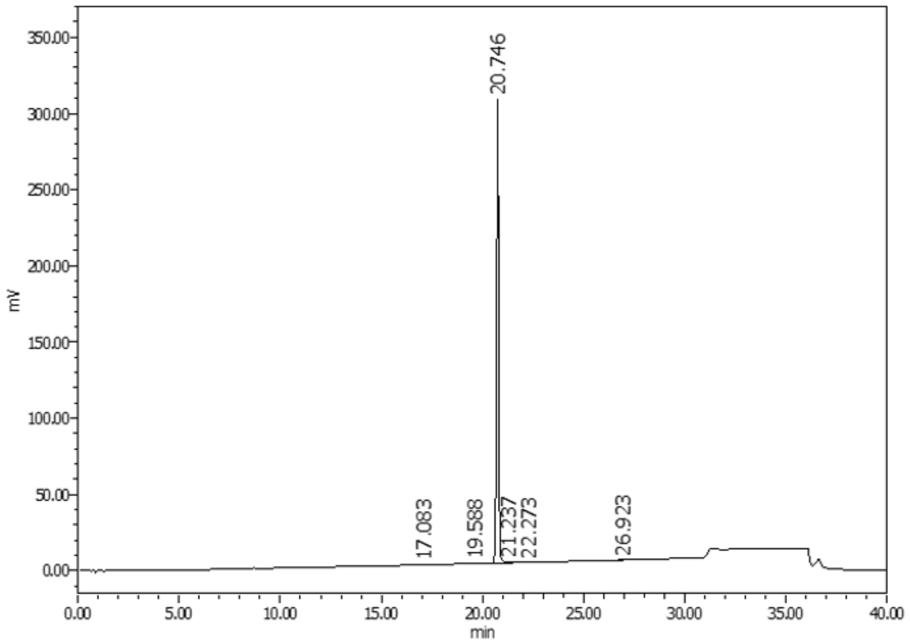


Time-program for gradient elution

| Time (min) | Mobile phase B (%) |
|---|---|
| 0 | 10 |
| 30 | 70 |
| 30.01 | 90 |
| 35 | 90 |
| 35.01 | 10 |
| 40 | 10 |

| | Retention time (min) | Peak area (µV·sec) | Peak height (µV) | % area |
|---|---|---|---|---|
| 1 | 17.1 | 1843 | 127 | 0.08 |
| 2 | 19.6 | 1616 | 79 | 0.07 |
| 3 | 20.7 | 2200909 | 304418 | 99.39 |
| 4 | 21.2 | 7832 | 820 | 0.35 |
| 5 | 22.3 | 959 | 54 | 0.04 |
| 6 | 26.9 | 1250 | 62 | 0.06 |

### HPLC chromatogram of compound 18

Analytical liquid chromatography of compound **18** was performed on a Unison C18 Column (3.0 µm, 4.6 mm I.D.x75 mm, gradient from 10% to 90% B [A = 0.1% THF in $H_2O$, B = MeCN], flow rate: 1.0 mL/min, Method : 0-15min [B]=10-90%, 15-20min [B]=90%) using a Shimadzu Prominence system equipped with LC-20AD binary gradient module and SPD-20AV detector (detection at 254 nm)

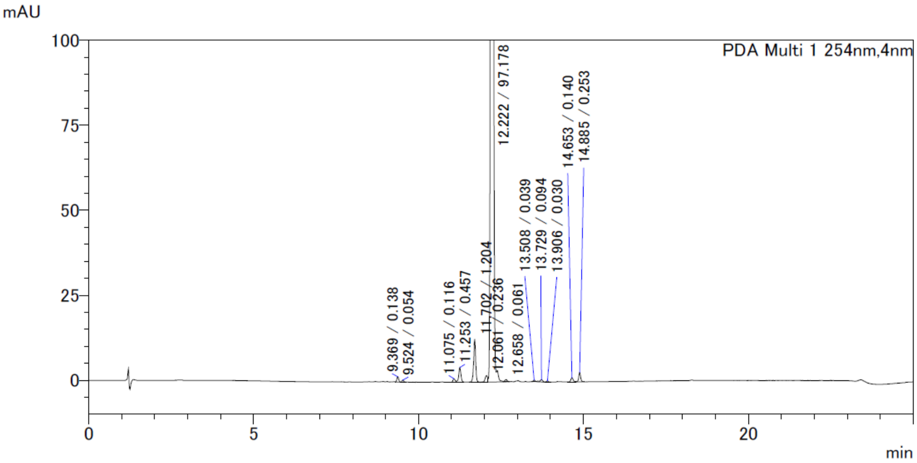


| | Retention time (min) | Peak area (µ·sec) | Peak height (µV) | % area |
|---|---|---|---|---|
| 1 | 9.369 | 6118 | 1610 | 0.138 |
| 2 | 9.524 | 2379 | 619 | 0.054 |
| 3 | 11.075 | 5167 | 1062 | 0.116 |
| 4 | 11.253 | 20259 | 4374 | 0.457 |
| 5 | 11.702 | 53425 | 12660 | 1.204 |
| 6 | 12.061 | 10481 | 1938 | 0.236 |
| 7 | 12.222 | 4310762 | 1100924 | 97.178 |
| 8 | 12.658 | 2702 | 568 | 0.061 |
| 9 | 13.508 | 1733 | 462 | 0.039 |
| 10 | 13.729 | 4191 | 674 | 0.094 |
| 11 | 13.906 | 1315 | 337 | 0.03 |
| 12 | 14.653 | 6196 | 1242 | 0.14 |
| 13 | 14.885 | 11223 | 2737 | 0.253 |

#### Physicochemical Properties of 17 (S-892216)

#### X-ray Powder Diffraction（XRPD）

A Crystal form of **S-892216** was analyzed by X-ray powder diffraction (XRPD) using SmartLab (Rigaku Corporation, Tokyo, Japan). The samples were placed in each hole (diameter = 2 mm, depth = 0.1 mm) on an aluminum plate, and their surfaces were smoothed by using a spatula. X-ray diffraction of the samples was performed with a 9 kW rotating anode using Cu Kα radiation (λ = 0.154186 nm) and a HyPix-3000 detector. The distance between the sample and the detector was 331 mm, and the diffractometer was equipped with a cross-beam optic. A parallel-slit collimator with 2.5° and a slit of 0.05 mm height and 0.5 mm width was used. The scan was performed from 3° to 32° (2θ) with β axis rotation (20 rpm) during data collection; the sampling step was 0.02° (2θ), and the count time was 40 s. The data were analyzed using Smart Lab Studio II X64 version 4.2.111.0 (Rigaku Corporation, Tokyo, Japan).

#### Thermogravimetry–Differential Thermal Analysis（TG-DTA）

Simultaneous measurements of the changes in weight and thermal profile as a function of temperature were performed by thermogravimetry–differential thermal analysis (TG–DTA) using a STA7200RV instrument (Hitachi High-Tech Science Corporation, Tokyo, Japan). The sample was placed into an aluminum pan and measured at 10 °C/min up to 350 °C. The results were analyzed using TA7000 standard analysis, version 11.2 (Hitachi High-Tech Science Corporation, Tokyo, Japan).

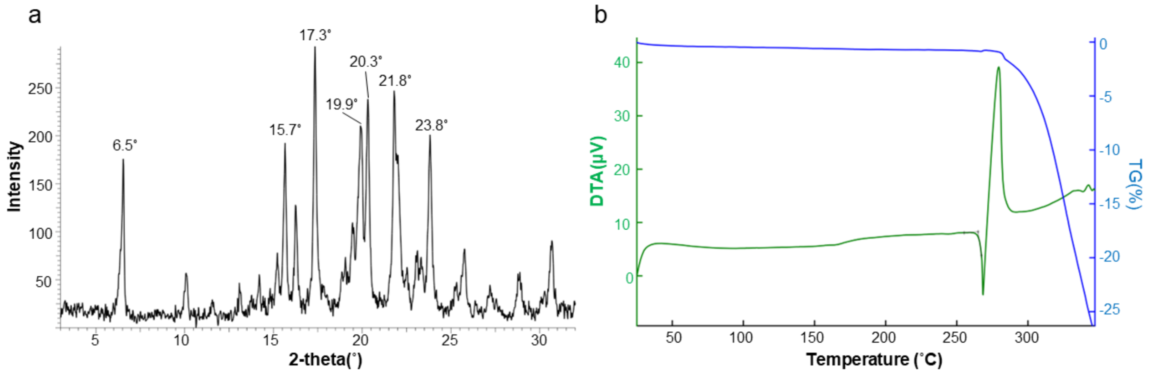


**Figure S2**. Physicochemical Properties of **S-892216**. (a) XRPD patterns. (b) TG-DTA.

**HRMS Spectra for compound 17 (S-892216)**

High-resolution mass spectrometry (HRMS) data were gathered on a Thermo Fisher Scientific Q Exactive Focus using Heatead-electrospray ionization using a Shimadzu Nexera system (LC-30AD, CBM-20A, SIL-30ACAM, CTO-20AC, DGU-20A$_{5R}$, FCV-20AH$_2$). Elution conditions: column, YMC-Triat C18 Column, 2.1 × 30 mm, 2.5 µm particle size; column temperature, 30°C; solvent A, water (0.1% formic acid); solvent B, acetonitrile (0.1%formic acid); gradient: 5−95% B in 3.5 min, 100% B in 0.50 min, 5% B in 1.0 min, 5.0 min total run time; flow rate, 0.5 mL/min. LC/MS conditions, ESI in positive mode; temperature, 380°C; ion spray voltage, 3500 V.

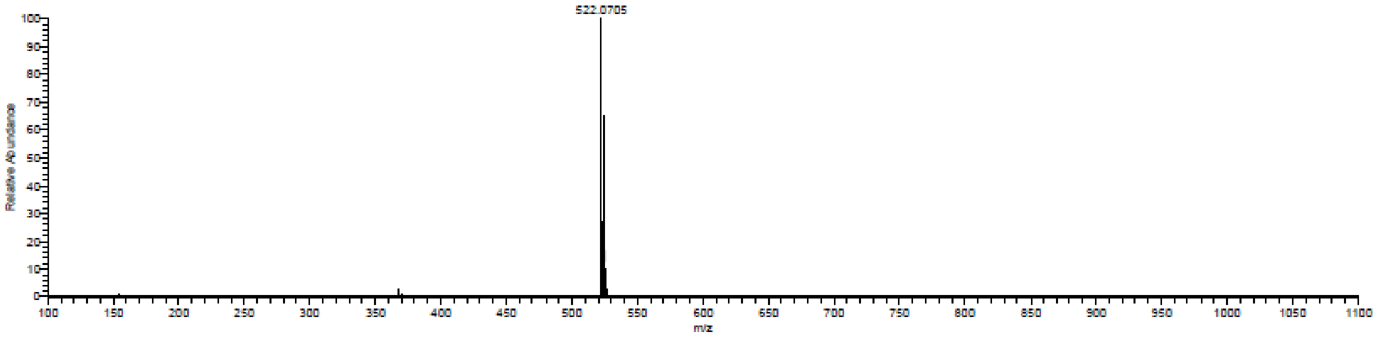


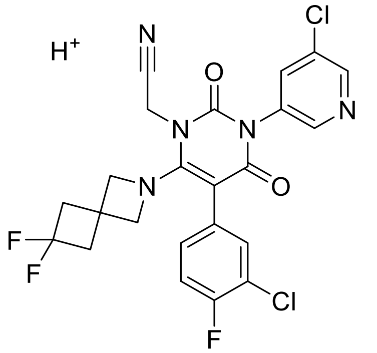

Chemical Formula: $C_{23}H_{17}Cl_2F_3N_5O_2^+$
Exact Mass: 522.0706

S2. Uehara, T.; Yotsuyanagi, H.; Ohmagari, N.; Doi, Y.; Yamato, M.; Imamura, T.; Sakaguchi, H.; Fukushi, A.; Takeda, Y.; Baba, K.; Nobori, H.; Miyamoto, T.; Arita, S.; Dodo, R.; Shimba, A.; Fukao, K.; Shishido, T.; Tsuge, Y.; Mukae, H. Ensitrelvir treatment-emergent amino acid substitutions in SARS-CoV-2 3CLpro detected in the SCORPIO-SR phase 3 trial, *Antiviral Research*, in press

S3. Kuroda, T.; Nobori, H.; Fukao, K.; Baba, K.; Matsumoto, K.; Yoshida, S.; Tanaka, Y.; Watari, R.; Oka, R.; Kasai, Y.; Inoue, K.; Kawashima, S.; Shimba, A.; Hayasaki-Kajiwara, Y.; Tanimura, M.; Zhang, Q.; Tachibana, Y.; Kato, T.; Shishido, T. Efficacy comparison of 3CL protease inhibitors ensitrelvir and nirmatrelvir against SARS-CoV-2 in vitro and in vivo. *J. Antimicrob. Chemother.* **2023**, 78 (4), 946-952, DOI: 10.1093/jac/dkad027

S4. Uemura, K.; Nobori, H.; Sato, A.; Toba, S.; Kusakabe, S.; Sasaki, M.; Tabata, K.; Matsuno, K.; Maeda, N.; Ito, S.; Tanaka, M.; Anraku, Y.; Kita, S.; Ishii, M.; Kanamitsu, K.; Orba, Y.; Matsuura, Y.; Hall, WW.; Sawa, H.; Kida, H.; Matsuda, A.; Maenaka, K. 2-thiouridine is a broad-spectrum antiviral nucleoside analogue against positive-strand RNA viruses. *Proc. Natl. Acad. Sci. USA.* **2023**, 120 (42), e2304139120, DOI: 10.1073/pnas.2304139120. Epub 2023

S5. Sahara, N.; Hirano, Y.; Thien, P. L.; Yamakawa, K.; Ohashi, E.; Shibuya, S.; Komano, K.; Hirai, K.; Masui, M.; Kato, G.; Higuchi, T.; Shimazaki, Y.; Kawajiri, T.; Oohara, T.; Tsuno, N.; Ohara, T. Development of a Manufacturing Process for S-892216 Part I: A Novel Method for Constructing a Multi-Substituted Barbiturate Skeleton for Scalable Synthesis. *ChemRxiv.* **2025**, DOI:10.26434/chemrxiv-2025-sdmw6 This content is a preprint and has not been peer-reviewed.

S6. Le, T. P.; Kawajiri, T.; Sahara, N.; Kato, G.; Hosoya, M.; Oohara, T.; Agura, K.; Ohara, T.; Goda, S. Development of a Manufacturing Process for S-892216 Part II: Improvements Toward Commercially Feasible Process. *ChemRxiv.* **2025**, DOI:10.26434/chemrxiv-2025-0wdzt This content is a preprint and has not been peer-reviewed.
